# Deep learning uncovers sequence-specific amplification bias in multi-template PCR

**DOI:** 10.1101/2024.09.20.614030

**Authors:** Andreas L. Gimpel, Bowen Fan, Dexiong Chen, Laetitia O. D. Wölfle, Max Horn, Laetitia Meng-Papaxanthos, Philipp L. Antkowiak, Wendelin J. Stark, Beat Christen, Karsten Borgwardt, Robert N. Grass

## Abstract

Multi-template polymerase chain reaction is a key step in many amplicon sequencing protocols enabling parallel amplification of diverse DNA molecules sharing common adapters in applications, ranging as wide as quantitative molecular biology and DNA data storage. However, this process results in a skewed amplicon abundance, due to sequence-specific amplification biases. In this study, one-dimensional convolutional neural networks (1D-CNNs) were trained on synthetic DNA pools to learn the PCR amplification efficiency of individual templates. These 1D-CNN models can predict poorly amplifying templates based solely on sequence information, achieving an AUROC/AUPRC of up to 0.88/0.44 with very imbalanced prevalence of 2%, thereby greatly outperforming baseline models relying only on GC content and nucleotide frequency as predictors. A new, general-purpose framework for interpreting deep learning models, termed `CluMo` provides mechanistic insights into the amplification biases. Most strikingly, specific amplification reactions were identified as suffering from adaptor-template self-priming a mechanism previously disregarded in PCR.

## Introduction

The technology to amplify DNA via polymerase chain reaction (PCR) is one of the key pillars of molecular diagnostics, enabling many qualitative and quantitative analytical techniques. While classically, PCR is used to amplify a single target sequence, the advent of massive parallel sequencing required simultaneous amplification of many short sequences (i.e., templates) sharing only short terminal adapters. Such multi-template PCR is now fundamental to many routine sequencing preparation workflows and is used across numerous fields, ranging from metabarcoding to DNA data storage^1–3^. However, the introduction of biases upon parallel amplification of sequence templates remains a key concern for multi-template PCR^4^, as any bias in amplification leads to skewed abundance data, thereby limiting method sensitivity and quantitative accuracy^5–11^.

The relevance of bias in multi-template PCR is highlighted by the ongoing efforts for its circumvention. In DNA- and RNA-sequencing, PCR bias has prompted the development of unique molecular identifiers^12–14^ and PCR-free workflows^7,15,16^ to mitigate or preclude biased abundance data in high-throughput sequencing. In DNA data storage, strategies for DNA immobilization^17,18^ and constrained coding^19,20^ have emerged to reduce the pool inhomogeneity and sequence dropout caused by deep replication of oligo pools via multi-template PCR. However, despite these efforts to mitigate PCR bias during multi-template PCR, a systematic understanding of its magnitude and mechanism - as well as tools for its investigation - are still missing.

While classical PCR with a single, well-defined target sequence is commonly optimized to ensure high amplification efficiency (e.g. via primer design and choice of annealing temperature),^21,22^ multi-template PCR faces another, separate problem: even a minor difference in amplification efficiency between templates leads to vastly different product-to-template ratios after exponential amplification^1,8,10^.

Commonly reported reasons for such differences in amplification efficiency include degenerate primers, amplicon length, template-product inhibition, amplicon GC content, polymerase choice, temperature profile, and stochastic effects^1,5,9,23–27^. However, manifestations of PCR bias are also commonly observed in the field of DNA data storage,^17,18,23,28,29^ where the properties of the template sequences are well-defined, and sequence information is often deliberately devoid of undesired properties (i.e. extreme GC content, long homopolymers, or secondary structure)^3,17–20^. The presence of skewed amplification rates under these optimal conditions therefore suggests the existence of additional factors contributing to sequence-specific PCR bias independent of most factors previously reported in studies on biological samples, and which is not readily identifiable via traditional statistical methods^23,27,29^.

Given the complexities of amplification biases, advanced computational and analytical techniques, particularly deep learning, offer promising avenues for deciphering the underlying patterns and mechanisms. Recent advancements in deep learning have revolutionized DNA sequence analysis, revealing complex characteristics and interactions previously obscured by traditional machine learning methods. For instance, Convolutional Neural Networks (CNNs) have significantly improved predictions of DNA-protein interactions^30^ and effects of non-coding variants^31,32^, while also enhancing our understanding of chromatin accessibility^33^. However, despite their predictive power and capability to handle large-scale datasets, the ‘black-box’ nature of deep learning models often limits their ability to elucidate the underlying molecular mechanisms^30,34^.

In contrast, traditional motif discovery methods such as MEME^35^ and Gibbs Sampling^36^, grounded in statistical principles, offer better interpretability through the identification of recurrent, short subsequences^35,37^. Later methods such as Weeder Web^38^ and DREME^39^, employing exhaustive search and differential analysis, further expanded the landscape of motif discovery. These statistical methods are often computationally intensive and may lack the discriminative power of deep learning. Therefore, a growing body of research seeks to bridge this gap by extracting interpretable motifs directly from ‘black-box’ models. DeepBind^30^ pioneered this approach by using CNNs for motif discovery, successfully identifying motifs correlated with DNA-protein binding sites by interpreting the convolutional filters in the first layer within the CNNs. Subsequently, DeepLIFT^40^ and SHAP^41^ further improved feature attribution analysis for general deep learning models, providing nucleotide-level attribution scores based on specific tasks. However, a key limitation of these methods lies in their focus on individual sequences, often neglecting a comprehensive overview of common motifs across the entire dataset. Their utility is therefore limited for global model explanation and general motif discovery, which would be crucial to better understand effects across multiple sequences as required for the understanding of the multi-template PCR bias.

To overcome these limitations, we employed synthetic oligonucleotide pools to generate large, reliably annotated datasets of sequence-specific amplification efficiencies, and train deep learning models to identify poorly amplifying sequences. Moreover, we present a novel motif discovery method – termed Motif Discovery via Attribution and Clustering (`CluMo`) – to identify specific sequence motifs linked to poor PCR efficiency and quantify their population-level importance. Aided by the model interpretability of `CluMo`, we attain a mechanistic understanding of the template-dependent PCR inhibition that causes PCR-induced and workflow-dependent bias in multi-template PCR.

## Results

### PCR amplification progressively skews coverage distributions

To systematically investigate the inherent bias in multi-template PCR, we first experimentally analyzed the PCR efficiency of individual, synthetic DNA sequences in multi-template PCR reactions. For this, the change in amplicon coverage was tracked for 12,000 random sequences with common, terminal primer binding sites (i.e., truncated Truseq adapters) over 90 PCR cycles using a serial amplification protocol (see Fig. 1). Specifically, six consecutive PCR reactions with 15 cycles each were performed, yielding a sample ready for sequencing at each iteration to quantify precise amplicon composition along the multi-template PCR amplification trajectory (see Methods).

**Fig. 1.**
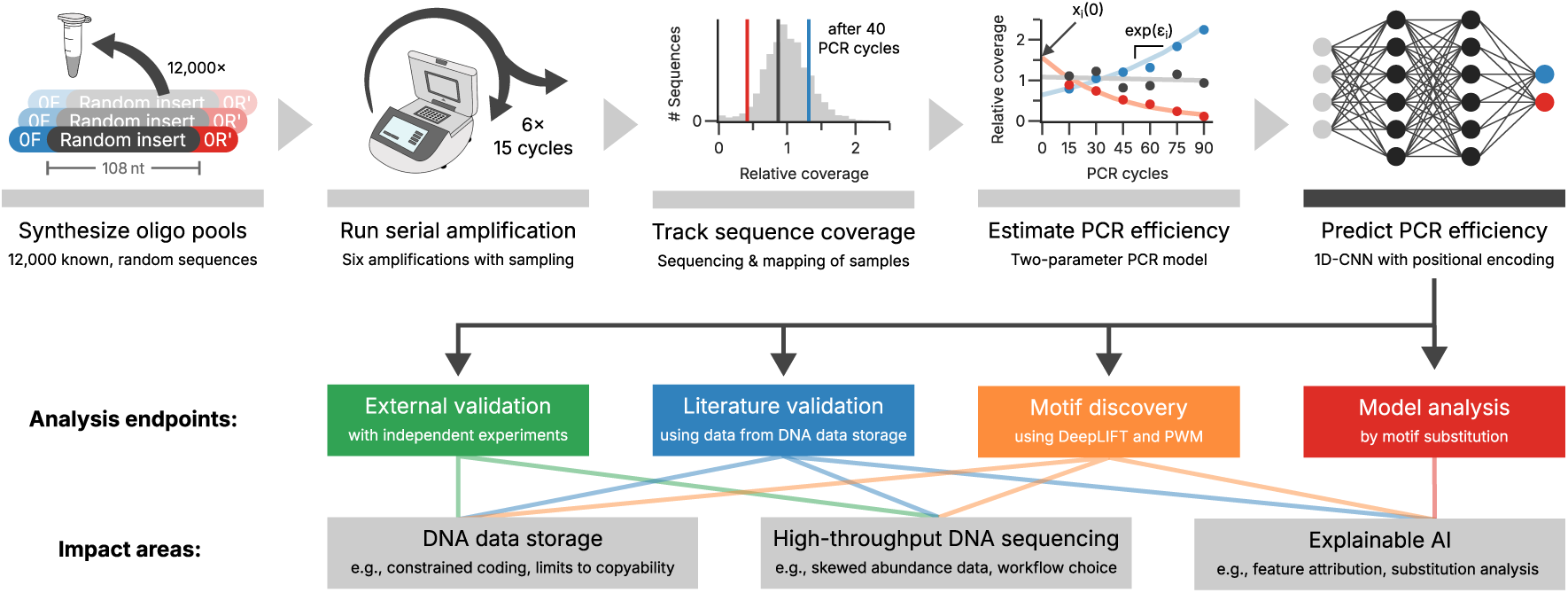
Overview of the workflow and the analysis endpoints. The workflow starts with the synthesis of randomized oligonucleotide pools with constant adapters (0F, blue and 0R’, red), which are consequently amplified serially to generate six samples with differing numbers of PCR cycles (from 15 to 90 cycles). After sequencing, the evolution of each sequence’s coverage as a function of cycle number is used to estimate the PCR efficiency in the two-parameter PCR model (see Methods). These estimates of the PCR efficiencies are used in the training of an 1D-CNN model for the binary classification of PCR efficiency.

During serial amplification, a progressive broadening of the coverage distribution was observed (see Fig. 2a), as previously reported^19,28^. Whereas the overall coverage distribution only changed marginally, a considerable number of amplicon sequences were either severely depleted or even no longer present in the sequencing data (see Fig. 2b). Because the presence of a GC-bias in amplification and sequencing is known^9,23,24,42^, the experimental analysis was performed on a second synthetic oligonucleotide pool in which the random sequences were constrained to 50% GC content (called GCfix). However, the progressive skew of the coverage distribution with increased PCR cycles and the increased fraction of sequences with low coverages was comparable between the GCall and GCfix pools (see Supplementary Fig. 17), suggesting the observed PCR bias is not caused by the sequences’ GC content.

**Fig. 2.**
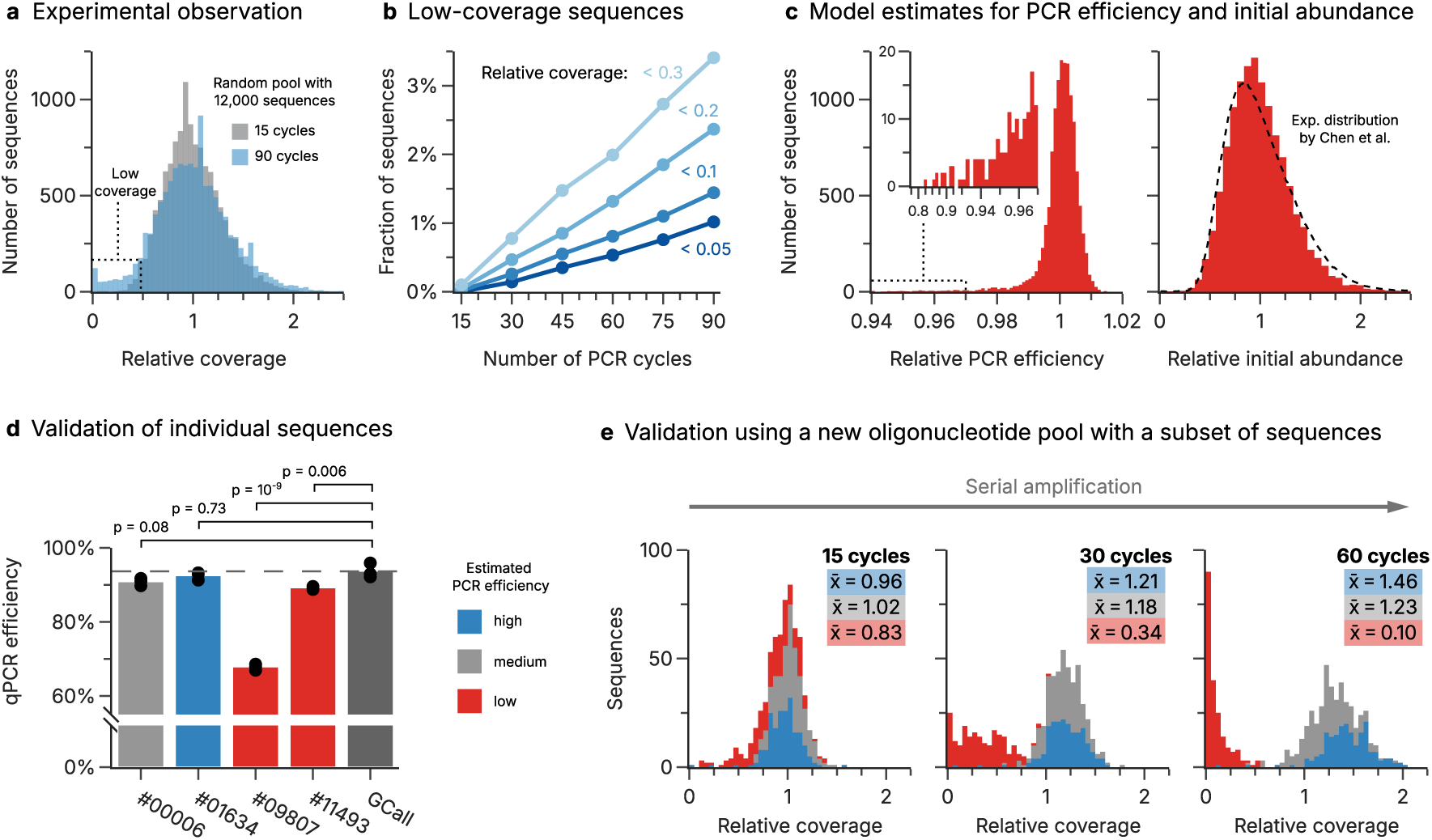
Estimation of sequence-level PCR efficiency and its experimental validation. **(a)** Observed normalized coverage distributions of the GCall pool after the first (15 cycles, grey) and the sixth round of serial amplification (90 cycles, blue). **(b)** Observed fractions of underrepresented sequences in the GCall pool over the course of serial amplification. Low-coverage sequences are further grouped by their relative coverage, from occurring less frequently than 30% (light blue) to lower than 5% (dark blue). **(c)** Distributions of the relative PCR efficiency (left) and relative initial abundance (right) estimated from the experimental data using a two-parameter fit to exponential PCR amplification (see Methods). The inset for the PCR efficiency shows the subset of sequences with very low amplification efficiency. The dotted line superimposed onto the distribution of relative initial abundance shows the experimentally-determined distribution by Chen et al.^29^, using a ready-to-sequence pool. **(d)** qPCR efficiencies of four individually synthesized, and arbitrarily selected sequences from the GCall pool (#00006 through #11493), and of the GCall pool itself, as measured with qPCR dilution curves. Two of the individual sequences had shown a low amplification efficiency during the serial amplification (#09807 and #11493, red bars). Samples #00006 and #01634 had shown average (grey) and good amplification performance (blue) respectively. Amplification efficiencies were significantly different by one-way ANOVA (*N* = 3 per sample, *F* (4, 10) = 252, *p* = 5 × 10^-10^), and the results of a post-hoc Tukey’s range test are shown above the bars (see Supplementary Tables 2 and 3). **(e)** Observed normalized coverage distributions after 15, 30, and 60 cycles of serial amplification (iterations 1, 2, and 4 respectively) of a new pool containing a subset of the sequences present in the GCall and GCfix pools. Sequences were again selected by their estimated PCR efficiencies, and grouped by a high (blue), medium (gray), or low (red) PCR efficiency (see Methods). Insets show the mean coverage across all sequences in each category for that experiment.

### Sequence-specific amplification efficiencies for multi-template PCR

In order to translate the observed changes in sequencing coverage to a quantifiable PCR bias, a simple fit of the sequencing data to an exponential PCR amplification process^5,6^ was performed (see Methods). This fit uses two parameters per amplicon sequence: the initial bias caused by uneven coverage after synthesis^29^, and the PCR-induced bias caused by each sequence’s individual amplification efficiency (*ɛ_i_*, see Supplementary Fig. 1 for an illustration). The obtained estimates for the initial coverage bias were comparable to experimental data using PCR-free sequencing of oligonucleotide pools^29^ (see Fig. 2c, dashed line) and the distributions of amplification efficiencies were comparable across both datasets (see Supplementary Fig. 17). The data revealed a small subset of sequences (representing around 2% of the pool) with very poor amplification efficiency in both datasets (see Fig. 2c, inset). With estimated efficiencies as low as 80% relative to the population mean (equivalent to a halving in relative abundance every 3 cycles), these sequences were often no longer present in the sequencing data after 60 cycles.

### Poor amplification is reproducible and independent of pool diversity

Two orthogonal experiments were conducted to verify that the sequencing-based quantification of the PCR efficiency is reproducible. For this, the sequences from the two pools (GCall and GCfix, 12,000 sequences each) were categorized as either average, good, or poorly amplifying. In the first experiment, four such sequences were arbitrarily selected and their efficiencies were evaluated using dilution curves in single-template qPCR (see Methods). Indeed, the sequences which were identified as poorly amplifying from the sequencing data also had significantly lower amplification efficiencies in qPCR, as shown in Fig. 2d. For the second experiment, a new oligo pool was synthesized, comprising 1000 of the categorized sequences from the original GCall and GCfix experiments (see Methods). Fig. 2e shows the evolution of their sequence coverage with increasing PCR cycles during serial amplification, stratified by the sequences’ assigned category. While the difference between average and well amplifying sequences is less evident, virtually all sequences previously categorized as poorly amplifying were drastically under-represented even after just 30 PCR cycles, and effectively drowned out completely by cycle number 60 (see Supplementary Fig. 20 for full data). These results show that there are specific amplicon sequences which significantly and reproducibly amplify less efficiently than the remaining sequences in the pool, and this bias does not depend on the composition of the pool, but rather on the sequence of individual oligos.

### Positional sequence information is critical to predict poor amplification with deep learning

In order to understand why the amplification of a fraction of sequences in multi-template PCR is hampered, we focused on the worst-performing 2% of sequences from the GCfix and GCall experiments (see Fig. 3a and Methods, other thresholds in Supplementary Figs. 22-24). Motivated by literature on GC-induced PCR bias,^23,24,27^ we first attempted to explain the data using a Lasso regularized logistic regression (LR) model, with GC content and base frequencies as features. However, the regression model performance was poor (see Fig. 3b+c), with a prediction accuracy close to a random classifier. This indicates that poor amplification of some sequences cannot be explained by the base composition or GC content alone, which is also confirmed by the similarity between the datasets with variable and fixed GC content (GCall and GCfix, see Supplementary Fig. 17). In order to improve upon the regression model, three more models based on established deep-learning architectures were trained on the datasets: a RNN, a 1D-CNN, and a 1D-CNN with positional encoding. While all deep-learning models outperformed the regression model considerably (see Supplementary Fig. 21), the 1D-CNN with positional encoding - outlined in Fig. 3a - was selected for further investigation due to its superior predictive power. When evaluated within our datasets (GCall and GCfix), the 1D-CNN models with positional encoding show an average AUROC (Area Under the Receiver Operating Characteristic curve) of 0.88 and 0.87, and an average AUPRC (Area Under the Precision-Recall Curve) of 0.42 and 0.44, respectively (see Fig. 3b) using 5-fold cross-validation, thereby improving upon the regression baseline by a factor of *>*10x. The evaluation across the two datasets shows the generalizability of the models, achieving largely identical performance when evaluated on each other’s data (see Fig. 3c). Importantly, the performance of the 1D-CNN was greatly affected by the introduction of positional encoding, indicating that the PCR bias is mainly attributable to position-specific features within the template sequence.

**Fig. 3.**
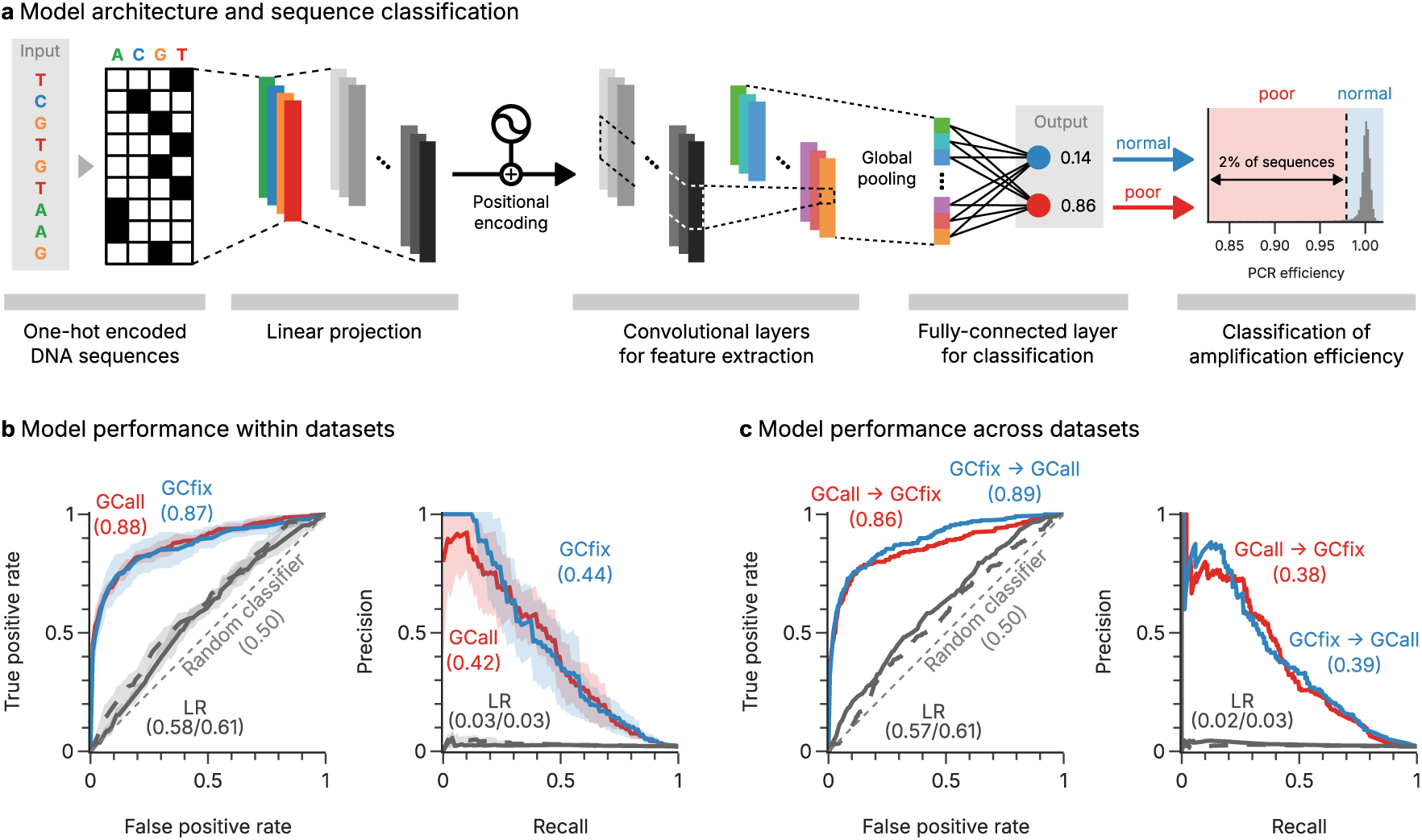
Classification of DNA sequences by estimated amplification efficiency. **(a)** A depiction of the 1D-CNN model with positional encoding to classify sequences’ amplification efficiency based on their structural attributes. Batch normalization and Rectified Linear Unit (ReLU) activation are used in-between the convolutional layers. **(b)** Evaluation of the models’ performance within the GCall (red) and GCfix (blue) datasets, presenting AUROC (left) and AUPRC (right) scores, with the shaded area showing the uncertainty in the results from the five-fold cross-validation. The performance of the baseline, the logistic regression (LR) model, on the datasets is also shown (gray, GCall solid, GCfix dashed). **(c)** Evaluation of the model’s performance across datasets, from GCall to GCfix (red) and GCfix to GCall (blue), presenting AUROC (left) and AUPRC (right) scores. The performances of the baseline LR model on the datasets is also shown (gray, GCall to GCfix solid, GCfix to GCall dashed).

### Discovery and validation of positional motifs associated with low PCR efficiency

The analysis above shows that 1D-CNN models perform well in identifying sequences with poor amplification efficiency from their sequence information alone. However, the black-box nature of the 1D-CNN models does not allow an interpretation of the sequence features responsible for this performance. To remedy this, we developed a novel motif discovery approach called `CluMo` to elucidate sub-sequence features, based on DeepLIFT^40^, which was extended using k-mer analysis and clustering to interpret the trained model (see Fig. 4a and Methods). Applying this analysis to the GCall and GCfix datasets identified several positional motifs strongly associated with poor amplification efficiency (see Fig. 4b). Interestingly, most identified motifs included a common *CGTG* subsequence, usually flanked by similar nucleotides with lower attribution scores. Further analysis reveals that these motifs exhibit a marked propensity to occur at the beginning of poorly amplified sequences (i.e. adjacent to the primer binding site), a trend that is not observed in sequences with normal amplification efficiency (see Fig. 4c).

**Fig. 4.**
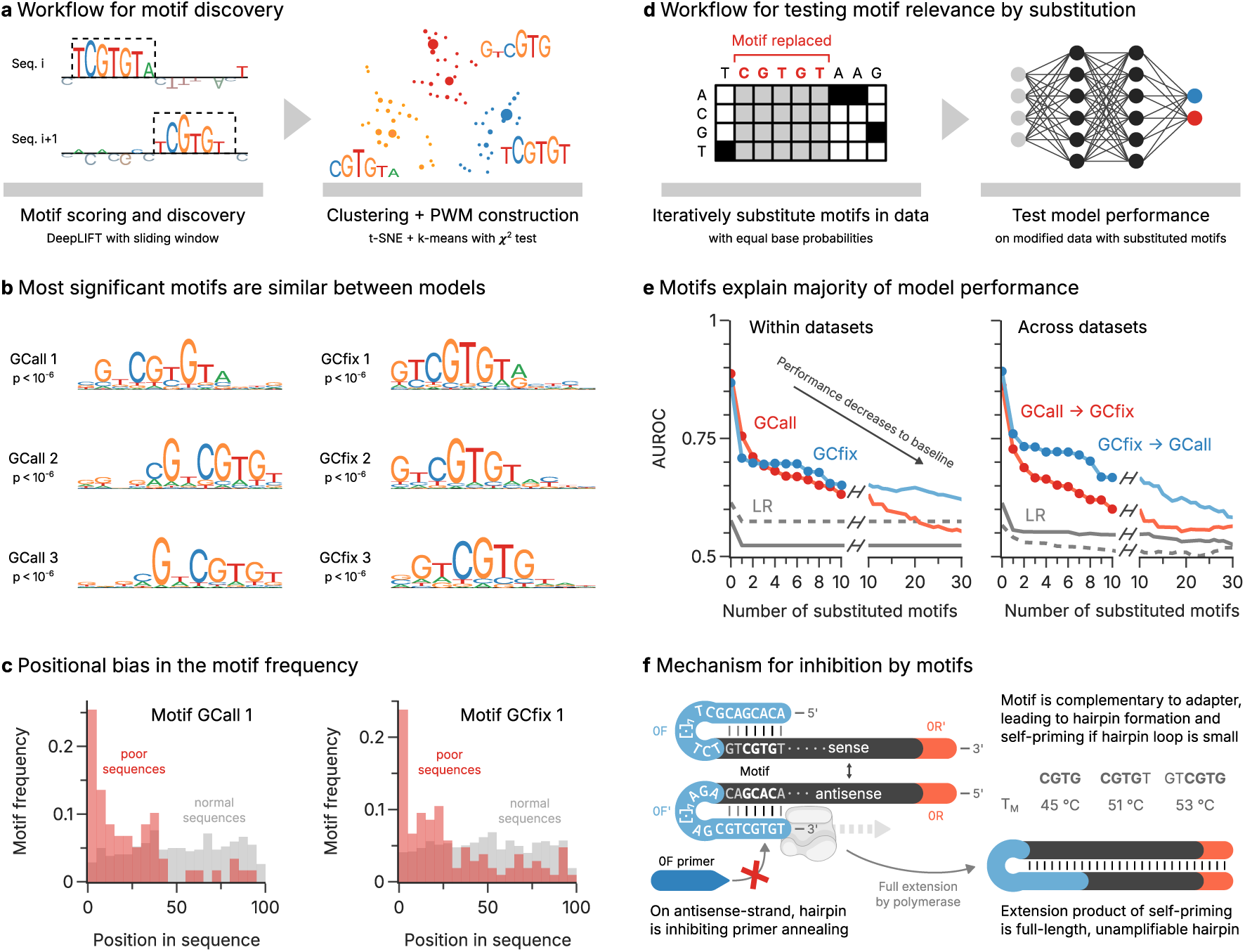
Discovering motifs affecting amplification efficiency and testing their relevance. **(a)** Workflow for discovering motifs with `CluMo`, based on motif extraction with DeepLIFT^40^, t-distributed stochastic neighbor embedding (t-SNE), and k-means clustering. The significance of the resulting position weight matrices (PWM) are assessed with a chi-squared test. **(b)** Most significant motifs identified for the GCall (left column) and GCfix datasets (right). The significance of the chi-squared test of each PWM is also shown. **(c)** Positional bias in the occurrence of the two motifs GCall 1 and GCfix 1 (see Panel b). The motifs in the poorly amplifying sequences (red) are more frequent at the beginning of the sequence, whereas there is no bias in normal sequences (gray). More data is shown in Supplementary Fig. 11. **(d)** Workflow for validation the discovered motifs in the trained model by iteratively replacing the motifs in the test data (ordered by p-values) without further retraining. The decrease in predictive power of the model upon motif replacement is correlated to that motif’s relevance to the model output. **(e)** Evolution of model AUROC as a function of the number of substituted motifs in the test data, either using internal validation (left) or testing across datasets (right). The models trained on GCall (red) and GCfix (blue) approach the performance of the baseline LR model (gray; GCall solid, GCfix dashed) as the number of substituted motifs increases. An evaluation using AUPRC as model performance metric is shown in Supplementary Fig. 4. **(f)** Hypothesized inhibition mechanism explaining the amplification disadvantage conveyed by the CGTG (sub-)motif. The motif is complementary to the 5’-adapter present on all oligos, thereby enabling hairpin formation and selfpriming. This inhibits primer annealing and leads to the formation of full-length hairpins which cannot be amplified further. An identical mechanism also enables hairpin formation at the 3-adapter, see Supplementary Fig. 16. Melting temperatures of the hairpins involving different motifs were calculated with mfold^43^.

In order to assess the importance of motifs for model performance, we iteratively replaced motifs, ranked by their p-values, in the test data with elements having equal base probabilities (see Fig. 4d). Evaluating the models’ classification performance after the substitution of each motif highlighted a pronounced decline toward baseline performance, as quantified by a considerable drop in AUROC (see Fig. 4d+e). This degradation highlights the critical role these motifs play in the predictive power of the model.

### Adapter-template self-priming impedes template amplification

The interpretability of the trained 1D-CNNs afforded by `CluMo` provided two important insights into the PCR-induced bias in the GCall and GCfix datasets: the role of *CGTG*-based sequence motifs, and their proximity to adapter sequences. Motivated by the latter, we looked for potential interactions between the terminal adapter and the uncovered sequence motifs. Indeed, all motifs identified by `CluMo` share short complementarities to the 5’ end of the adapter which support the formation of hairpins (see Fig. 4f). Importantly, a similar hairpin can also form with the complementary template strand, involving the 3’ end of the adapter. This hairpin structure inhibits primer annealing and enables self-priming, both of which will decrease amplification efficiency.

To investigate the plausibility of the hypothesized amplicon inhibition through self-priming, we first considered thermodynamic stability of the proposed hairpin during the PCR’s annealing step (i.e., at 54 *^◦^*C with 50 mM Na^+^) by hybridization prediction^43^. Despite the small size of its stem, hairpin formation was competitive to primer annealing, with predicted melting temperatures of 45-53 *^◦^*C (see Fig. 4f and Supplementary Fig. 14c). In line with literature data^44,45^, extending the hairpin loop by moving the motif downstream decreased its thermodynamic stability, and thereby diminished its expected impact on amplification efficiency (see Supplementary Fig. 14a). This explains the observed positional enrichment of motifs towards the 5’-end in poorly amplifying sequences (see Fig. 4c). Consistent with a self-priming induced inhibition mechanism, specific motifs with complementarity to the 3’end of the opposite adapter were also highly enriched within poorly amplifying amplicon sequences (see Supplementary Fig. 15).

To further validate whether self-priming by motif-adapter hairpins affects amplification efficiency, we determined amplification rates for the four individual sequences presented in Fig 2d using degenerate primers and qPCR (see Methods). These primers extended the previously used primers with four degenerate A/T nucleotides at each 5’-end, in order to introduce complementary A/T tails into template sequences during amplification. Due to these tails’ inability to hybridize with the template - thereby impeding extension by polymerases - we expect self-priming to be suppressed and amplification efficiency to be restored to normal levels (see Supplementary Fig. 16b for illustration). Indeed, qPCR with degenerate primers showed full recovery of amplification efficiency for the previously poorly amplifying sequence with a motif (#09807, from 68 ± 2% to 81 ± 5%), whereas all other sequences’ efficiencies decreased slightly (see full data in Supplementary Fig. 7). These results strongly support the proposed mechanism of inhibited amplification by adapter-mediated self-priming, which was entirely inspired by the interpretation of the trained 1D-CNN with our motif discovery approach `CluMo`.

### Generalization and benchmarking to literature datasets

To assess the generalizability of our model and benchmark the existence of similar motif-dependent biases in other workflows, we investigated the performance of the 1D-CNN model on a range of literature datasets from the DNA data storage community^17–19,46,47^. Contrary to literature data from biological fields, all of these datasets include sequencing data of *deep copies* (i.e., after at least 100 PCR cycles), and use well-defined synthetic DNA sequences. However, these datasets differ in their experimental conditions (e.g., choice of polymerase and adapter, see Supplementary Table 7). The performance of the 1D-CNN models trained and evaluated across all datasets is shown in Fig. 5 (for AUPRC, see Supplementary Fig. 5). As expected, all models perform best internally (diagonal in Fig. 5), but performance on other datasets is generally poor. Notable exceptions are two groups of datasets whose models exhibit some transferability between them: GCall, GCfix, and Koch et al.^46^ (mean AUROC/AUPRC: 0.71/0.17), as well as Erlich et al.^19^ and Gao et al.^17^ (0.87/0.17).

**Fig. 5.**
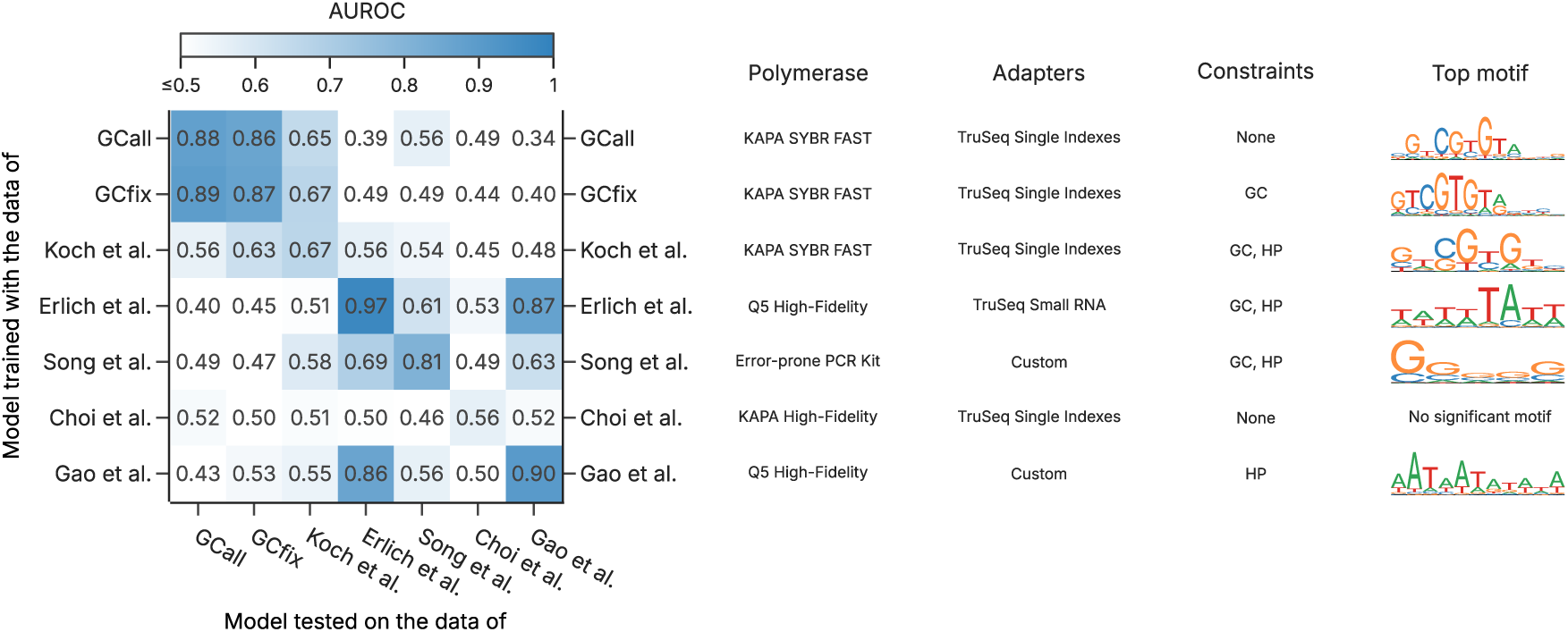
Assessing model performance across literature datasets. Heatmap of the area under the receiver-operating characteristic (AUROC) metric for the models trained and tested on the different literature datasets. Additional information on the choice of polymerase and amplification adapters is provided, together with constraints used during sequence design and the highest-ranked motif identified by `CluMo`. Additional information on the literature datasets is provided in Supplementary Table 7, and the corresponding heatmap using the area under the precision-recall curve (AUPRC) is given as Supplementary Fig. 5.

For both groups of datasets exhibiting model transferability, the use of common experimental conditions (i.e., polymerase and/or adapters, see Fig. 5 and Supplementary Table 7) suggests a workflow-dependent nature of PCR bias. Indeed, using `CluMo` to investigate relevant submotifs, the previously discussed *CGTG* motif also arises in the model of Koch et al.^46^, with a similarly strong positional preference towards the 5’-adapter (see Supplementary Fig. 11). The presence of this motif in the data of Koch et al.^46^ therefore underscores the predictive power of the models trained on GCall or GCfix, and illustrates the presence of the PCR bias hypothesized to stem from adapter-mediated self-priming in literature datasets.

In contrast, the transferability between models trained on Erlich et al.^19^ and Gao et al.^17^ likely stems from the Q5 polymerase used in both studies. Analysis by `CluMo` reveals a strong association between low amplification efficiency and A/T-rich regions at either end of the sequence for these two models (see Supplementary Figs. 11 and 12). This low GC-content, particularly localized to within a few nucleotides downstream of the adapter, is known to affect amplification efficiency due to next-base effects^48,49^ and the GC-preferences of Q5 polymerase^11^ and processivity-enhancing dsDNA binding domains^50,51^. A similar argument might also explain the GC-bias extracted by `CluMo` in the data of Song et al.^47^ using an error-prone polymerase for amplification.^24^

### External validation of model performance and motif effects

To validate both the performance of the trained 1D-CNN models and the inhibitory effects of the identified motifs, another oligo pool was prepared and amplified in an external laboratory. This pool, containing 10,000 random sequences and 2,000 sequences with deliberately inserted motifs, also underwent serial amplification, once with the experimental conditions of GCall and GCfix (KAPA SYBR FAST, 54 *^◦^*C annealing), and once with an altered protocol based on Erlich et al.^19^ (i.e., Q5 HiFi polymerase and 60 *^◦^*C annealing, see Fig. 6a and Methods).

**Fig. 6.**
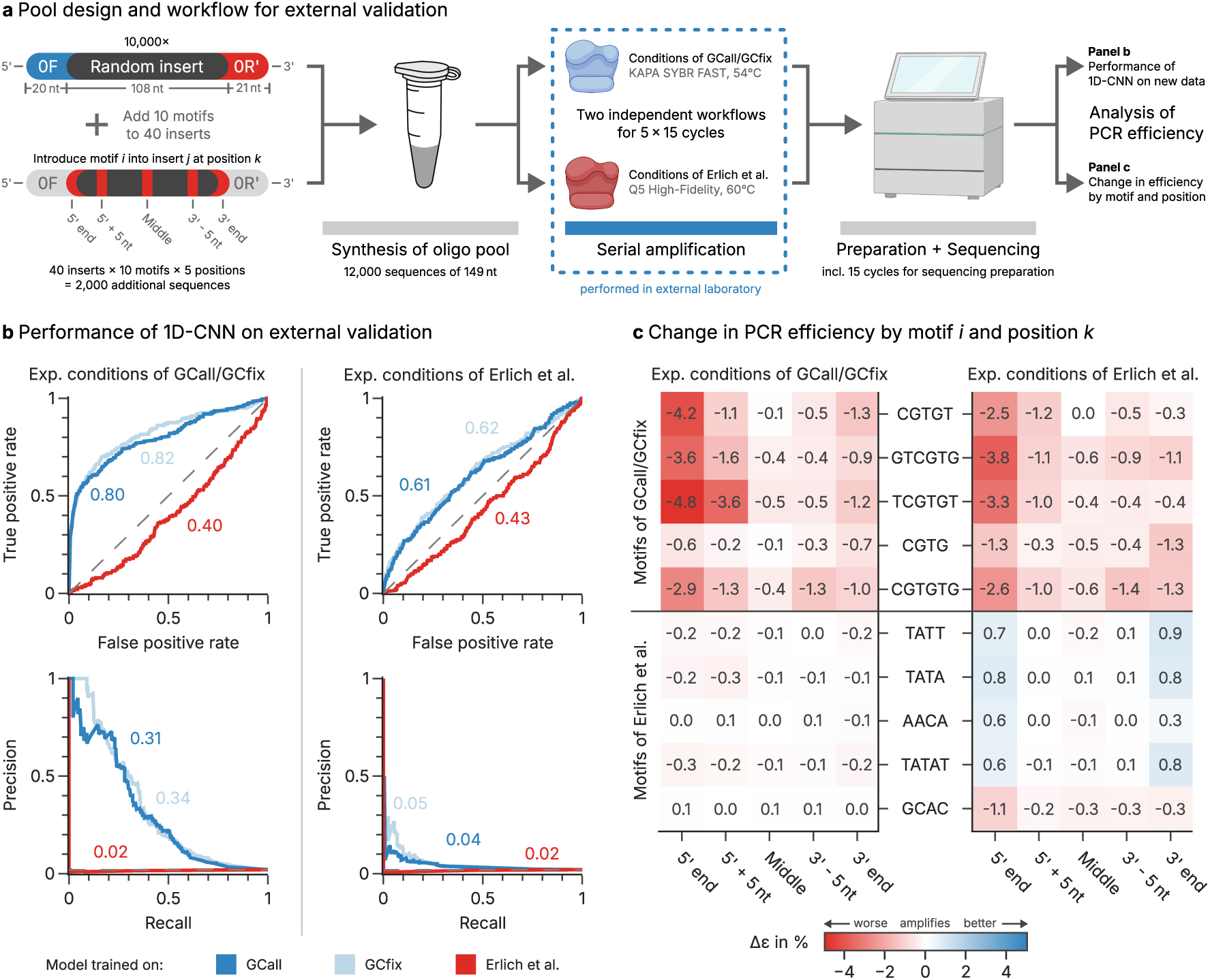
Externally validating model performance and motif discovery with a motif-enriched oligonucleotide pool. **(a)** For external validation, a new oligonucleotide pool consisting of 10,000 random sequences and an additional 2,000 sequences with deliberately inserted motifs was used. This pool underwent serial amplification in two independent, parallel workflows, which differed in the polymerase and the temperature profile used during PCR. The resulting sequencing data and estimated PCR efficiencies were used for external validation of model performance and motif discovery. **(b)** Performance metrics of the 1D-CNN models trained on the GCall (dark blue), GCfix (light blue), or Erlich et al. (red) datasets when tested on the data of the external validation experiments. The area under the receiver operating characteristic (AUROC, left) and the area under the precision-recall curve (AUPRC, right) are shown in the plot. The performance metrics are shown both for the datasets created with the conditions of GCall/GCfix (left column) and Erlich et al. (right column). **(c)** The mean change in amplification efficiency (*Δɛ*) across all sequences with a specific, deliberately inserted motif (vertical axis) at a specific position (horizontal axis) compared to the sequence without inserted motif. In both workflows (left column: conditions of GCall/GCfix, right column: Erlich et al.), the presence of the motifs identified during motif discovery for the GCall/GCfix models lead to a considerable decrease in amplification efficiency if present at, or close to, the 5’ end of the sequence.

Focussing on the experimental conditions of GCall and GCfix, we first evaluated the performance of the models trained with the GCall and GCfix data on the external validation dataset, using only the sequences with random inserts. Reassuringly, these models reached virtually identical predictive power as previously found in internal validation (AUROC 0.8, AUPRC 0.3, see left half of Fig. 6b), underlining the robustness of the models and the reproducibility of the PCR bias across laboratories. Turning to the sequences with inserted motifs, we quantified the amplification disadvantage conveyed by a deliberately inserted motif with the change in PCR efficiency between an individual sequence and its copy with the inserted motif (replacing existing nucleotides, see Fig. 6c). Across the ten motifs deliberately inserted into forty randomly selected sequences at five different positions each (see Fig. 6a), we observed profound inhibitory effects on PCR efficiency depending on motif and position (see Fig. 6c, left). In line with the hypothesized mechanism of hairpin-mediated self-priming shown in Fig. 4f, the effect of *CGTG*-derived motifs were most prominent at or close to the 5’- and 3’-ends. At the extreme, the motif *TCGTGT* inserted directly downstream of the 5’-adapter led to a mean decrease in PCR efficiency of 4.8 ± 2.4%, equivalent to a halving of the relative abundance approximately every 14 cycles.

Focussing next on the experimental conditions altered to reflect the polymerase and annealing temperature chosen by Erlich et al.^19^, the aforementioned workflow-dependence of the PCR bias becomes obvious. Assessing the performance of the model trained with the literature data of Erlich et al.^19^ on the validation dataset revealed a very poor predictive power (AUROC 0.43, AUPRC 0.02; see the right half panel of Fig. 6b). Interestingly, a residual inhibitory effect on amplification efficiency is still conveyed by the motifs previously identified to partake in adapter-mediated self-priming (see Fig. 6c, upper right). This is not surprising, given that a key difference between the experimental conditions by Erlich et al.^19^ and those used here is the choice of adapters, which appears to supersede or otherwise interact with the GC-based PCR bias conveyed by the Q5 polymerase.

Taken together, the results of the external validation strongly support the motif effects identified by `CluMo` and the robustness of the trained models. At the same time, the poor performance of the models in the external validation using a mixed condition of polymerase, annealing temperature, and adapters illustrates the strong workflow dependence of the identified PCR bias. Nonetheless, the external validation proved that, as long as all amplification conditions are kept identical, the 1D-CNN models trained on the GCall/GCfix datasets perform well on data generated by an external laboratory and exhibit the same motif-dependent amplification inhibition that we hypothesize to be caused by adapter-mediated self-priming.

## Discussion

Leveraging the versatility and flexibility of synthetic oligonucleotide pools, this work identifies multiple motif-dependent biases in multi-template PCR using explainable machine learning. For this, an experimental method was established to reproducibly annotate synthetic sequences with their PCR efficiencies, and a novel framework of motif discovery and analysis – called `CluMo`– was developed to interpret deep learning models trained on this data. With these tools, we identify a PCR bias affecting about 2% of random sequences in our datasets, and exploit the insights generated by explainable machine learning to pinpoint short position-dependent sequence motifs conveying an amplification disadvantage by adapter-mediated self-priming (shown in Fig. 4f).

Compared to the existing body of research^5,6,9–11,27^ on PCR-induced bias in multi-template PCR, our use of synthetic oligonucleotide pools overcomes key bottlenecks which previously precluded exploitation of deep learning, namely dataset size and annotation quality. We found 1D-CNN models with positional encoding can accurately identify poorly amplifying sequences in our datasets (AUROC *>*0.8, AUPRC *>*0.4), and demonstrated their reliability with data generated in an external laboratory. Importantly, using deep learning enabled a hypothesis-free investigation into PCR bias, without relying on pre-defined predictors such as GC content or k-mers. As a result, testing model generalization with literature datasets exposed the workflow-dependence of PCR bias while interpreting model performance with `CluMo` revealed adapter- and polymerase-specific motifs as its source. The learning from this analysis also applies to classical single-template PCR, where short-motif self-priming should also be accounted for to optimize amplification efficiency.

While hairpin formation is a common design consideration for primers^22^, its ability to enable self-priming of templates by short-ranged, motif-driven interactions with PCR adapters has only been reported for reverse transcription^52–54^ (i.e., at 37°C). However, our thermodynamic analysis and qPCR-based experiments suggest that these motif-adapter hairpins convey an amplification disadvantage in multi-template PCR by adapter-mediated self-priming, despite PCR’s higher temperatures (54-60 *^◦^*C) and the motifs’ short lengths (4-6 nt).

`CluMo` extends beyond the specific application in this work and provides a general framework for interpreting any deep neural networks utilizing sequence information. It identifies sequence motifs relevant to a user-defined task, and quantifies their global significance and their interactions, generalizing attribution methods, such as DeepLIFT^40^ and SHAP^41^, that solely analyze the independent contribution of single nucleotide variants within a given sequence. Notably, unlike traditional methods designed to visualize convolutional filters in the first layer of a CNN, such as DeepBind^30^, DeepSEA^31^, and CKN^55^, our approach is model-agnostic that can be flexibly applied to deep or other types of NNs. This versatility allows it to be applied to a broad range of sequence analysis tasks.

We envision accurate predictors for amplification biases and the tools required to develop them, such as those presented in this work, will find applications in many fields. Besides their use in constrained coding for DNA data storage^19,20^, research without alternatives to PCR-based workflows for high-throughput sequencing - such as metabarcoding or RNA sequencing - will benefit from the ability to identify and correct for PCR-induced biases in quantitative sequencing data.

## Methods

### Design of oligonucleotide pools

All oligonucleotide pools used in the experiments of this study were purchased from Twist Biosciences (Piscataway, NJ, United States) and designed with a fixed length of 149 nt. In all oligo pools, each design sequence contained a unique subsequence of 108 nt flanked by primer adapters (0F, 20 nt and 0R, 21 nt) for sequencing preparation following the Illumina Truseq protocol, according to established protocols.^3^

The two oligonucleotide pools comprised 12,000 sequences – either with (GCfix) or without (GCall) a fixed GC content of 50% – contained fully randomly generated subsequences. These pools and their sequencing data have previously been used to estimate error rates and biases in the DNA data storage workflow.^28^

The test pool used for assessing the reproducibility of the amplification bias comprised 1,000 sequences which were selected in equal parts from the sequences in the GCall and GCfix pools. The selection of sequences was based on the parameter estimates of two fitting procedures: one set of parameter estimates using the simple exponential PCR equation of Eq. 1, and a second set of parameter estimates using an extended fitting procedure including a parameter for sequencing efficiency and an estimate of parameter variability (not discussed in main text, see Supplementary Note 2). For both fitting procedures, the best- and worst-performing 100 sequences of the GCall and GCfix pools were selected. Due to a small intersection of these two sets, a total of 400 sequences were selected from the simple parameter-fitting procedure and only 346 sequences from the extended parameter-fitting procedure. The remaining 254 sequences were randomly selected from the sets of sequences from both pools. As the estimates of the simple parameter-fitting procedure matched the experimental results at least as well as the extended parameter-fitting procedure (see Supplementary Fig. 19 and Supplementary Note 2), the 346 sequences of the latter were not considered further. As a result, the analysis in Fig. 2e only shows the 654 individual sequences selected from the simple parameter-fitting procedure and the random sampling.

The validation pool used for external validation of the machine learning model and the effect of motifs comprised a total of 12,000 sequences. Of those, 10,000 were fully randomly generated without any constraint on GC content. For the remaining 2,000 sequences, we selected the five most significant motifs inferred from the models trained on the GCall/GCfix datasets or the literature dataset by Erlich et al.^19^ and created additional sequences with these motifs. To do so, 40 sequences of the randomly generated subset that did not already contain any of the motifs were selected, and each motif was inserted into each sequence once at the start, the end, the middle, 5 nt from the start, and 5 nt from the end of the sequence, replacing the nucleotides present there. This resulted in 50 additional sequences for each of the 40 sequences selected from the subset, for a total of 2,000 additional non-random sequences. Due to the overlapping nature of some motifs (e.g. motif CGTG is contained in motif CGTGT), the resulting set of sequences contained duplicates (e.g. if the sequence randomly featured T at the position following the inserted motif CGTG). In the analysis of the sequencing data, these sequences were always associated with the longer motif (e.g. to CGTGT in the example above) to isolate the effects of short motifs as best as possible.

### Serial amplification of pools

All oligonucleotide pools were dissolved to 10 ng µL^-1^ in ultrapure water, vortexed, and a 500x dilution in ultrapure water was used as starting point for serial amplification. All experiments except the second serial amplification of the validation pool used KAPA SYBR FAST polymerase master mix from Sigma-Aldrich (St. Louis, MI, United States) for amplification, employing a temperature profile with an initial denaturation at 95 *^◦^*C for 3 min, followed by 15 cycles at 95 *^◦^*C for 15 s, 54 *^◦^*C for 30 s, and 72 *^◦^*C for 30 s. Finally, a final extension at 72 *^◦^*C for 3 min was performed. The second serial amplification of the validation pool used Q5 Hot Start High-Fidelity master mix (Catalog# M0494) from New England Biolabs (Ipswich, MA, United States) instead, employing a temperature profile with an initial denaturation at 98 *^◦^*C for 30 s, followed by 15 cycles at 98 *^◦^*C for 10 s, 60 *^◦^*C for 30 s, and 72 *^◦^*C for 30 s.^19^ Finally, a final extension at 72 *^◦^*C for 5 min was performed. For each amplification, 5 µL of sample were mixed with 10 µL of 2x master mix, 3 µL of ultrapure water, and 1 µL each of the forward and reverse primer at 10 µM (Microsynth AG, Balgach, Switzerland). An overview of the primer sequences used in our experiments is given in Supplementary Table 6.

The serial amplification of oligonucleotide pools followed an iterative protocol, as previously described,^28^ to prevent resource exhaustion during PCR. In short, each iteration started by diluting 1 µL of the sample from the previous iteration by a factor of 3800x in ultrapure water (or 7600x, if the sample had approached the plateau phase after 15 cycles in the previous iteration). Then, the sample was amplified for 15 cycles in two wells: once using the standard primers (0F/0R), and once using primers with an overhang containing indexed sequencing adapters (2FUF/2RIF). The PCR product with sequencing adapters was then stored at −20 *^◦^*C, whereas the PCR product with the standard primers was directly used for the next iteration.

In the case of the validation pool, the amplifications with the standard primers (0F/0R) were performed at the Institute of Microbiology of the University of Stuttgart. The amplification with sequencing adapters (2FUF/2RIF) of the validation pool, as well all other serial amplification experiments and sequencing were performed at the Functional Materials Laboratory of ETH Zurich.

### Sequencing and data preprocessing

The PCR product with indexed sequencing adapters was purified by excision of the appropriate band on an agarose gel (E-Gel EX Agarose Gels 2%, Invitrogen) with a 50 bp ladder (Invitrogen), and subsequent spin-column purification (ZymoClean Gel DNA Recovery Kit, ZymoResearch). All samples were quantified by fluorescence (Qubit dsDNA HS Kit, Invitrogen) prior to dilution to 1 nM with ultrapure water. Multiple samples were then pooled, further diluted to 50 pM, and finally sequenced on the iSeq 100 (Illumina) with 150 bp paired reads.

The demultiplexed sequencing data was post-processed by adapter trimming and read mapping using BBMap^56^ (v39.01) against the pool’s reference sequences. Reference sequences whose reads occurred in fewer than two sequencing runs across the dataset were removed from the data. The read counts for all remaining reference sequences, normalized by the mean number of reads per reference sequence in the dataset, were used as coverage distributions for further analysis.

### Parameter Estimation from Coverage Distributions

To estimate the synthesis bias *x_i_*(0) and the relative amplification efficiency *ɛ_i_* of each reference sequence *i* in a set of serial amplification experiments, we model the evolution of the relative sequence coverage *x_i_*(*c_j_*) after *c_j_* cycles as shown in Eq. 1.^5,6,28^

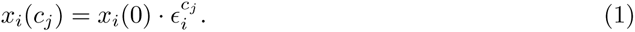

Full estimation of the parameters of all *N* reference sequences across all *M* serial amplification experiments is given by the solution to the least-squares problem of the sparse, log-linearized system of equations described by the PCR model in Eq. 2:

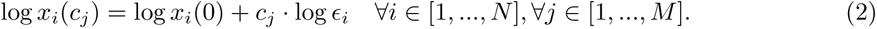

The estimated parameters were finally normalized to their mean. To validate the chosen approach, artificial sequencing data was generated *in-silico* using different defined distributions of initial synthesis bias and relative amplification efficiency. In a first test, a simple model based on Eq. 1 was used to generate sequencing data without the stochastic effects of PCR, dilution, and sequencing. In a second test, the full workflow was implemented in a digital twin of the DNA data storage process^28^ to investigate the approach’s robustness against stochastic noise. These validations of the model showed sufficient reliability of the parameter estimates under the expected experimental noise (e.g., stochastic sampling, or stochastic PCR effects) and no sensitivity to the underlying distribution of the parameters (see Supplementary Figure 26). Consequently, it was chosen over a more complex statistical model (see Supplementary Note 3 and Supplementary Figure 19). However, the accuracy of the parameter estimates decreased if a sequence was observed only in few sequencing runs (Supplementary Figure 26). Thus, we did not consider sequences which occurred less than two times across a set of sequencing data in the following analysis.

### Efficiency measurements by qPCR

Experimental quantification of the qPCR efficiency was performed for four arbitrarily selected sequences of the GCall oligonucleotide pool. Of the four selected sequences, #00006 had an estimated amplification efficiency of 0.999, #01634 of 1.014, #11493 of 0.854, and the efficiency of #09807 could not be estimated because it was filtered out due to occurring in only one sequencing run (relative coverage after first amplification: 0.21, thereafter 0). Sequence #09807 was included nonetheless to assess whether such sequences were missing in the sequencing data due to stochastic effects or because of an extremely poor amplification efficiency. These oligonucleotide sequences were synthesized individually by Microsynth (Balgach, Switzerland), dissolved with ultrapure water to 100 µM, serially diluted five times by factors of 10x, and each dilution measured by qPCR with the standard primers (0F/0R) in duplicates to create a calibration curve. A total of three independent dilutions and qPCR runs were performed for each oligonucleotide sequence, with the results shown in Supplementary Figure 2 and Supplementary Table 2. The qPCR efficiency of the GCall oligonucleotide pool itself was also measured in triplicates with the same range of dilutions. A comparison of qPCR-derived efficiencies was performed using one-way independent ANOVA after testing for homoscedasticity with Levene’s test, followed by post-hoc testing using Tukey’s range test.

An additional qPCR-based efficiency measurement of the four sequences (#00006, #01634, #09807, and #11493) with 5’-degenerate primers (four W at the 5’-end) was performed as above. The results of the calibration curves are shown in Supplementary Figure 3.

### Selection of literature datasets

Multiple additional sequencing datasets from the literature were selected to test and train the classification model. For parameter estimation, datasets must include sequencing data for at least two different PCR cycle counts and their sequencing coverage must be sufficiently high to yield accurate coverage distributions. Moreover, to preclude any possible bias stemming from sequences with biological function or extreme sequence properties (such as GC content or long nucleotide repeats), we limited our search to datasets derived from synthetic oligonucleotide pools which contain close-to-random sequences. Multiple such datasets were identified in the DNA data storage literature, from Erlich et al.^19^, Koch et al.^46^, Song et al.^47^, Choi et al.^18^, and Gao et al.^17^ and processed as described above. A detailed overview of the experimental parameters and sequencing endpoints of all literature datasets is given in Supplementary Table 7.

### Machine learning models

In this work, the main model we propose to use is a 1D-CNN model to predict whether a DNA sequence is of low PCR amplification efficiency given the sequence data. Due to the absence of prior knowledge regarding the specific threshold for low PCR efficiency in randomly synthesized sequences, we empirically determine a 2% threshold to differentiate between low-efficiency sequences and normal-efficiency sequences (see above). This binary categorization then facilitates the formulation of the prediction task as a classification problem.

1D-CNN models have demonstrated superior efficacy compared to traditional machine learning methods across a wide range of DNA sequence property prediction tasks, as discussed in the introduction. While 1D-CNNs can implicitly capture sequential order through multiple layers, they lack explicit modeling of the nucleotide positions. This explicit positional information could be crucial for identifying motifs, especially in shallow models where the implicit capture of order might not be sufficient. For example, in eukaryotic chromosomes, telomeres are repetitive nucleotide sequences at each end of a chromosome. The specific motif sequence often comprises repetitions of a short DNA sequence (like ‘TTAGGG’ in vertebrates). The positional specificity of these motifs at the very ends of the chromosomes is vital for their function in protecting the chromosome from deterioration or fusion with neighboring chromosomes^57,58^. Motivated by this, an additional positional encoding (PE) component, which was first introduced with the transformer model^59^, is incorporated into the 1D-CNN model.

Specifically, positional encoding represents nucleotide positions as real-valued vectors, which can then be added to the embeddings of nucleotides. It is defined as:

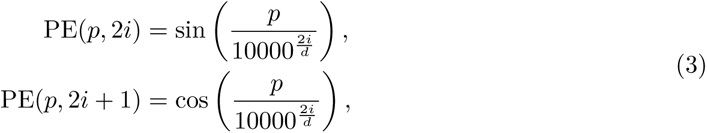

where sinusoidal functions are used to encode each position *p* into a vector of dimension *d* (assumed to be an even number), and *i* ∈ [0*, ^d^/*2] represents the dimension index as used in Vaswani et al. 2017^59^. In our implementation, the one-hot encoded DNA sequence is first projected into a higher dimensional space (of the same dimension as PE’s dimension). Then, we perform an element-wise addition between the PE and the high-dimensional projection of the sequence.

#### Baseline models

To demonstrate the efficacy of the 1D-CNN model in identifying sequences with lower PCR efficiency, we established several baseline models for comparative evaluation of the proposed 1D-CNN model with PE. The baseline models include:

- 1D CNN model without PE.
- Recurrent neural network (RNN)-based model.
- Lasso regularized logistic regression (LR) model with the frequency of each nucleotide and the GC content in the sequence.

More details of the baseline models and the architecture of our proposed model can be found in Supplementary Note 1.

#### Experimental setup

Given the binary classification nature of the task, we minimize the binary cross-entropy loss during training. We use two metrics for our analysis: the Area Under the Receiver Operating Characteristic (AUROC) and the Area Under the Precision-Recall Curve (AUPRC). We consider two evaluations: the standard within-dataset evaluation and external validation. For within-dataset evaluation, we employ stratified 5-fold nested cross-validation (CV) to preserve the percentage of samples for each class across folds. In each fold, 10% of the training data is selected as the validation set, which is used only for hyperparameter selection. We perform a randomized search^60^ over 50 iterations, each corresponding to a hyperparameter configuration including the learning rate, width, depth, batch size, weight decay, dropout, etc, detailed in Table 1 with an equivalent search conducted for the RNN-based model as described in supplementary Table 1. The best-performing model parameter configuration is determined by the highest AUPRC over all 5 validation sets and we report the mean and std of the performance on the 5 test sets. To assess the generalizability of the model, the best-performing hyperparameters from within-dataset evaluations are used to train the model on the entire dataset, which is subsequently tested on different external datasets. We also use class weights inversely proportional to the frequency ratio of the positive class to the negative class, enabling the loss function to more effectively balance the contribution of each class during training. The detailed architecture of the model is illustrated in Figure 3.

**Table 1.** Hyperparameter grid and ranges for the hyperparameter search of the 1D-CNN model.

| Hyperparameter | Search values |
| --- | --- |
| Number of convolutional layers | 1, 2, 3 |
| Number of convolutional filters | 32, 64, 128 |
| Length of convolutional filters | 4, 8, 12 |
| Learning rate | $10^{-3}$, $10^{-4}$, $10^{-5}$ |
| Global pooling | mean, max |
| Batch size | 64, 128, 256 |
| Weight decay | 0, $10^{-3}$, $10^{-4}$ |

### CluMo: Motif Discovery via Attribution and Clustering

DNA motifs are short and recurring subsequences found within DNA sequences, which are believed to play critical biological roles. In this study, we introduce `CluMo`, a novel method for discovering functional motifs within sequences. Unlike traditional motif discovery tools, `CluMo` specifically targets motifs associated with user-defined functions, such as PCR amplification efficiency *ɛ*. It achieves this by integrating feature attribution techniques with k-mer analysis and clustering. This general approach is applicable to any biological sequence and allows researchers to uncover motifs significantly correlated with any sequence property or function predicted by a deep learning model. We will now delve into the details of `CluMo`.

#### Step 1: Feature attribution analysis

`CluMo` begins by applying a feature attribution method to interpret the prediction made by the trained deep model. In this study, we use DeepLIFT (Deep Learning Important FeaTures)^40^, a method that assigns attribution scores to each nucleotide by comparing the activation of neurons to reference activation. Note that our approach is not limited to DeepLIFT and can leverage any attribution analysis method as a replacement, such as Integrated Gradients^61^ and SHAP^41^. Using the chosen feature attribution method, nucleotide-level attribution scores are obtained, indicating the impact of each individual nucleotide on the model prediction of each DNA sequence.

#### Step 2: Significant k-mers identification

To identify the sequence segments with the strongest influence on the model’s prediction, we utilize a sliding window approach. We iterate through the sequence with window sizes ranging from 4 to 12 nucleotides (k-mers). For each window size and sequence position, we calculate a cumulative attribution score. This score is the sum of the individual attribution scores (obtained in the previous step) assigned by the deep learning model to each nucleotide within the window. The k-mer with the highest cumulative score, for a given window size, represents the subsequence with the most significant impact on the model’s prediction for that specific region of the sequence. These k-mers are selected as the candidates for the next step.

#### Step 3: k-mers clustering

We then analyze the relationships between the identified k-mers. To achieve this, we calculate the Hamming distance between each pair of k-mers for each window size. The Hamming distance simply counts the number of nucleotide positions that differ between two k-mers. Next, we employ t-SNE (t-distributed Stochastic Neighbor Embedding)^62^ to embed the k-mers into a lower-dimensional (2D) space. This allows us to visualize the relationships between k-mers based on their sequence similarity. Next, we utilize weighted k-means clustering^63^ to group similar k-mers. This approach assigns greater weight to more frequent k-mers, ensuring that prevalent sequence patterns exert a stronger influence on the formation of clusters.

Since we lack prior knowledge about the number of functional motifs present, we determine the optimal cluster number *C* using the silhouette score^64^ within the t-SNE embedded space. This score measures the cohesion within clusters and the separation between them. We systematically test values of *C* from 2 to 12 and select the one that maximizes the average silhouette score across all data points. Finally, for each cluster, we construct a Positional Weight Matrix (PWM) that captures the k-mer frequency within the cluster. This PWM essentially represents the motif for that cluster. Sequence logos can be generated from these PWMs, providing a visually intuitive representation of the identified motifs.

#### Step 4: Motif presence and enrichment analysis

To identify occurrences of the discovered motifs (PWMs) within a target sequence, we calculate a presence score. This score represents the strongest match between the PWM and any k-mer window across the entire target sequence. The calculation is as follows:

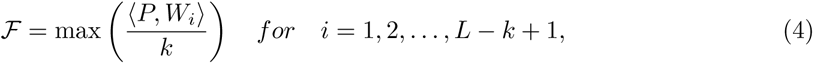

where *P* denotes the PWM representing the motif, *W_i_* represents the corresponding window in the sequence, *k* is the window size, and *L* is the length of the target sequence. Essentially, the presence score is the maximum dot product between the PWM and each possible k-mer window in the target sequence, normalized by the window size. A score of 1 indicates a perfect match between the PWM and a k-mer in the sequence, while scores below 1 reflect varying degrees of mismatch. We define a presence threshold of 0.5 for the score F. If the score for a motif in a given sequence exceeds this threshold, we consider the motif to be present. Otherwise, it is considered absent. Following this definition, we can analyze the association between motifs and a specific sequence property. Here, we focus on sequences with low PCR amplification efficiency (positive set). We create the contingency table summarizing the presence/absence of each motif across the test sequences.

Next, we employ a chi-squared test (*χ*^2^ test) to assess the statistical significance of each motif’s association with the positive set. The null hypothesis for this test is that there is no connection between the presence of a motif and a sequence belonging to the low PCR efficiency group. The chi-squared statistic is calculated by comparing the observed frequencies of motif presence/absence in the contingency table with the expected frequencies under the null hypothesis. A statistically significant chi-squared value (p-value *< α* with *α* = 0.05) indicates a rejection of the null hypothesis. This suggests a non-random association between the motif and the positive set, implying that the motif might influence PCR efficiency. To account for multiple testing when analyzing numerous motifs, we apply the Bonferroni correction. This adjusts the significance threshold to control for the increased chance of false positives.

#### Step 5: Motif substitution analysis

To evaluate the influence of each identified motif on the model’s prediction, we employ a motif substitution strategy. We substitute the discovered motifs (ranked by their p-value, with the most significant ones first) within the test set sequences. This substitution involves replacing each motif occurrence with an average subsequence. In simpler terms, the original motif is replaced with a random sequence segment with equal probabilities for each nucleotide.

This substitution process is performed solely on the test set, allowing us to directly compare the model’s performance on the original sequences versus the sequences with each motif substituted.

By observing these performance changes, we can quantify the impact of each motif on the model’s predictions, both within a single dataset and across different datasets.

The detailed algorithm `CluMo` is described in Algorithm 1.

##### Algorithm 1

`CluMo`: Motif Discovery via Attribution and Clustering

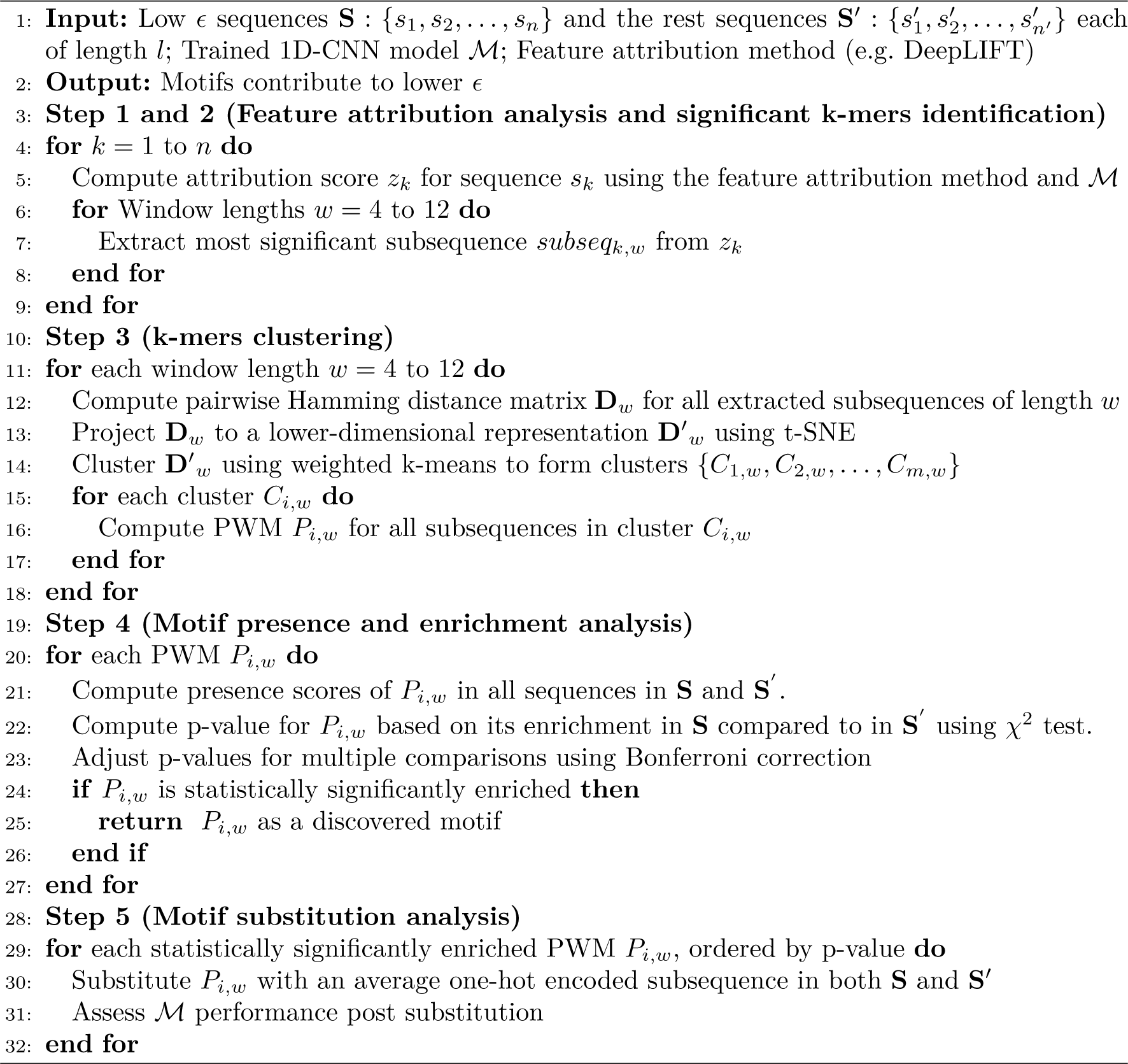

## Data availability

The experimental sequencing data generated in this study have been deposited in the European Nucleotide Archive under accession code PRJEB77604. Additional sequencing datasets are available from Gimpel et al.^28^ (PRJEB65931), Koch et al.^46^ (PRJEB35217), Erlich et al.^19^ (PRJEB19305 and PRJEB19307), Song et al.^47^ (doi:10.6084/m9.figshare.16727122.v2, doi:10.6084/m9.figshare.17193128.v1, doi:10.6084/m9.figshare.18515045.v1), Gao et al.^17^ (pers. communication), and Choi et al.^18^ (PRJNA555140).

## Code availability

The implementation of `CluMo`, as well as all scripts used for data analysis and plotting are available on GitHub: github.com/BorgwardtLab/PCR-bias.

## Acknowledgements

This project was partially financed by the European Union’s Horizon 2020 Program, FET-Open: DNA-FAIRYLIGHTS, grant agreement no. 964995, and the European Union’s Horizon 2020 Research and Innovation Program under Marie Sklodowska-Curie Grant Agreement No. 813533. Core funding by the Max Planck Society (to K.B.) and ETH Zürich (to W.J.S.) is acknowledged. Data analysis was performed on the Euler cluster operated by the High-Performance Computing group at ETH Zürich. Figures were partially created with BioRender.com.

## Competing interests

The authors declare no competing interests.

## Authors’ contributions

**Conceptualization:** K.B., R.N.G.; **Methodology:** K.B., R.N.G., W.J.S., B.C., A.L.G., B.F., D.C., M.H.; **Supervision:** D.C., K.B., R.N.G.; **Resources:** W.J.S., K.B., R.N.G.; **Investigation:** A.L.G., P.A., L.O.D.W.; **Software:** B.F., D.C., M.H.; **Data analysis:** A.L.G., B.F., D.C., M.H., L.M., B.C.; **Writing-Draft:** A.L.G., B.F., D.C., B.C., R.N.G., K.B.; **Visualization:** A.L.G., B.F., D.C., B.C., R.N.G., K.B.; **Writing-Review:** all authors.

## Notes

### Competing Interest Statement

The authors have declared no competing interest.

