## Supplementary Text, Figures and Tables for "Deep learning uncovers sequence-specific amplification bias in multi-template PCR"

### Supplementary Notes

#### Supplementary Note 1: Details of the deep learning models

- **Input:** One-hot encoded DNA sequences $x$.
- $x \leftarrow$ Linear projection$(x)$ + Positional embedding$(x)$
- For $i = 1, \cdots, N$:
  - $x \leftarrow$ 1D convolutional layer$(x)$
  - $x \leftarrow$ BatchNorm$(x)$
  - $x \leftarrow$ ReLU$(x)$
- $x \leftarrow$ Global pooling$(x)$
- $x \leftarrow$ Linear classification layer$(x)$

Detailed architecture of the 1D-CNN model with PE, where $N$ is the total number of convolutional layers.

#### Supplementary Note 2: Design and analysis of parameter validation pool

To assess the reproducibility of our assignment of amplification efficiencies, we conducted a serial amplification experiment with a new oligo pool comprising 1000 selected sequences from the GCall and GCfix pools (see Methods). The selection of sequences was primarily based on the sequences' estimated amplification efficiencies (i.e., 400 sequences with poor, 200 with normal, and 400 with superior efficiency), however, secondarily, the pool was also used to compare two methods for estimating amplification efficiencies from the sequencing data.

Specifically, of the 400 sequences with either poor or superior efficiency, half of each were selected based on the simple model of exponential amplification presented in the main text (see Equation 1 and 2). The other half was selected based on a more complex model of the whole process, including dilution and sequencing steps.

Supplementary Fig. 19b shows how the relative coverage of all sequences evolves during serial amplification. Evidently, both methods for estimating amplification efficiencies successfully identify poorly amplifying sequences, as the sequences identified by either model are quickly depleted in the sequencing data (i.e., around cycle 60). The simple model (model 1 in the figure) appears to perform slightly better in identifying poorly amplifying sequences compared to the complex model (model 2 in the figure), as evidenced by the faster loss of its selected sequences in the sequencing data (i.e., between cycles 30 and 60). For this reason, as well as the simplicity of this model, we selected the method using a simple exponential model of the amplification process as our method for estimating the amplification efficiencies in all analyses.

Accordingly, we limited the sequences shown in Fig. 2e and Supplementary Fig. 20 to those selected by the simple model described in the main text, omitting the sequences selected by the complex model entirely. This reduces the number of sequences in the analysis from 1,000 to 654.

#### Supplementary Note 3: Complex statistical model

We introduce a statistical model to account for the concentration distribution in each step of the proposed sequencing experiment, and this model is modified based on the statistical model introduced in[65] for RNAseq. In this model, each DNA present in the original preparation, whose level is indicated as $N_i$, where $i$ is the index of the individual sequence, and since the abundance of each sequence is the same in all replicated samples, the sample index $j$ is omitted. Then each sample was subjected to different sequences of combinations of dilution steps and PCR amplification steps. The PCR amplification steps are made up of blocks of 15 cycles each, and the dilution steps are interleaved between the PCR steps to reduce the impact of the PCR bias.

##### *Dilution:*

As the dilution steps will simply dilute the concentration of the DNA molecules (the amount of DNA per unit volume) in a linear manner, so these separate dilution steps can simply be multiplied. The mean and variance of the concentration $X_i$ can be formulated as:

$$E[X_{i,j}] = d_j N_{i,j} \tag{1}$$

$$Var[X_{i,j}] = d_j^2 N_{i,j} \tag{2}$$

where $d_j$ is the multiplication of all dilution steps subject to each individual sample $j$.

##### *PCR amplification:*

The accumulation of products during the PCR reaction may be stochastically modeled by a Galton-Watson (GW) branching process[66–69]. We will consider the exponential phase of the PCR reaction, in which the reaction efficiency is constant in all cycles and also in different samples $j$ for the same sequence $i$. In particular, the abundance of DNA sequence $i$ after the $n+1$ PCR cycle can be expressed by the Markovian relation[69]:

$$L_i^{n+1} | L_i^n, \epsilon_i = L_i^n + Bionmial(L_i^n, \epsilon_i) \tag{3}$$

So, the abundance of DNA sequence $i$ after the $n+1$ PCR cycle, $L_i^{n+1}$, is only conditioned on the abundance of DNA sequence $i$ after the $n$ PCR cycle and the PCR efficiency of the sequence, $\epsilon_i$.

Despite the analytical intractability of the GW distribution $P(L_i^n|X_i, \epsilon_i)$, the relationship between the mean and variance can be explicitly derived by standard branching theory results[66,69] as:

$$E[L_{i,j}] = N_{i,j}(1+\epsilon_i)^{N_j^{PCR}} \tag{4}$$

$$\begin{aligned} Var[L_{i,j}] &= N_{i,j}\frac{1-\epsilon_i}{1+\epsilon_i}(1+\epsilon_i)^{N_j^{PCR}}[(1+\epsilon_i)^{N_j^{PCR}} - 1] \\ &\approx N_{i,j}\frac{1-\epsilon_i}{1+\epsilon_i}(1+\epsilon_i)^{2N_j^{PCR}} \end{aligned} \tag{5}$$

***Sequencing Run:***

For simplicity of the statistical model, we use a single parameter $p$ to represent the probability that a given molecule existing in the PCR amplified library will generate a sequence entry in the output of the sequence. This probability encapsulates signal loss due to many subfactors in the sequencing run, e.g. sequencing depth, library preparation efficiency, sequencing technology efficiency, etc. The whole sequencing step can be viewed as a Poisson process where the final sequncing reads are ditributed according to the Poisson distribution. However, as discussed in the previous paragraphs, $L_{i,j}$ is a random variable, and this makes the Poisson process a mixed Poisson process which is a convolution of two distributions: the distribution of $L_{i,j}$ and the Poisson distribution defined by the sequencing probability $p$, and this yields the following statistical model for the final sequencing reads per sequence per sample $Y_{i,j}$:

$$Y_{i,j}|p, L_{i,j} \sim Poisson(pL_{i,j}) \tag{6}$$

$$\begin{aligned} E[Y_{i,j}] &= E[E[Y_{i,j}|L_{i,j}, p]] = pE[L_{i,j}] \\ &= p(1+\epsilon_i)^{N_j^{PCR}} d_j N_i \end{aligned} \tag{7}$$

$$\begin{aligned} Var[Y_{i,j}] &= Var[E[Y_{i,j}|L_{i,j}, p]] + E[Var[Y_{i,j}|L_{i,j}, p]] \\ &= E[Y_{i,j}](E[Y_{i,j}]\phi + 1) \end{aligned} \tag{8}$$

$$\phi_{i,j} = \frac{(1-\epsilon_i)}{E[X_{i,j}](1+\epsilon_i)} \tag{9}$$

The variance of the mixed Poisson distribution is determined by the law of the total variance.

$$Var(X) = E[Var(X \mid Y)] + Var(E[X \mid Y]) \tag{10}$$

In addition, extensive numerical simulations have been conducted to assess whether the mixed Poisson distribution by less complex functions in this study[65]. The authors proved that the Linear

Quadratic Normal family (LQNO) can be used as a less complex alternative to approximate the mixed Poisson distribution, which is defined as:

$$Y_{i,j} \sim LQNO(\mu_{i,j}) = \frac{exp(-(Y_{i,j} - \mu_{i,j})^2/2\sigma_{i,j}^2}{\sqrt{2\pi}\sigma_{i,j}} \tag{11}$$

$$\sigma_{i,j}^2 = \mu_{i,j}(1 + \mu_{i,j}\phi_{i,j}) \tag{12}$$

The overall statistical model is optimized by minimizing the negative log-likelihood (NLL), which quantifies the difference between the model predictions and the observed data (i.e. the final sequencing reads $Y_{i,j}$). The optimization algorithm iteratively adjusts the model parameters to minimize the objective function, and this process involves evaluating the objective function and its derivatives at various points in the parameter space.

As this complex statistical method did not perform better than a two parameter model (see Methods section and Supplementary Figure 19), the two parameter model was chosen for all further analysis.

### Supplementary Figures

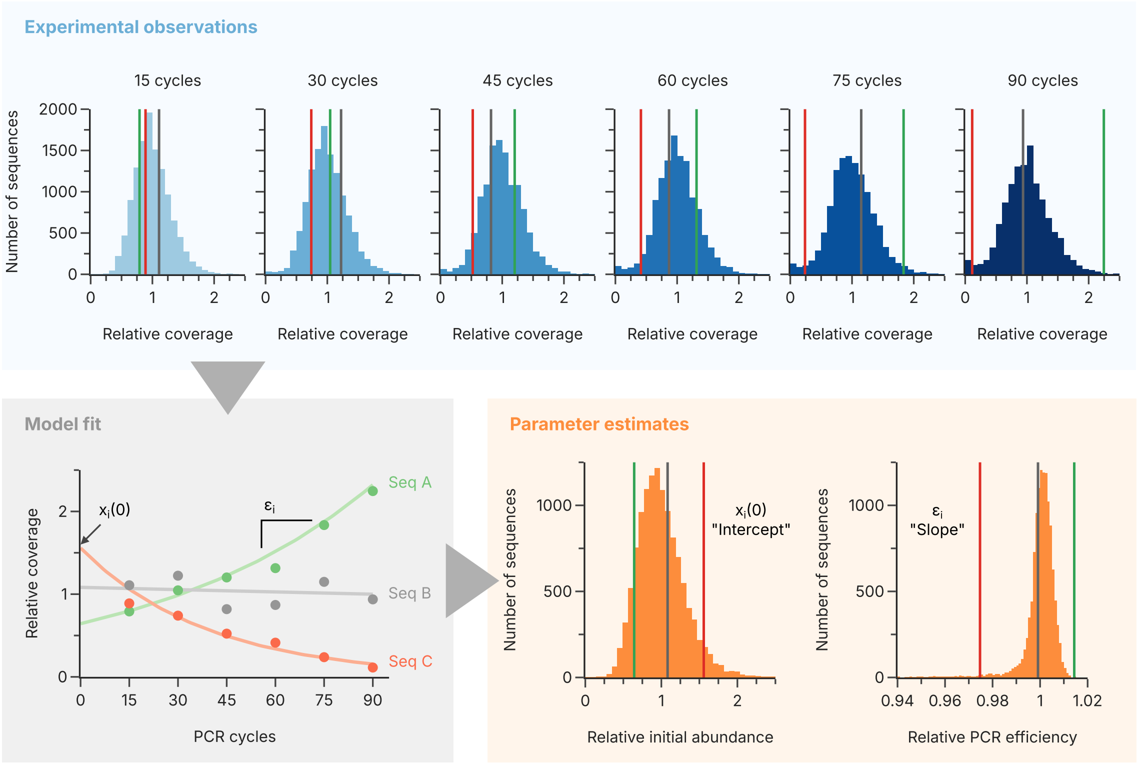


**Supplementary Fig. 1** Illustration of the model workflow. After each iteration of the serial amplification protocol, the sequencing data is mapped to the reference sequences to quantify the relative coverage of each sequence in the pool (top, blue box). For each sequence, the set of relative coverages at each iteration is used to fit the two parameters of the exponential PCR model: the initial abundance (intercept, $x_i(0)$) and the amplification efficiency (slope, $\epsilon_i$) (bottom left, gray box). The parameter estimates of all sequences in the pool yield distributions of the initial abundance and amplification efficiency (bottom right, orange box). All data shown corresponds to the GCall pool. The relative coverages and parameter estimates of three exemplary sequences (A, green, high amplification efficiency; B, gray, normal amplification efficiency; and C, red, low amplification efficiency) are highlighted across all panels.

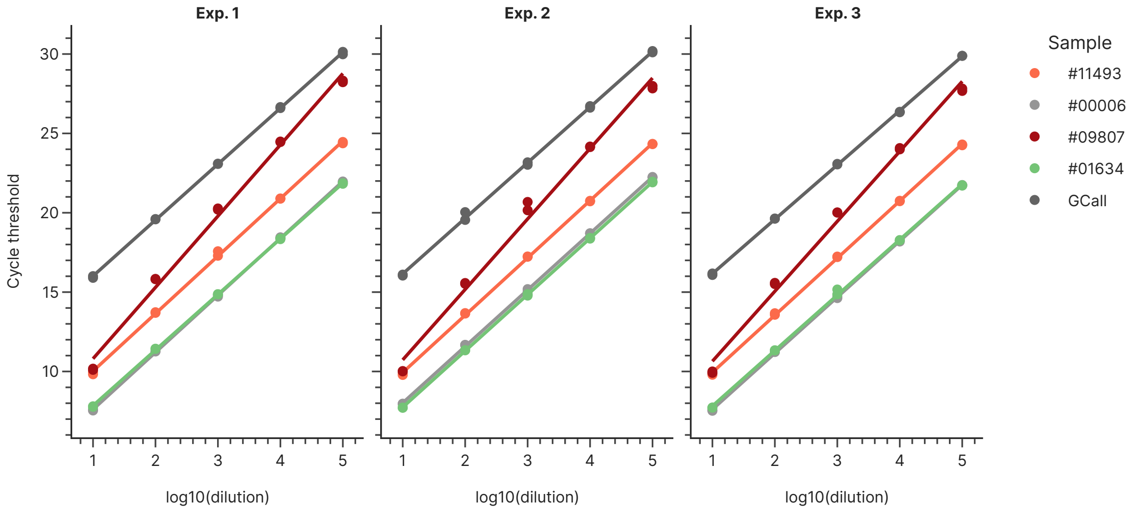


**Supplementary Fig. 2** qPCR dilution curves for selected sequences of the GCall pool. Four sequences from the GCall pool were selected based on their estimated amplification efficiency (see Methods), individually synthesized, and tested for amplification efficiency together with the GCall pool as a whole. The slopes of the dilution curves across three repetitions (Exp. 1 through Exp. 3) of the dilution and qPCR workflow were then used to estimate the amplification efficiency (see Supplementary Table 2).

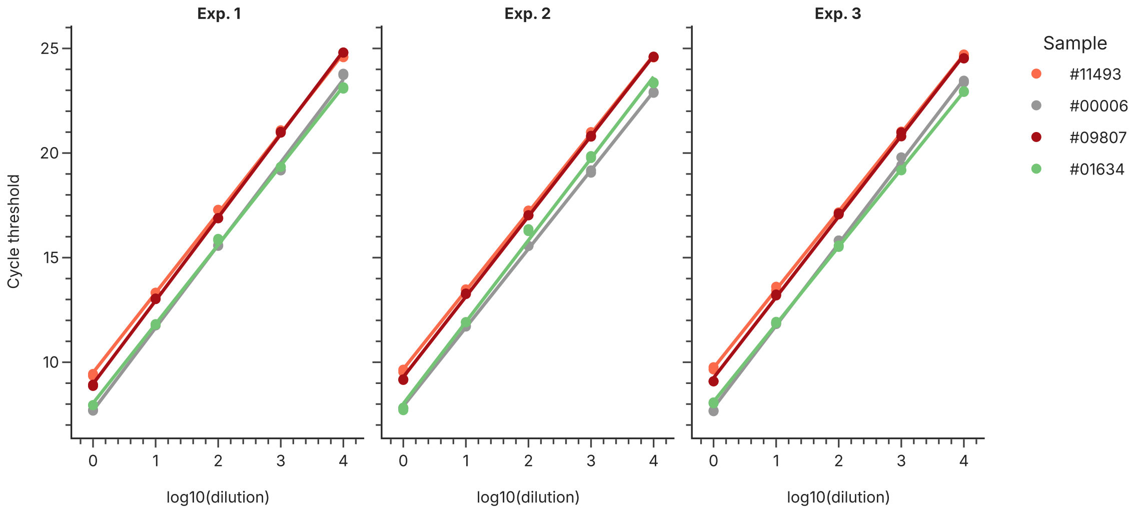


**Supplementary Fig. 3** qPCR dilution curves for selected sequences of the GCall pool amplified with degenerate primers. The same sequences outlined in Supplementary Fig. 2 were amplified with a degenerate primer to estimate their amplification efficiencies (see Methods).

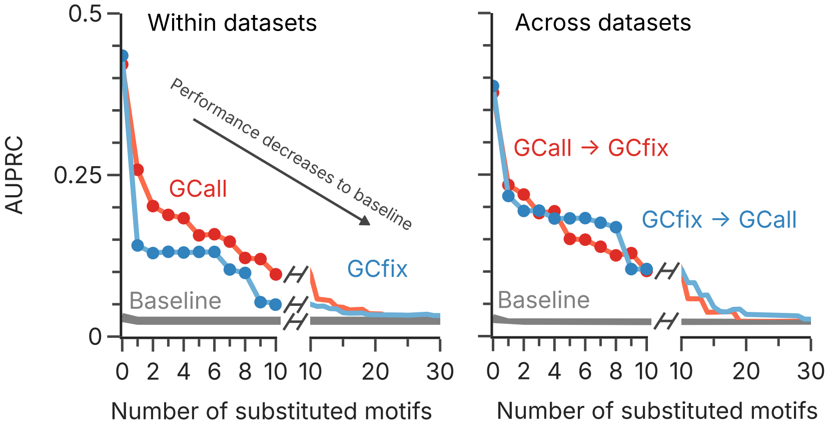


**Supplementary Fig. 4** 1D-CNN performance on the GCall and GCfix datasets during motif substitution using AUPRC. This figure shows the performance of the 1D-CNN models and the baseline models within (left) and across the GCall and GCfix datasets (right) as a function of the number of motifs replaced in the test data. The GCall model is shown in red, and the GCfix model is shown in blue. This figure is the equivalent to Fig. 4e using AUPRC as the performance metric.

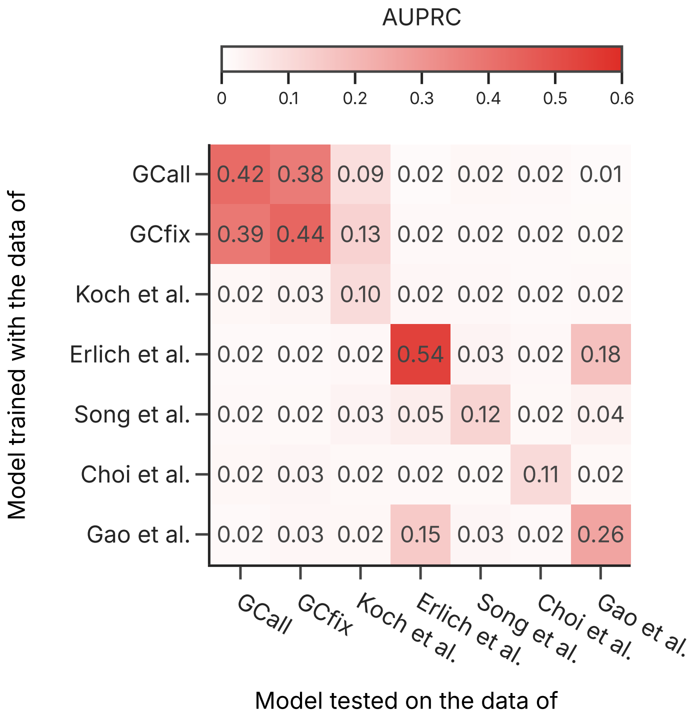


**Supplementary Fig. 5** 1D-CNN performance on all literature datasets using AUPRC. Heatmap of the area under the precision-recall curve (AUPRC) metric for the models trained and tested on the different literature datasets. This figure is equivalent to Fig. 5 using AUPRC as the performance metric.

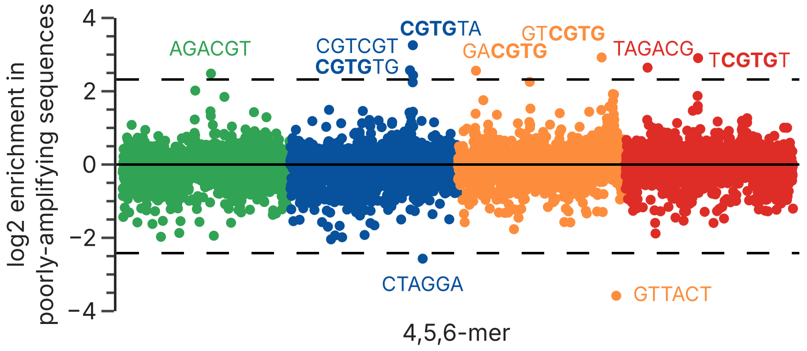


**Supplementary Fig. 6** K-mer frequency analysis for the combined GCall and GCfix datasets. All sequences of the GCall and GCfix pools were converted into overlapping k-mers (considering $k \in [4, 6]$), and their frequency in the positive (i.e. low efficiency) group compared to their frequency in the negative group (i.e. normal efficiency). For each k-mer, the plot shows its enrichment in the positive group. The solid horizontal line corresponds to an equal frequency in both groups (i.e. no enrichment), whereas the two dashed lines indicate five standard deviations (i.e. strong enrichment or depletion). k-mers outside of five standard deviations are annotated, with the CGTG submotif highlighted in bold where present. Colors indicate the starting nucleotide of the k-mer.

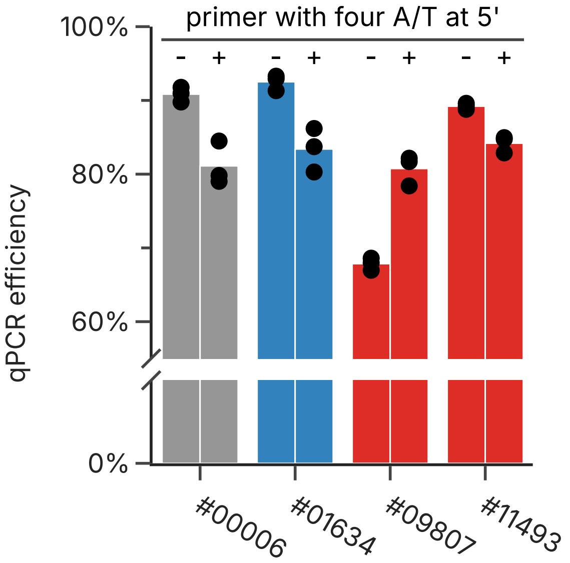


**Supplementary Fig. 7** Comparison of qPCR-based amplification efficiency for the four selected sequences of the GCall pool (see Methods), upon amplification with or without degeneracy in the primer. The qPCR-based amplification efficiency ($N = 3$) is based on the slope of the calibration curves shown in Supplementary Figs. 2 and 3.

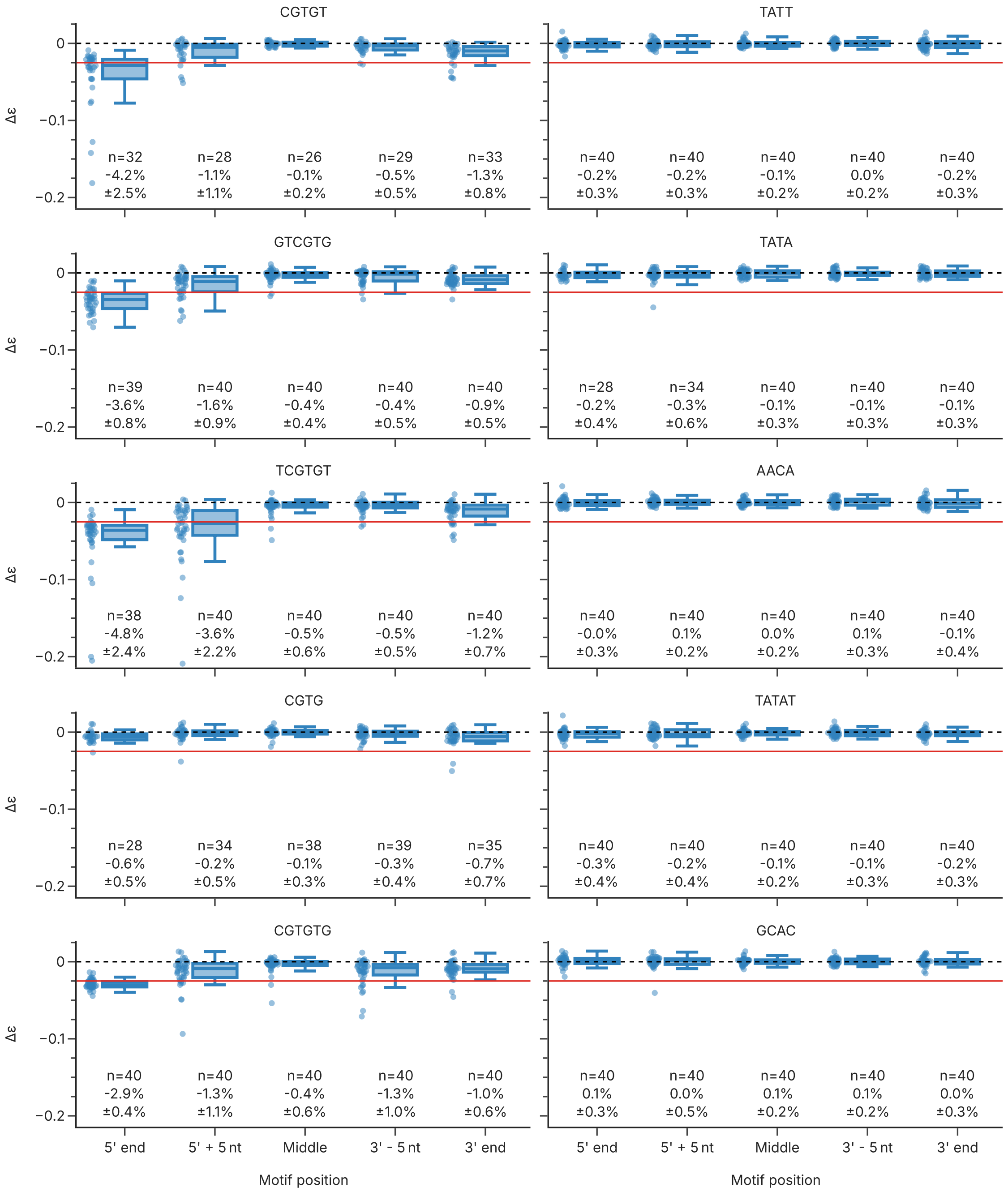


**Supplementary Fig. 8** Motif- and position-dependent change in amplification efficiencies for the workflow based on the GCall and GCfix pools. Each panel shows the difference in amplification efficiency ($\Delta\epsilon$) between the base sequence without motif and the same sequence with the corresponding motif inserted at the indicated position (x-axis). All motifs in the left column correspond to the GCall/GCfix datasets, whereas the motifs in the right column belong to the Erlich et al. dataset. The dotted line represent no change in amplification efficiency, whereas the solid red line indicates a considerable change in amplification efficiency (-2.5%, roughly the threshold corresponding to the classification as a poorly amplifying sequence in the GCall/GCfix datasets). The number of individual sequences ($n$) and the mean difference and the Bonferroni-corrected 95% confidence interval are also shown below each box.

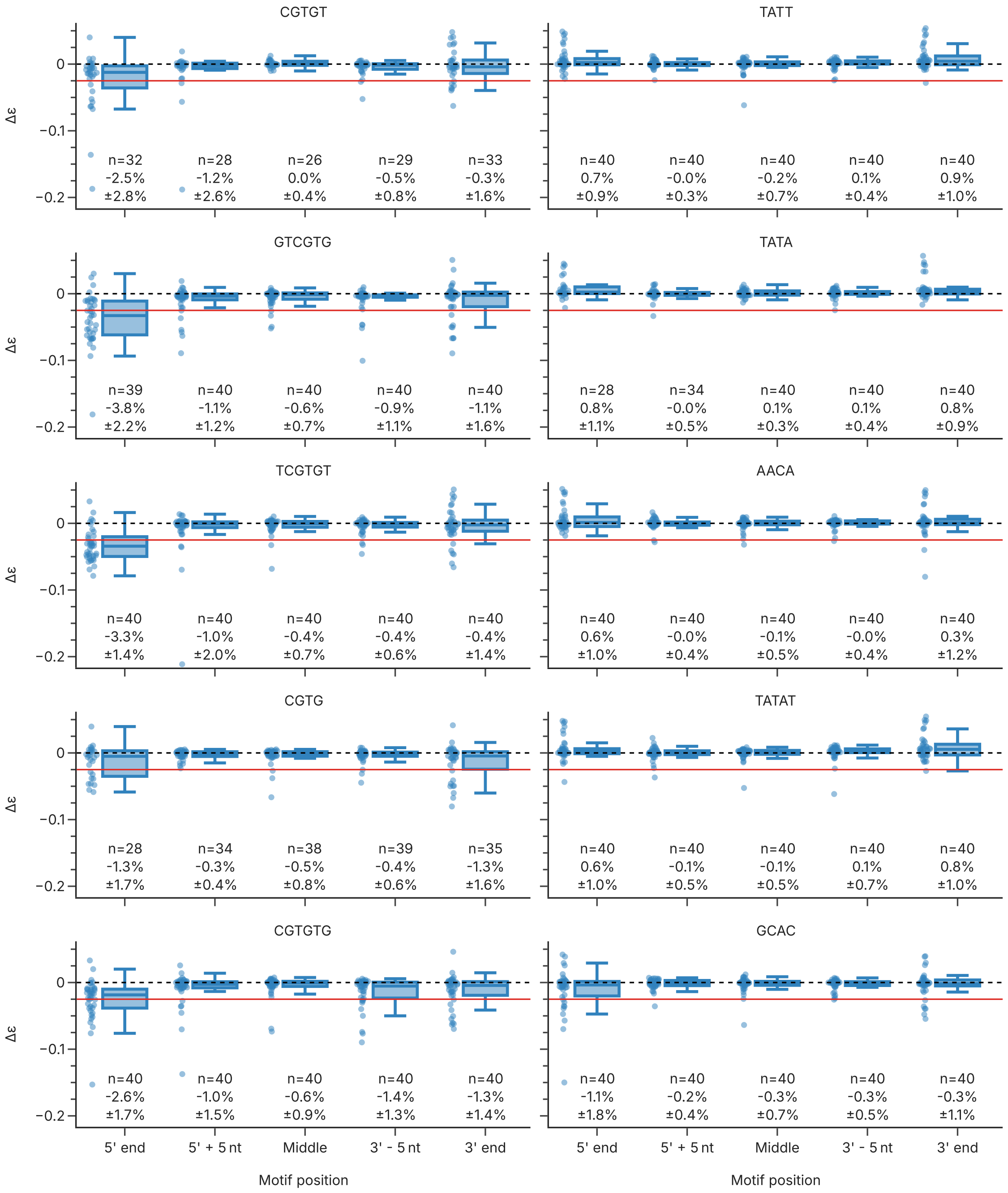


**Supplementary Fig. 9** Motif- and position-dependent change in amplification efficiencies for the workflow based on the Erlich et al. data. Each panel shows the difference in amplification efficiency ($\Delta\epsilon$) between the base sequence without motif and the same sequence with the corresponding motif inserted at the indicated position (x-axis). All motifs in the left column correspond to the GCall/GCfix datasets, whereas the motifs in the right column belong to the Erlich et al. dataset. The dotted line represents no change in amplification efficiency, whereas the solid red line indicates a considerable change in amplification efficiency (-2.5%, roughly the threshold corresponding to the classification as a poorly amplifying sequence in the GCall/GCfix datasets). The number of individual sequences ($n$) and the mean difference and the Bonferroni-corrected 95% confidence interval are also shown below each box.

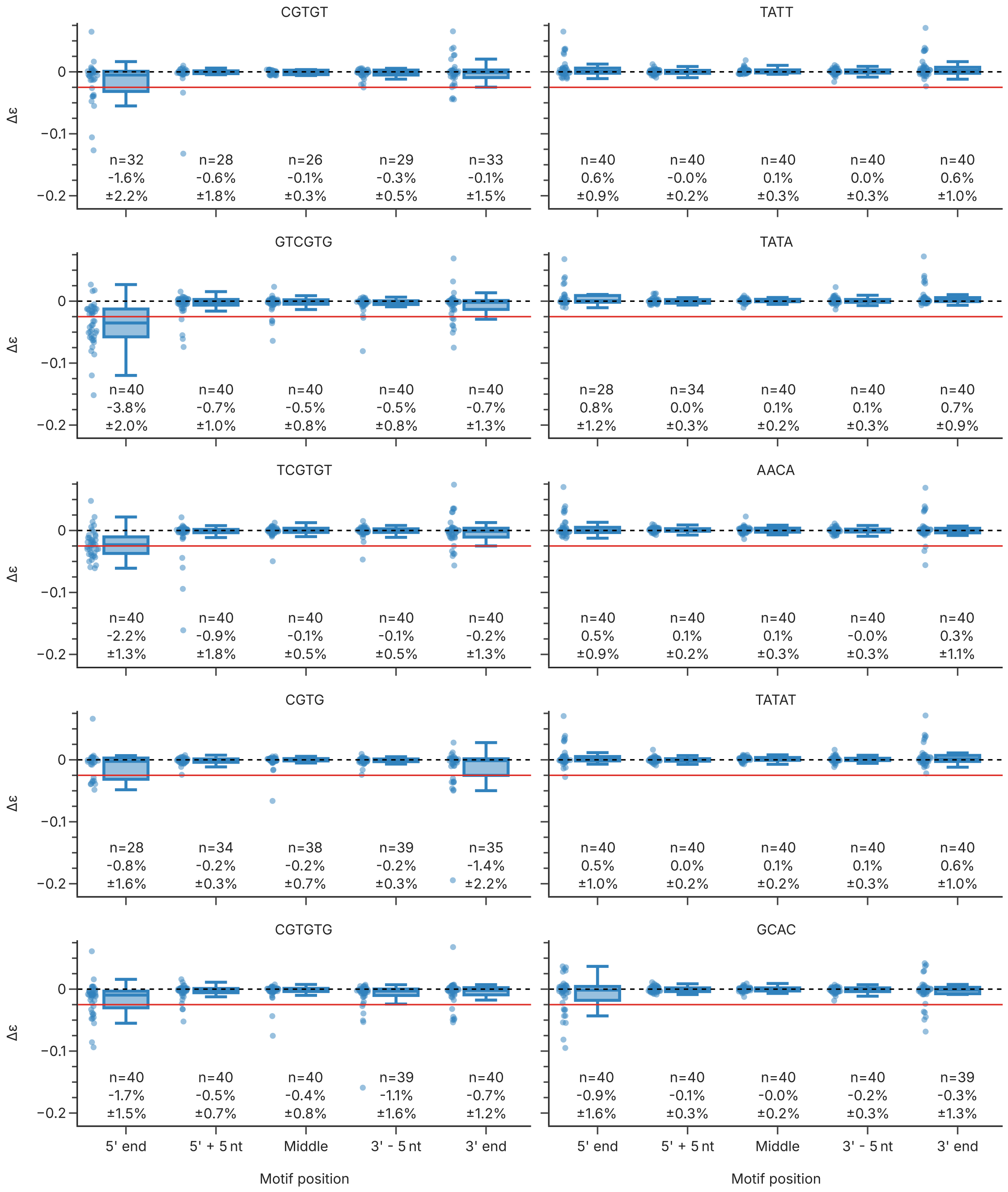


**Supplementary Fig. 10** Motif- and position-dependent change in amplification efficiencies for the workflow based on the Erlich et al. pool, in the internal repeat of the experiment. Each panel shows the difference in amplification efficiency ($\Delta\epsilon$) between the base sequence without motif and the same sequence with the corresponding motif inserted at the indicated position (x-axis). All motifs in the left column correspond to the GCall/GCfix datasets, whereas the motifs in the right column belong to the Erlich et al. dataset. The dotted line represents no change in amplification efficiency, whereas the solid red line indicates a considerable change in amplification efficiency (-2.5%, roughly the threshold corresponding to the classification as a poorly amplifying sequence in the GCall/GCfix datasets). The number of individual sequences ($n$) and the mean difference and the Bonferroni-corrected 95% confidence interval are also shown below each box.

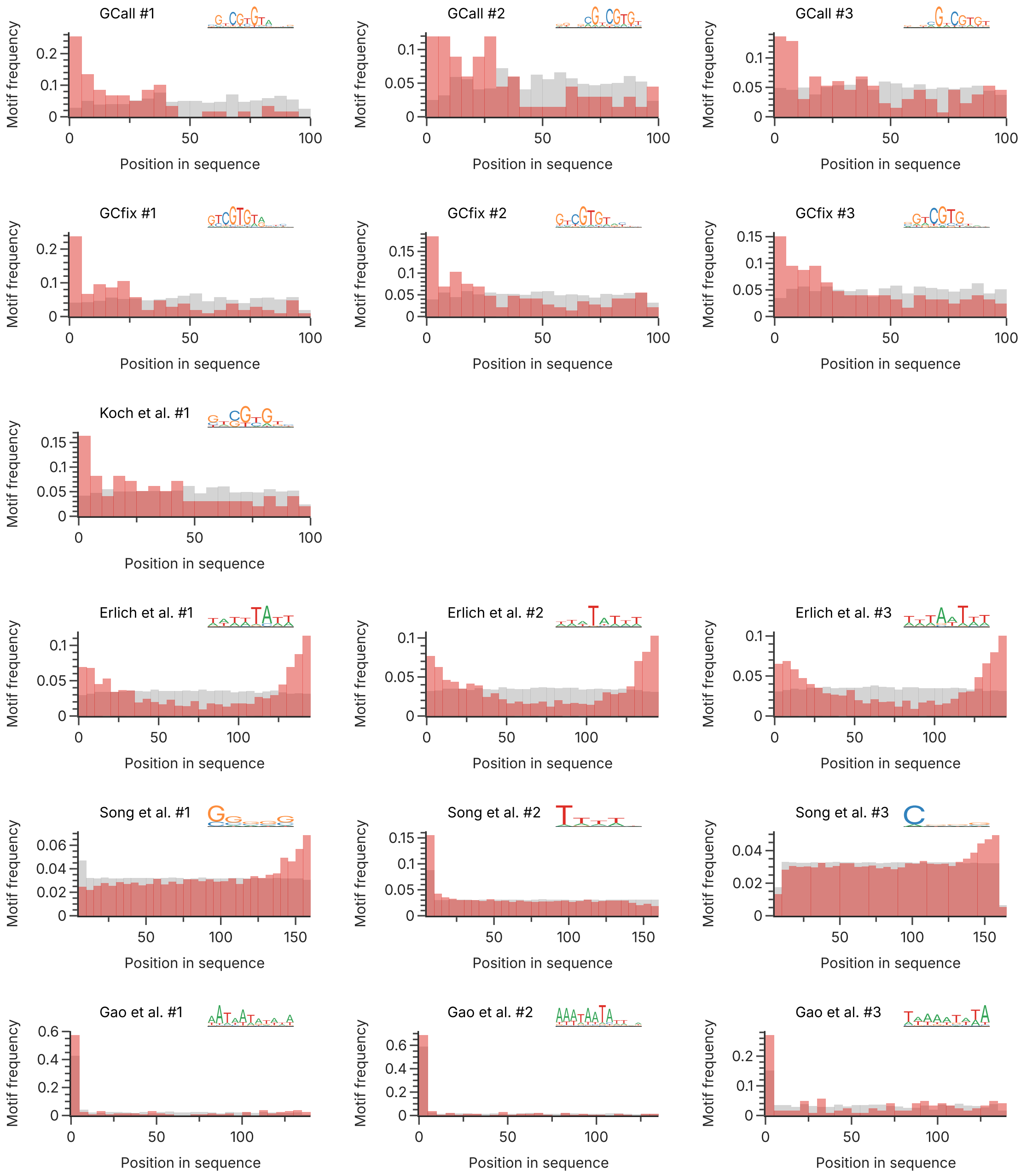


**Supplementary Fig. 11** Top motifs and their positional bias for all datasets. In each row, the (up to) top three motifs of the datasets considered in this study are shown, as well as the positional frequency of this motif in the sequences classified as poorly amplifying (red) and normal (gray). Due to the length of the motif window, the number of possible positions is lower than the total length of the sequence. The Koch et al. dataset featured only one significant motif.

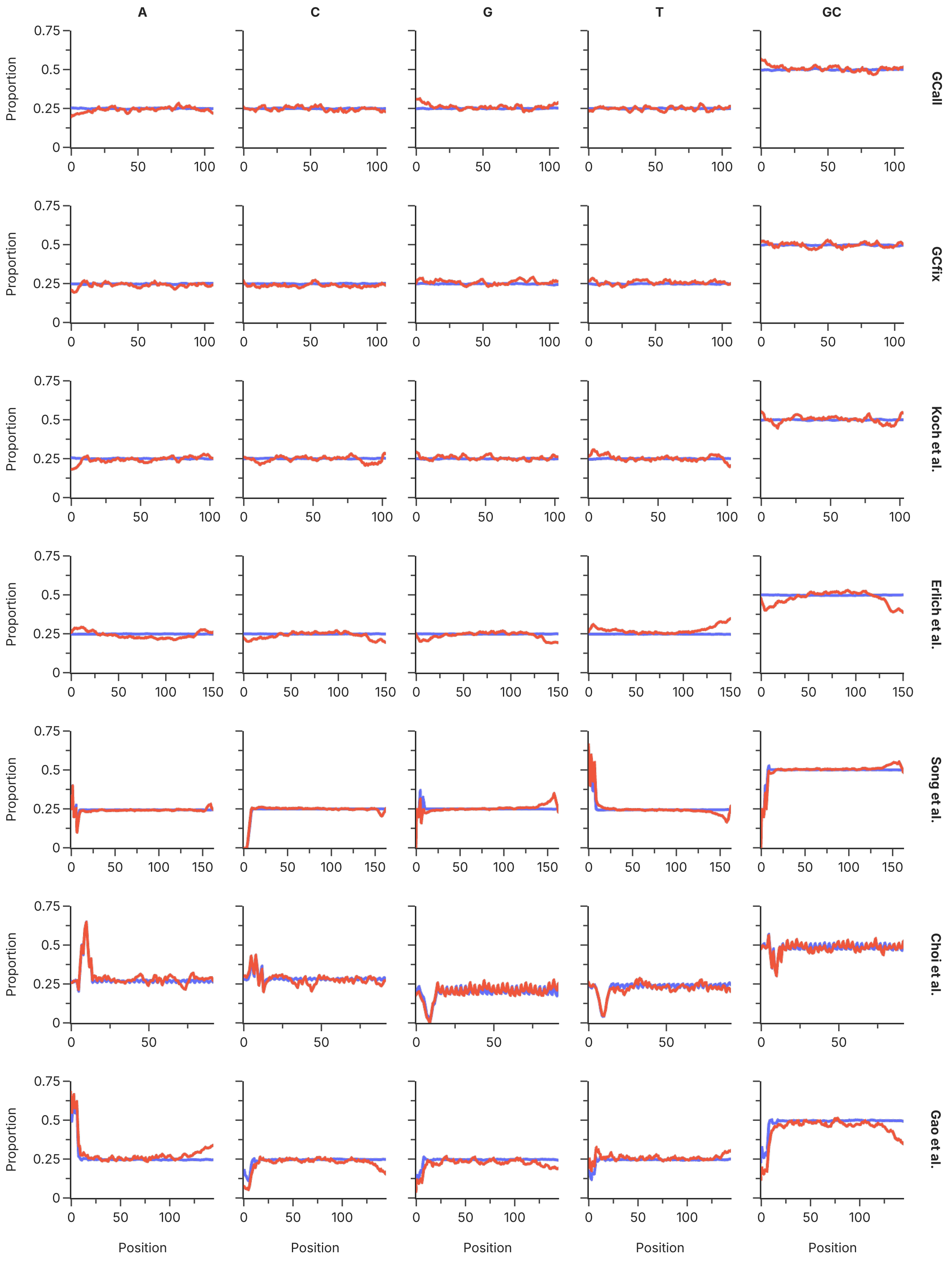


**Supplementary Fig. 12** Base composition by position in the design sequences for the datasets generated in this study and the datasets from the literature. Across all columns (specifying the fraction of A, C, G, T, and GC), each row shows only the design sequences from one dataset, separating the sequences with poor (red) and normal amplification efficiencies (blue). To reduce noise, the base composition is shown with a rolling mean, employing a 5 nt window size.

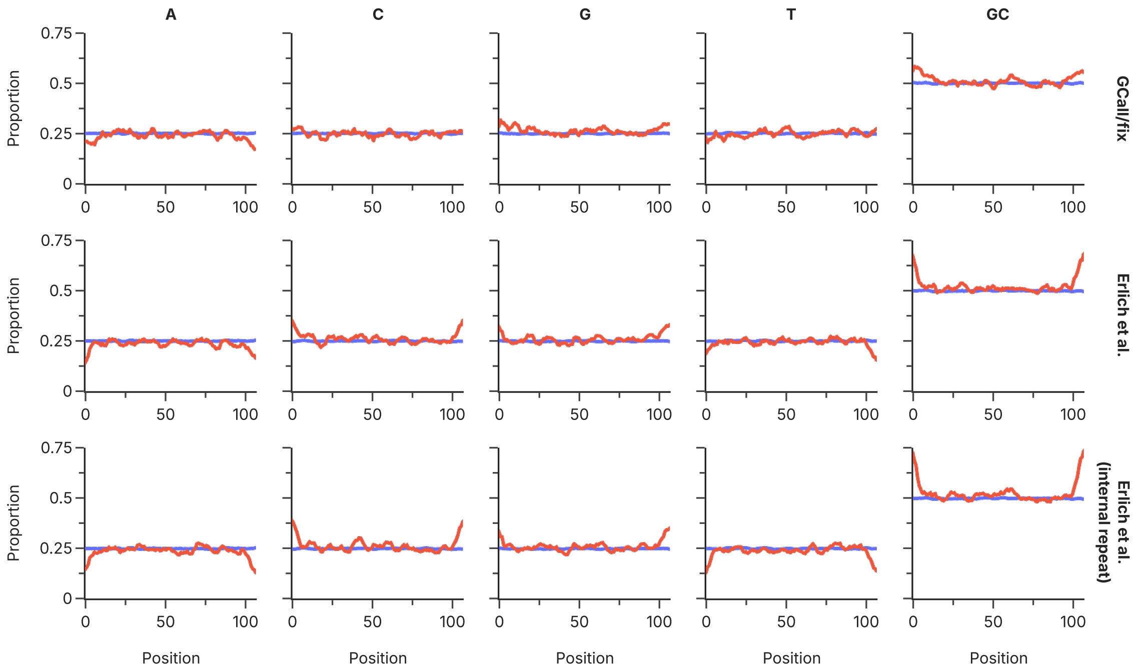


**Supplementary Fig. 13** Base composition by position in the design sequences for the external validation datasets in the different experiment conditions. Across all columns (specifying the fraction of A, C, G, T, and GC), each row shows the same set of design sequences, but each time separated by poor (red) and normal amplification efficiencies (blue). To reduce noise, the base composition is shown with a rolling mean, employing a 5 nt window size.

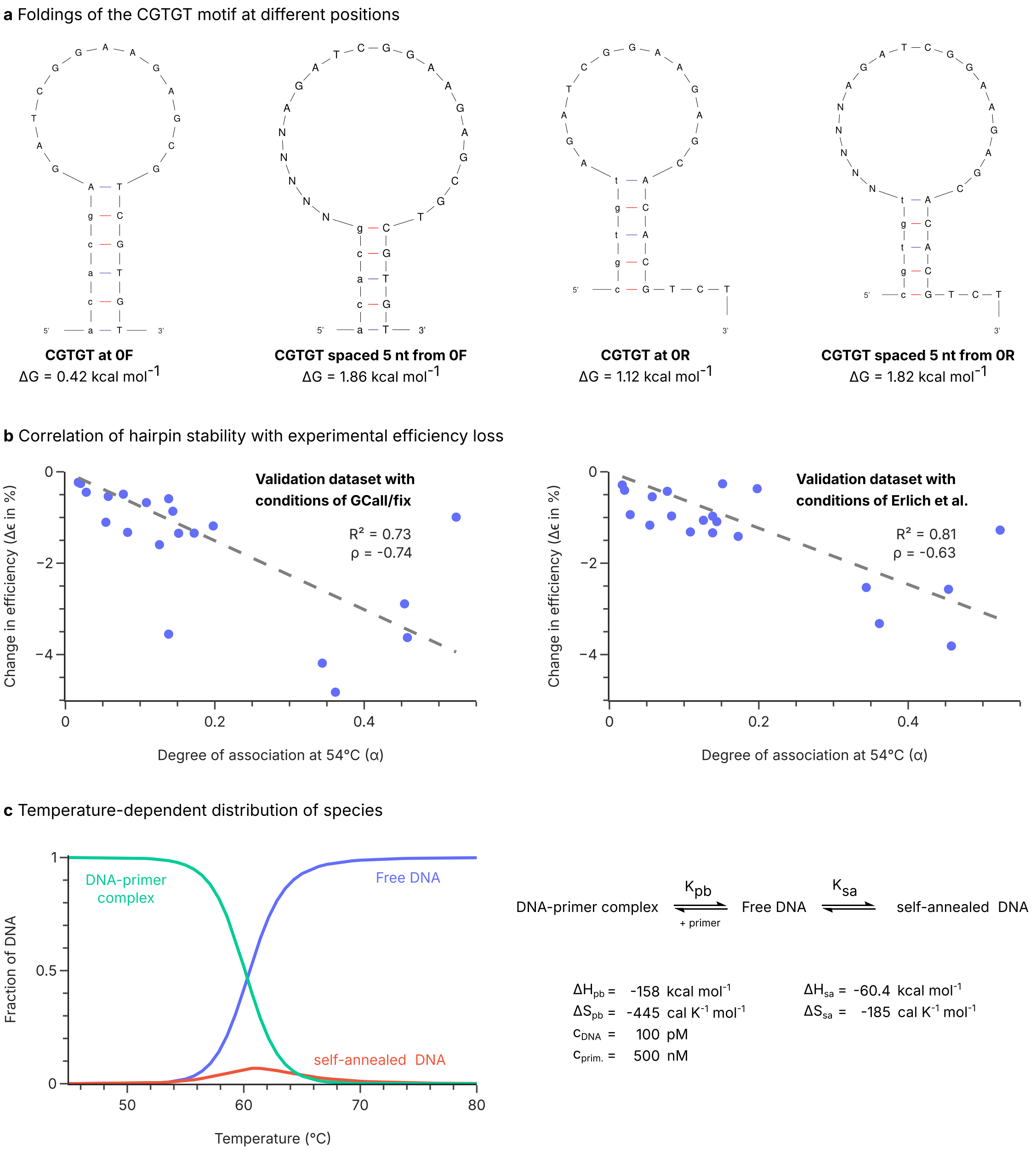


**Supplementary Fig. 14** Thermodynamic considerations for template-motif interactions. (a) Predicted foldings of the CGTGT motif (lowercase) with the 5'- and 3'-adapters (0F and 0R, respectively). To illustrate the effect of additional spacing between the motif and the end of the adapter, interactions are also shown including a 5 N spacer. (b) Correlation between the degree of association calculated from the thermodynamic parameters, and the change in amplification efficiency observed in the experiment (see Fig. 6) for each motif at each position of a design sequence. The left plot uses the experimental data using the conditions of GCall/GCfix, whereas the plot on the right uses the experimental conditions of Erlich et al. (c) Thermodynamic equilibrium between the free DNA, the DNA-primer complex, and self-annealed DNA, illustrated as an example for one motif. The distribution of free DNA (blue), the DNA-primer complex (green) and the self-annealed DNA (red) is shown as a function of temperature, assuming temperature-independent thermodynamic parameters. All thermodynamic parameters were calculated at 54 °C and 50 mM $Na^+$ using the mfold webserver.

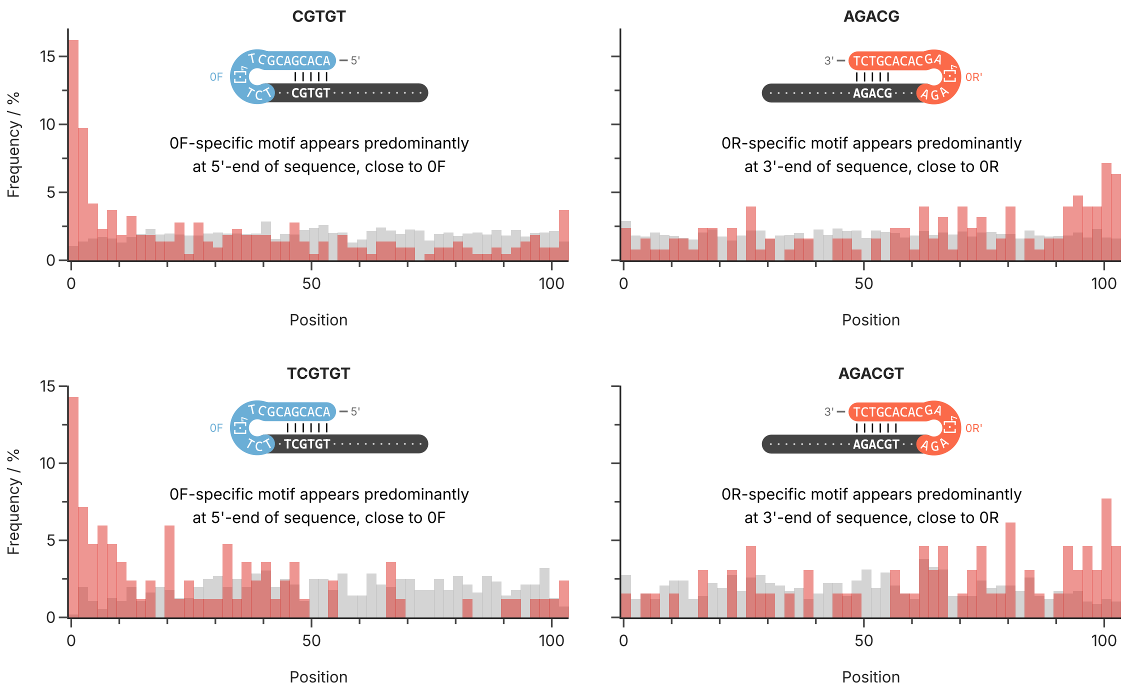


**Supplementary Fig. 15** Positional distribution of individual k-mers in the datasets of GCall and GCfix, within poorly (red) and normally amplifying sequences (gray). Histograms show the frequency in 2 nt-windows among all positions in which a k-mer occurs. Insets show the hypothesized self-annealing that occurs at the sequences' 5'- and 3'-ends.

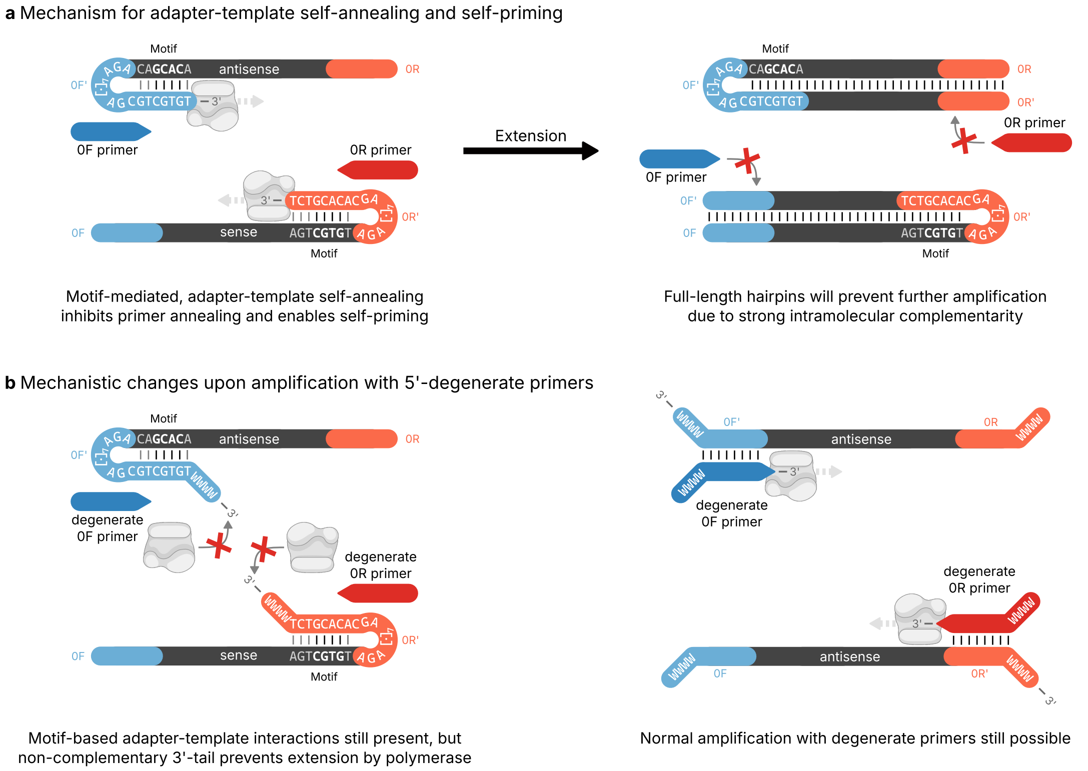


**Supplementary Fig. 16** Illustration of the proposed mechanism for self-priming. (a) Motif-mediated interactions between the adapter at the front (top) or the end (bottom) of the design sequence and the template lead to hairpin formation. Due to the free 3'-end of the adapter involved in the hairpin, annealing of the primer is inhibited, and extension of the hairpin by a polymerase is facilitated. This leads to full-length hairpins that may no longer be amplified due to their self-complementarity (right). (b) Upon amplification with 5'-degenerate primers, the hairpin formation is still possible, however, the non-complementary 3'-ends introduced into the template during amplification prevent extension of hairpins by polymerases (left). Meanwhile, normal amplification is still possible (right).

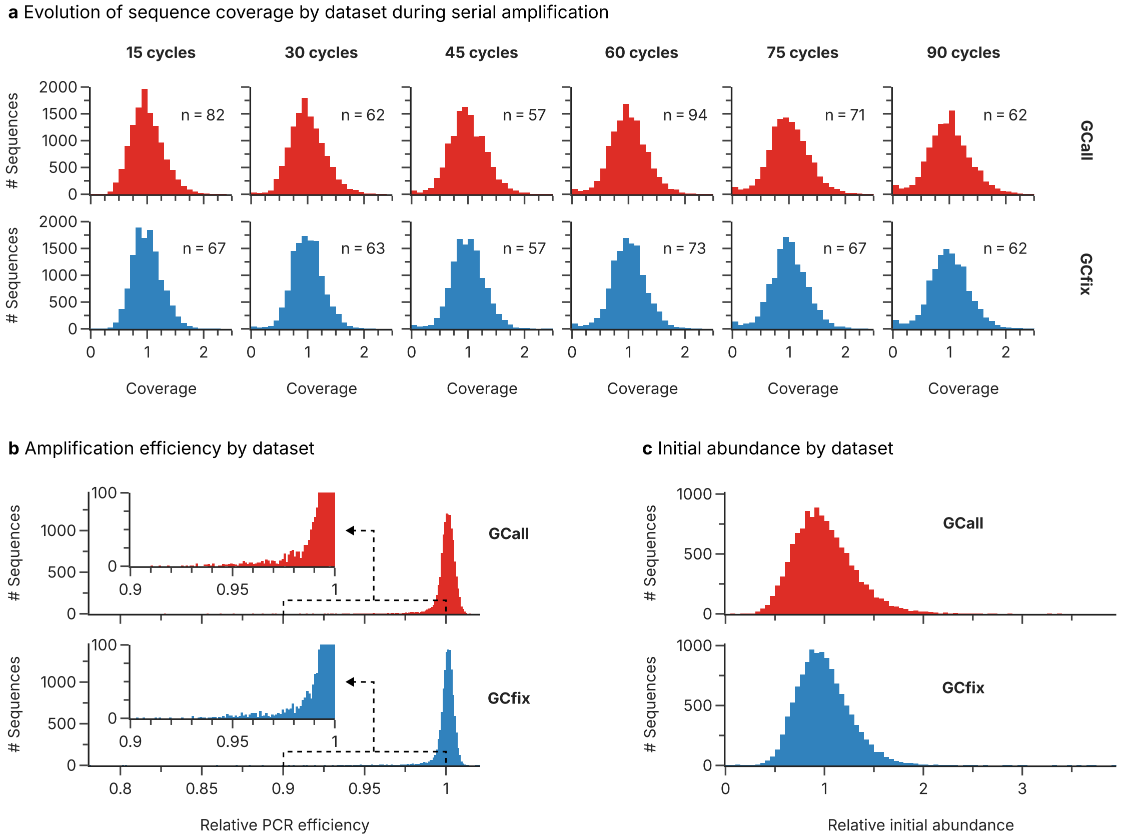


**Supplementary Fig. 17** Comparison of the GCall and GCfix pools during serial amplification. The evolution of sequence coverage during serial amplification (a) for the GCall (top, red) and GCfix (bottom, blue) pools is shown, with the mean sequencing coverage given for each sequencing endpoint ($n$). The distributions of the estimated amplification efficiency (b) and the initial abundance (c) between the GCall (top, red) and GCfix (bottom, blue) datasets are also very similar.

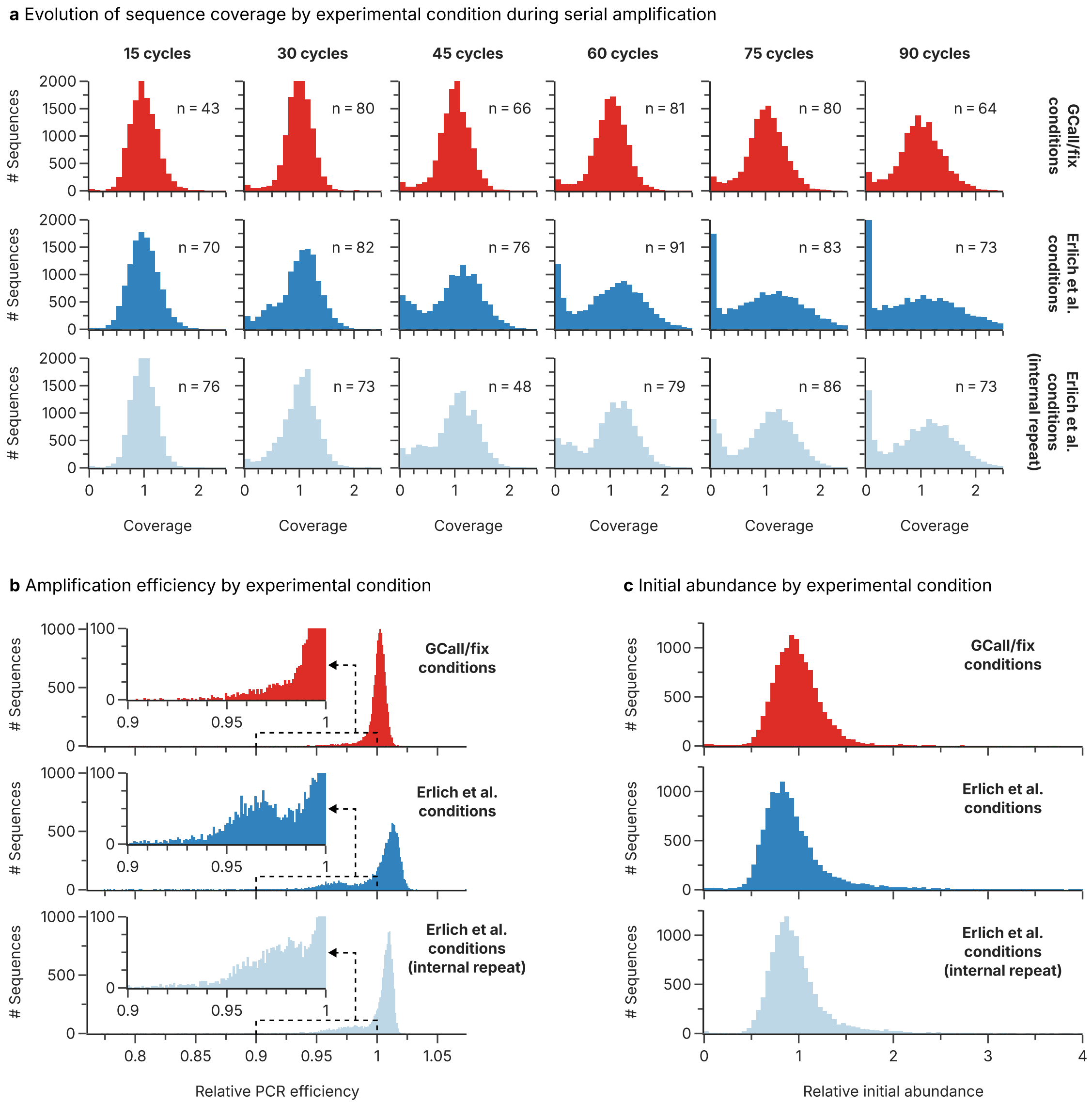


**Supplementary Fig. 18** Comparison of the validation pools during serial amplification. The evolution of sequence coverage during serial amplification (a) for the validation pool using the GCall/GCfix conditions (top, red), the Erlich et al. conditions (middle, dark blue), and the internal repeat of the Erlich et al. conditions (bottom, light blue) is shown, with the mean sequencing coverage given for each sequencing endpoint ($n$). The distributions of the estimated amplification efficiency (b) and the initial abundance (c) between the GCall/GCfix conditions (top, red), the Erlich et al. conditions (middle, dark blue), and the internal repeat of the Erlich et al. conditions (bottom, light blue) are dissimilar only for the GCall/GCfix conditions.

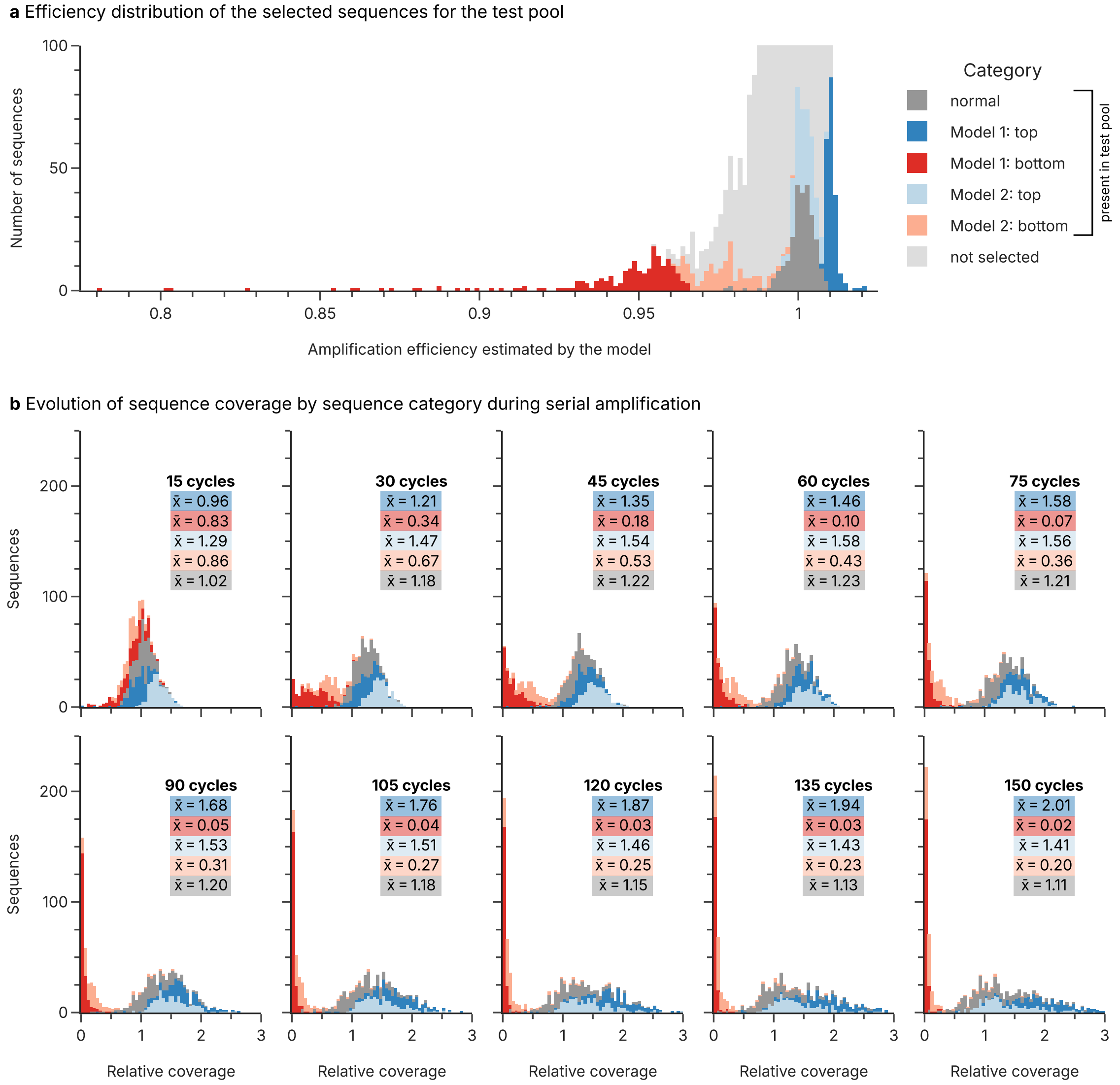


**Supplementary Fig. 19** Composition and serial amplification of the test pool. The test pool for validation of estimated amplification efficiencies was composed of selected sequences from both the GCall and GCfix pools. Selection of 1,000 sequences from all 24,000 sequences was performed based on the estimations of amplification efficiency by two models (a), see also Supplementary Note 3. The simpler model 1 considered only exponential amplification, whereas model 2 also considered dilutions and sequencing propensity. The evolution of the coverage of the selected sequences after synthesis and serial amplification (b) highlights that both models identify poorly amplifying sequences. In both figures, the sequences selected from model 1 are shown in dark colors (dark red: poorly amplifying, dark blue: well amplifying), whereas the selected sequences from model 2 are shown in light colors (light red: poorly amplifying, light blue: well amplifying). The randomly selected sequences (gray, labeled normal) reflect the evolution of an average sequence during serial amplification.

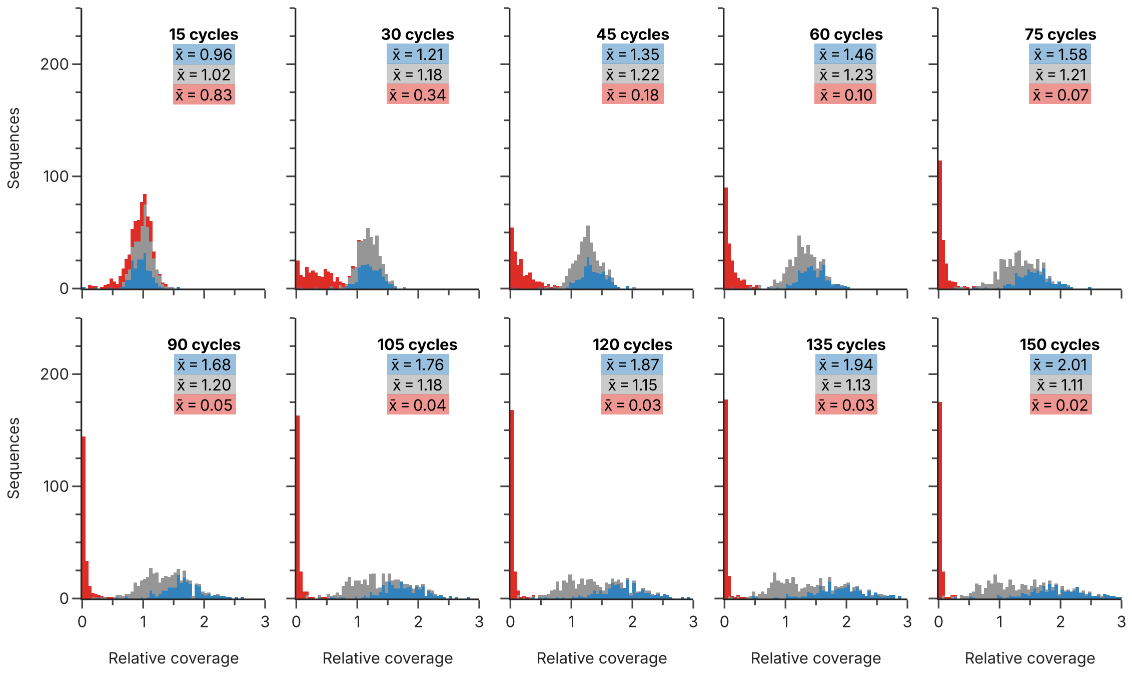


**Supplementary Fig. 20** Composition and serial amplification of the test pool. This figure is a variant of Supplementary Fig. 19b showing only the evolution of the sequences selected by model 1. The sequences identified by the model as poorly amplifying (red), or well amplifying (blue) are shown together with randomly selected sequences (gray) from the GCall and GCfix pools during serial amplification.

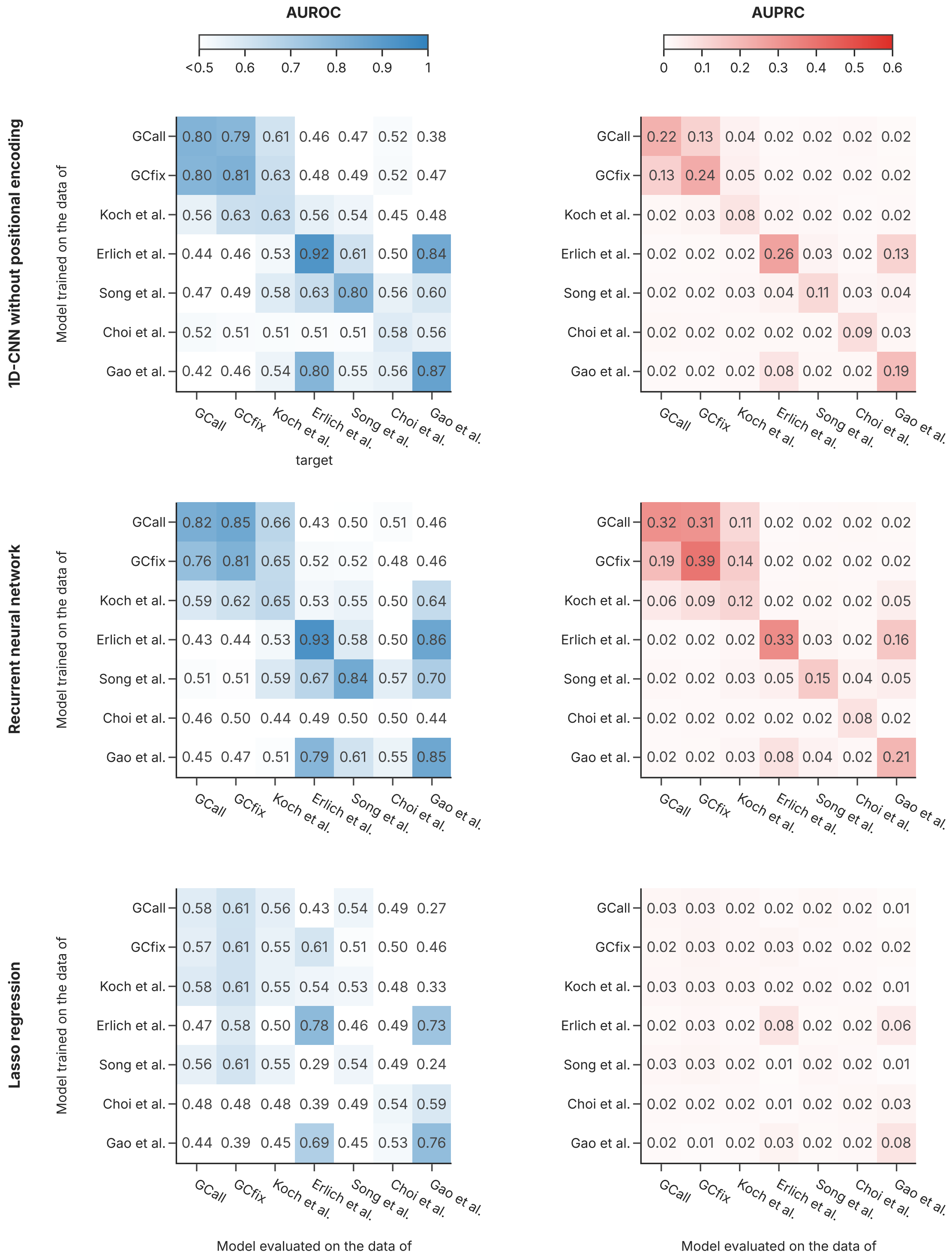


**Supplementary Fig. 21** This figure presents the AUROC and AUPRC for different baseline models, including 1D-CNN with and without positional encoding, recurrent neural network, and Lasso regularized logistic regression, trained and evaluated across various literature datasets. The models are evaluated on their ability to categorize DNA sequences into low-efficiency and normal-efficiency classes based on the empirically set 2% threshold.

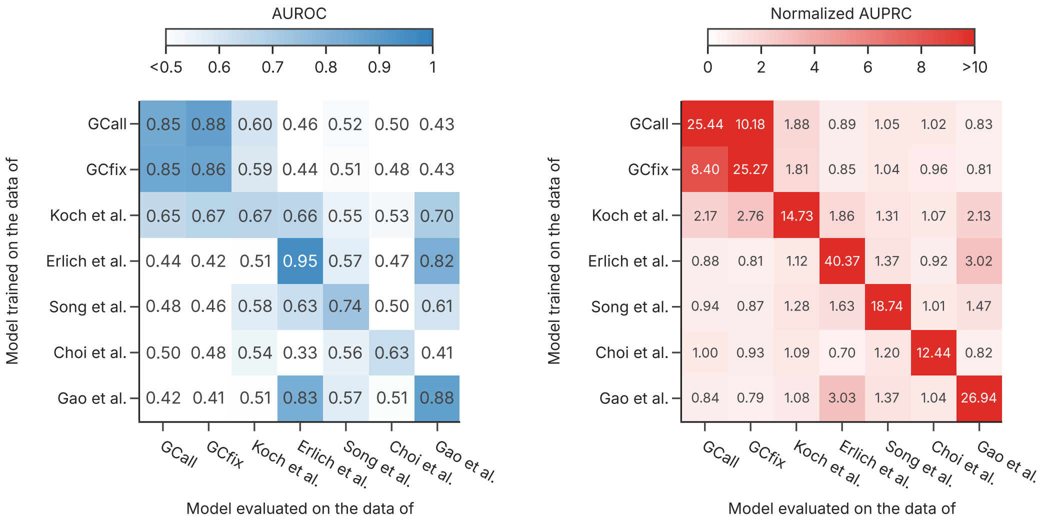


**Supplementary Fig. 22** This figure presents the AUROC and the normalized AUPRC (AUPRC over the positive class prevalence) for the proposed 1D-CNN + PE model trained and evaluated across various literature datasets. The models are evaluated on their ability to categorize DNA sequences into low-efficiency and normal-efficiency classes based on a threshold defined as 1 standard deviation below the mean PCR efficiency.

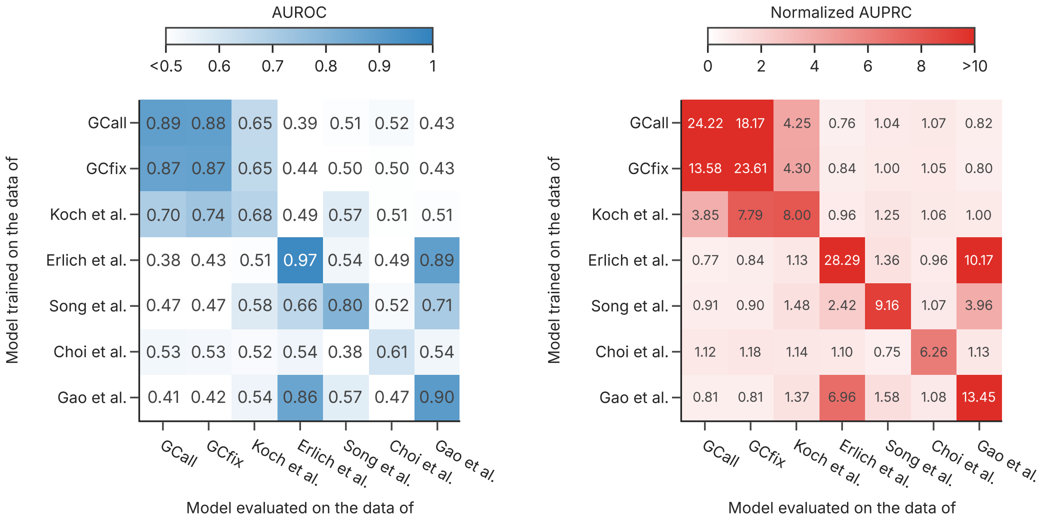


**Supplementary Fig. 23** This figure presents the AUROC and the normalized AUPRC (AUPRC over the positive class prevalence) for the proposed 1D-CNN + PE model trained and evaluated across various literature datasets. The models are evaluated on their ability to categorize DNA sequences into low-efficiency and normal-efficiency classes based on a threshold defined as 2 standard deviations below the mean PCR efficiency.

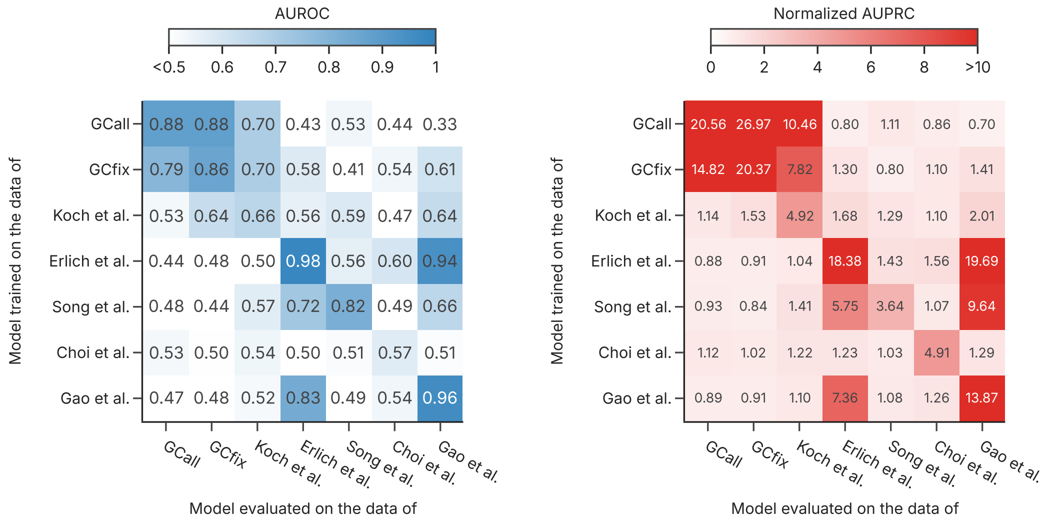


**Supplementary Fig. 24** This figure presents the AUROC and the normalized AUPRC (AUPRC over the positive class prevalence) for the proposed 1D-CNN + PE model trained and evaluated across various literature datasets. The models are evaluated on their ability to categorize DNA sequences into low-efficiency and normal-efficiency classes based on a threshold defined as 3 standard deviations below the mean PCR efficiency.

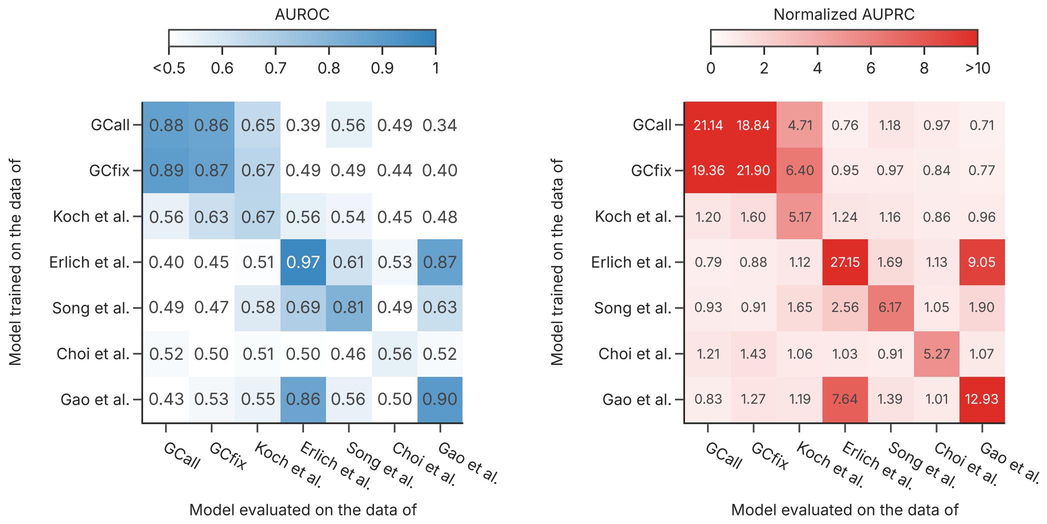


**Supplementary Fig. 25** This figure presents the AUROC and the normalized AUPRC (AUPRC over the positive class prevalence) for the proposed 1D-CNN + PE model trained and evaluated across various literature datasets. The models are evaluated on their ability to categorize DNA sequences into low-efficiency and normal-efficiency classes based on a threshold defined as 2% of the total sequences.

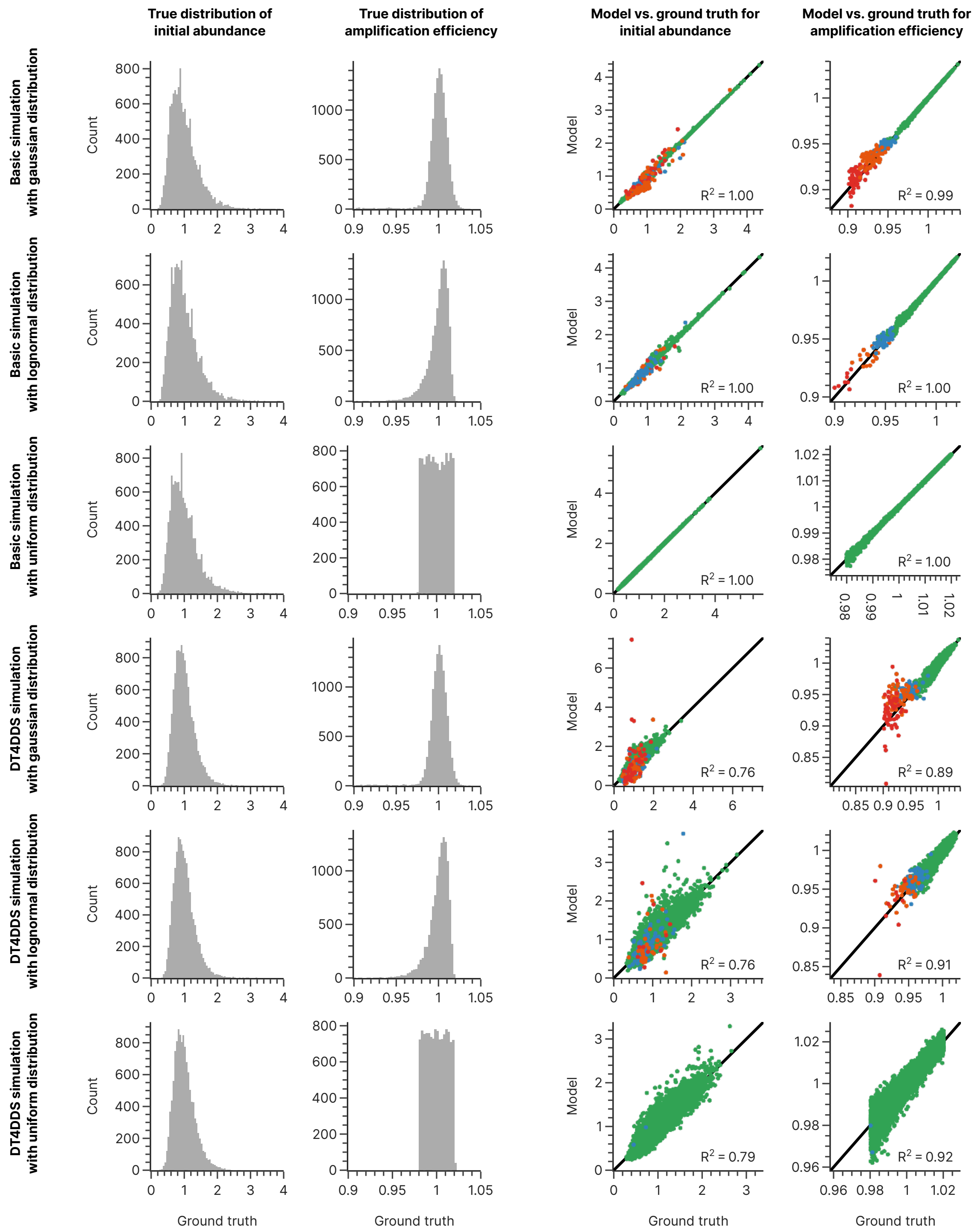


**Supplementary Fig. 26** Verification of the two-parameter fitting of the exponential PCR equation using in-silico simulations. Sequencing data with varying distributions of the true amplification efficiency and the true initial coverage bias were generated using a basic model and the DT4DDS software (left column, see Methods). The results of the parameter estimation using the exponential PCR equation were compared to the ground truth parameters used for simulation (right column), and show good agreement across all tested parameter distributions. The simulated datasets also consisted of six experimental endpoints at different cycle counts, identical to the experimental datasets. The color of the points in the right column corresponds to the frequency with which the sequence was not observed in the six experimental endpoints: never (i.e., always present, green), once (blue), twice (orange), and at least three times (red).

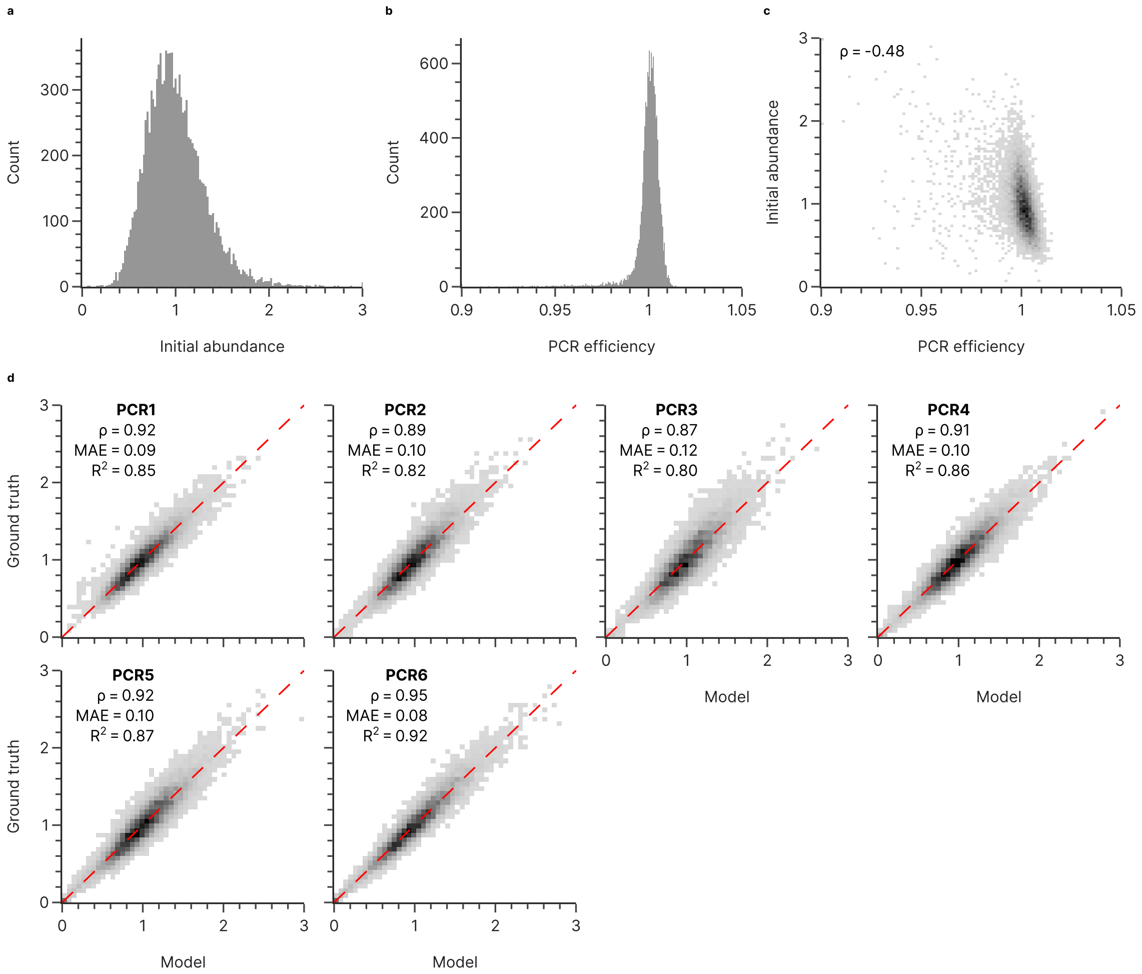


**Supplementary Fig. 27** Results and performance of the two-parameter fitting for the GCall dataset. The distributions of the fitted parameters of the initial abundance (a) and the amplification efficiency (b) are shown, as well as the correlation between both parameters (c). For each sequencing endpoint, the plots in (d) show the agreement between the experimental abundance (Ground truth, y-axis) and the expected abundance based on the PCR model and the fitted parameters (Model, x-axis). Panels (c) and (d) show relative density, and (d) includes the diagonal of perfect agreement (red, dashed line), as well as corresponding metrics (Spearman's rank correlation coefficient $\rho$, mean absolute error MAE, and coefficient of determination $R^2$). In panels (a-c), the values exceeding the axis have been clipped to the axis limits for visual clarity.

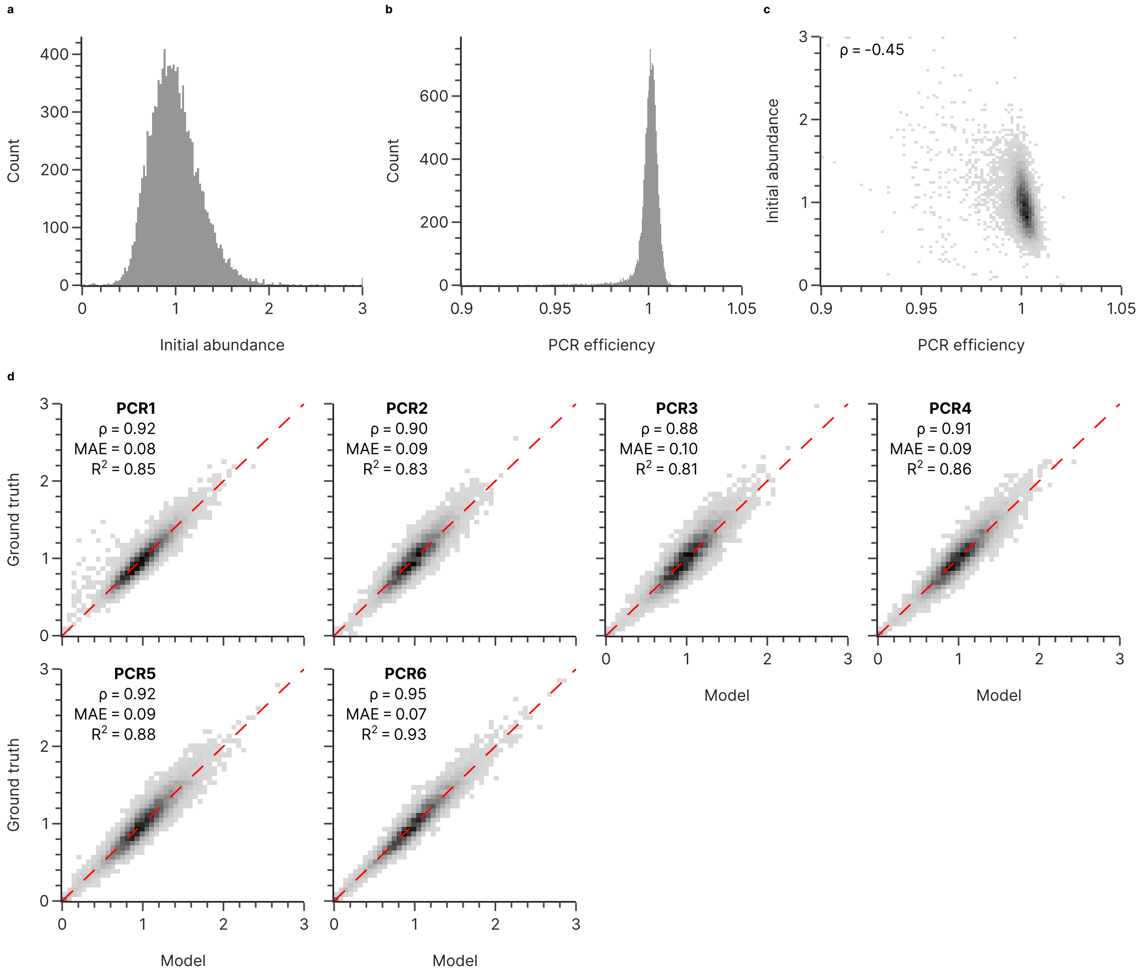


**Supplementary Fig. 28** Results and performance of the two-parameter fitting for the GCfix dataset. The distributions of the fitted parameters of the initial abundance (a) and the amplification efficiency (b) are shown, as well as the correlation between both parameters (c). For each sequencing endpoint, the plots in (d) show the agreement between the experimental abundance (Ground truth, y-axis) and the expected abundance based on the PCR model and the fitted parameters (Model, x-axis). Panels (c) and (d) show relative density, and (d) includes the diagonal of perfect agreement (red, dashed line), as well as corresponding metrics (Spearman's rank correlation coefficient $\rho$, mean absolute error MAE, and coefficient of determination $R^2$). In panels (a-c), the values exceeding the axis have been clipped to the axis limits for visual clarity.

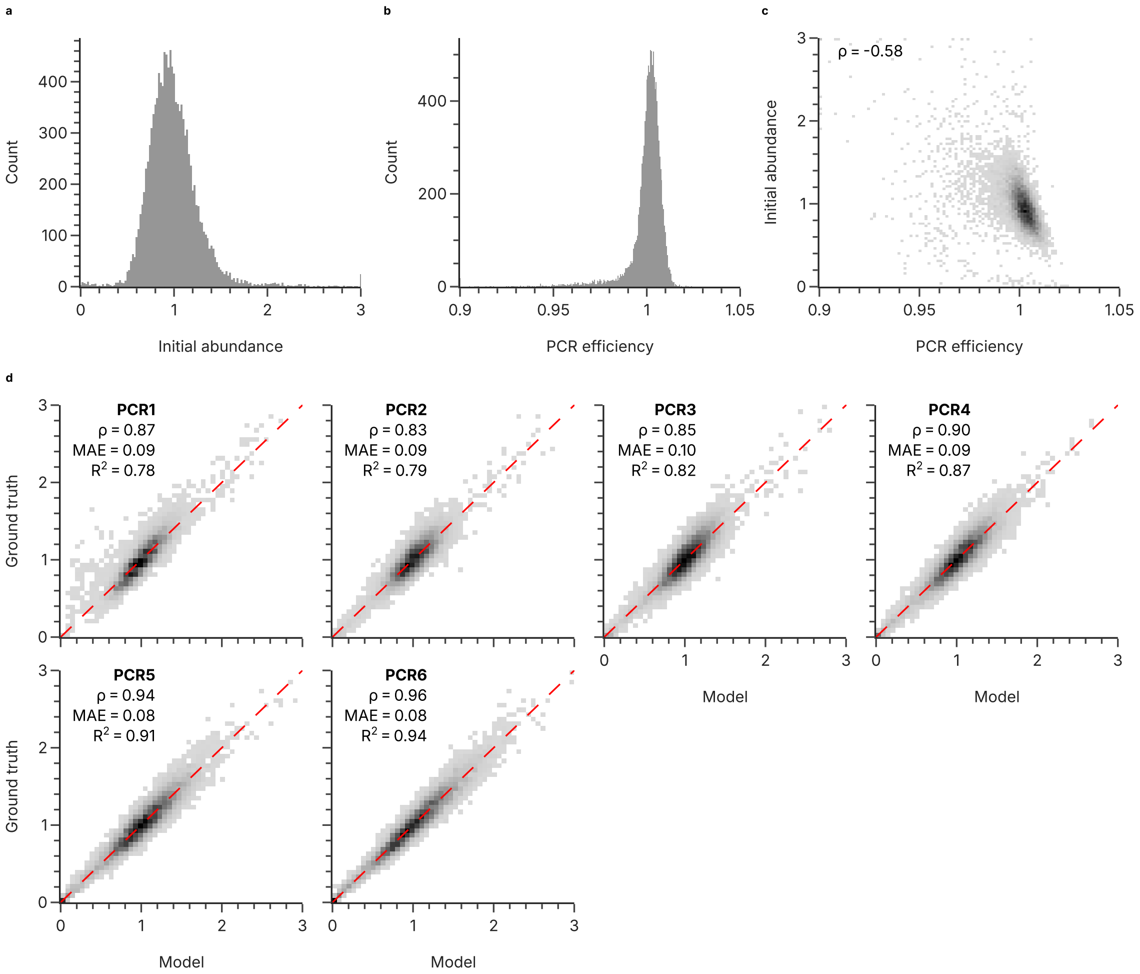


**Supplementary Fig. 29** Results and performance of the two-parameter fitting for the validation dataset using the conditions of GCall/GCfix. The distributions of the fitted parameters of the initial abundance (a) and the amplification efficiency (b) are shown, as well as the correlation between both parameters (c). For each sequencing endpoint, the plots in (d) show the agreement between the experimental abundance (Ground truth, y-axis) and the expected abundance based on the PCR model and the fitted parameters (Model, x-axis). Panels (c) and (d) show relative density, and (d) includes the diagonal of perfect agreement (red, dashed line), as well as corresponding metrics (Spearman's rank correlation coefficient $\rho$, mean absolute error MAE, and coefficient of determination $R^2$). In panels (a-c), the values exceeding the axis have been clipped to the axis limits for visual clarity.

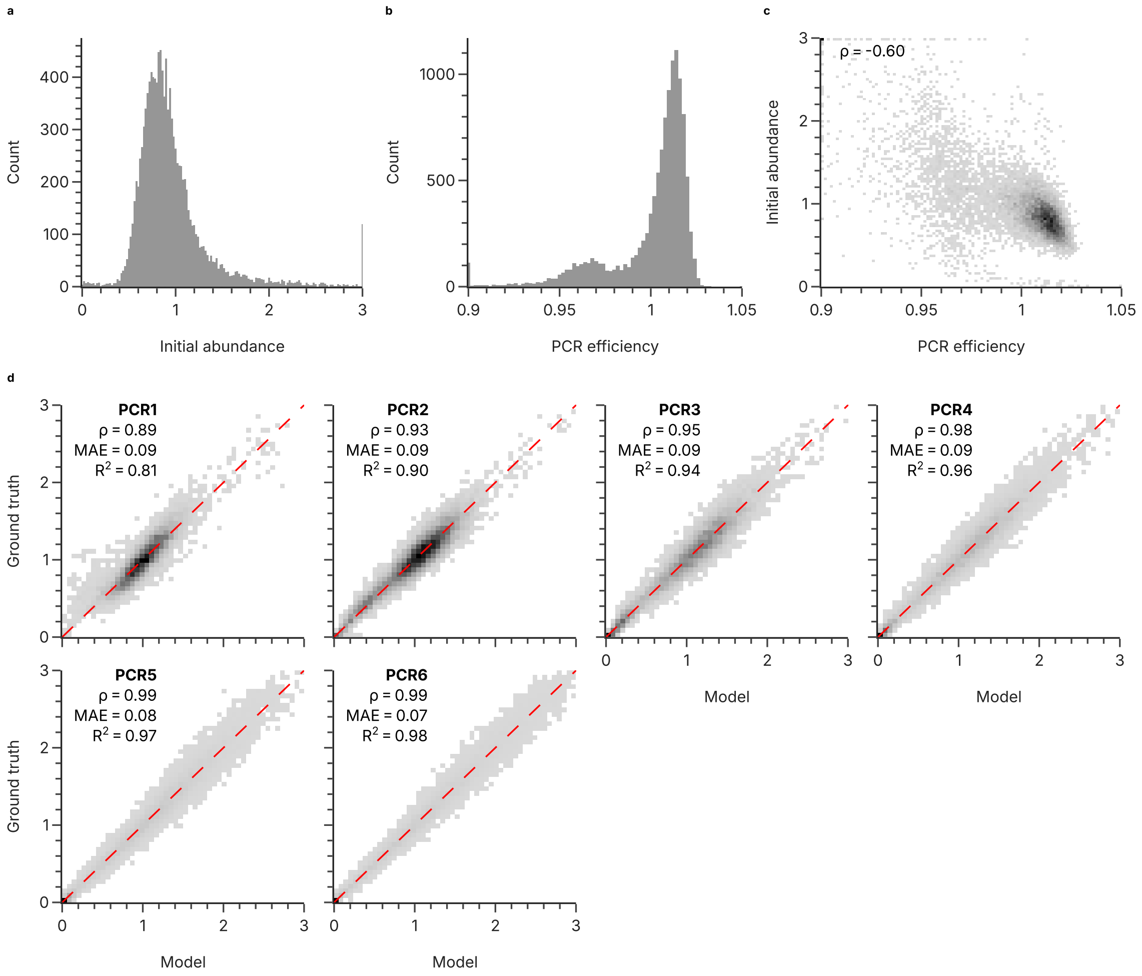


**Supplementary Fig. 30** Results and performance of the two-parameter fitting for the validation dataset using the conditions of Erlich et al. The distributions of the fitted parameters of the initial abundance (a) and the amplification efficiency (b) are shown, as well as the correlation between both parameters (c). For each sequencing endpoint, the plots in (d) show the agreement between the experimental abundance (Ground truth, y-axis) and the expected abundance based on the PCR model and the fitted parameters (Model, x-axis). Panels (c) and (d) show relative density, and (d) includes the diagonal of perfect agreement (red, dashed line), as well as corresponding metrics (Spearman's rank correlation coefficient $\rho$, mean absolute error MAE, and coefficient of determination $R^2$). In panels (a-c), the values exceeding the axis have been clipped to the axis limits for visual clarity.

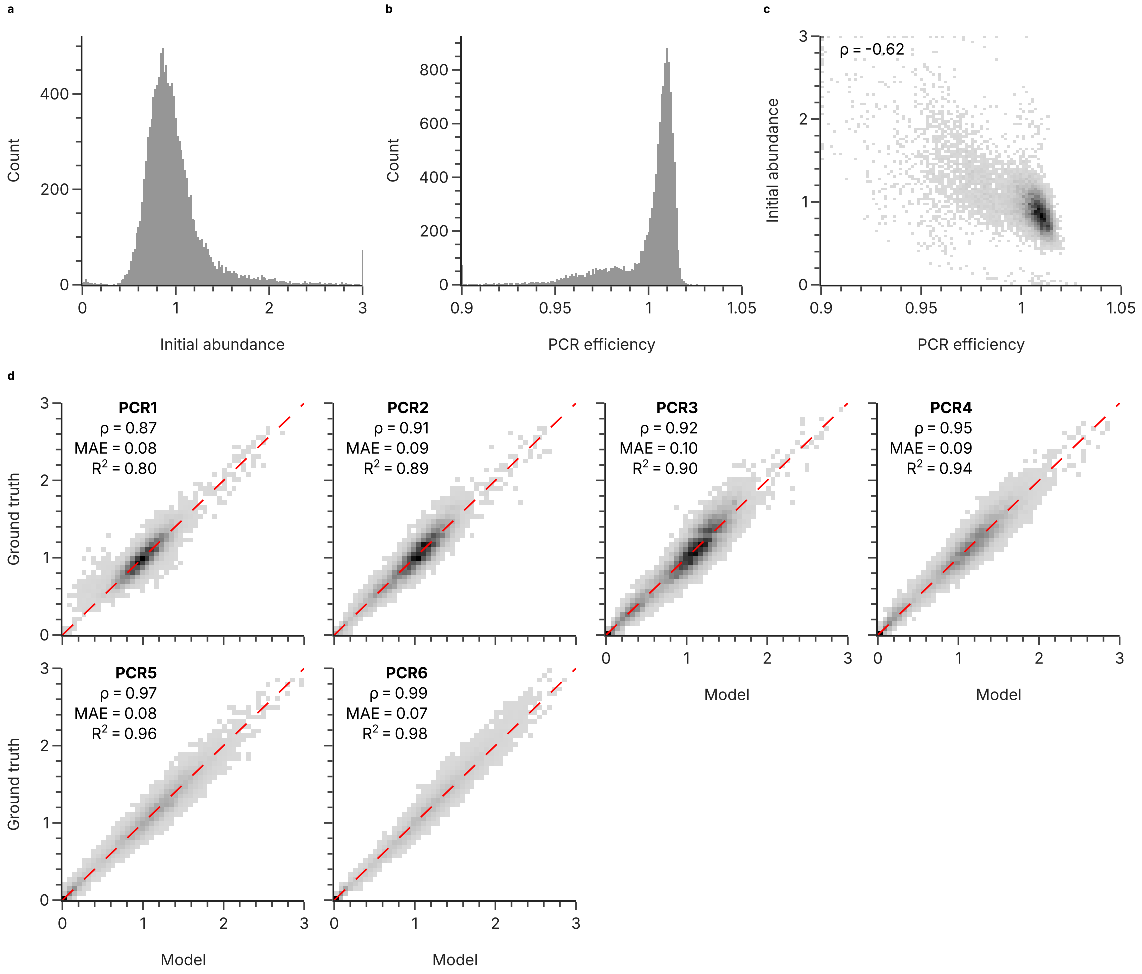


**Supplementary Fig. 31** Results and performance of the two-parameter fitting for the validation dataset using the conditions of Erlich et al., during an internal repeat of the experiment. The distributions of the fitted parameters of the initial abundance (a) and the amplification efficiency (b) are shown, as well as the correlation between both parameters (c). For each sequencing endpoint, the plots in (d) show the agreement between the experimental abundance (Ground truth, y-axis) and the expected abundance based on the PCR model and the fitted parameters (Model, x-axis). Panels (c) and (d) show relative density, and (d) includes the diagonal of perfect agreement (red, dashed line), as well as corresponding metrics (Spearman's rank correlation coefficient $\rho$, mean absolute error MAE, and coefficient of determination $R^2$). In panels (a-c), the values exceeding the axis have been clipped to the axis limits for visual clarity.

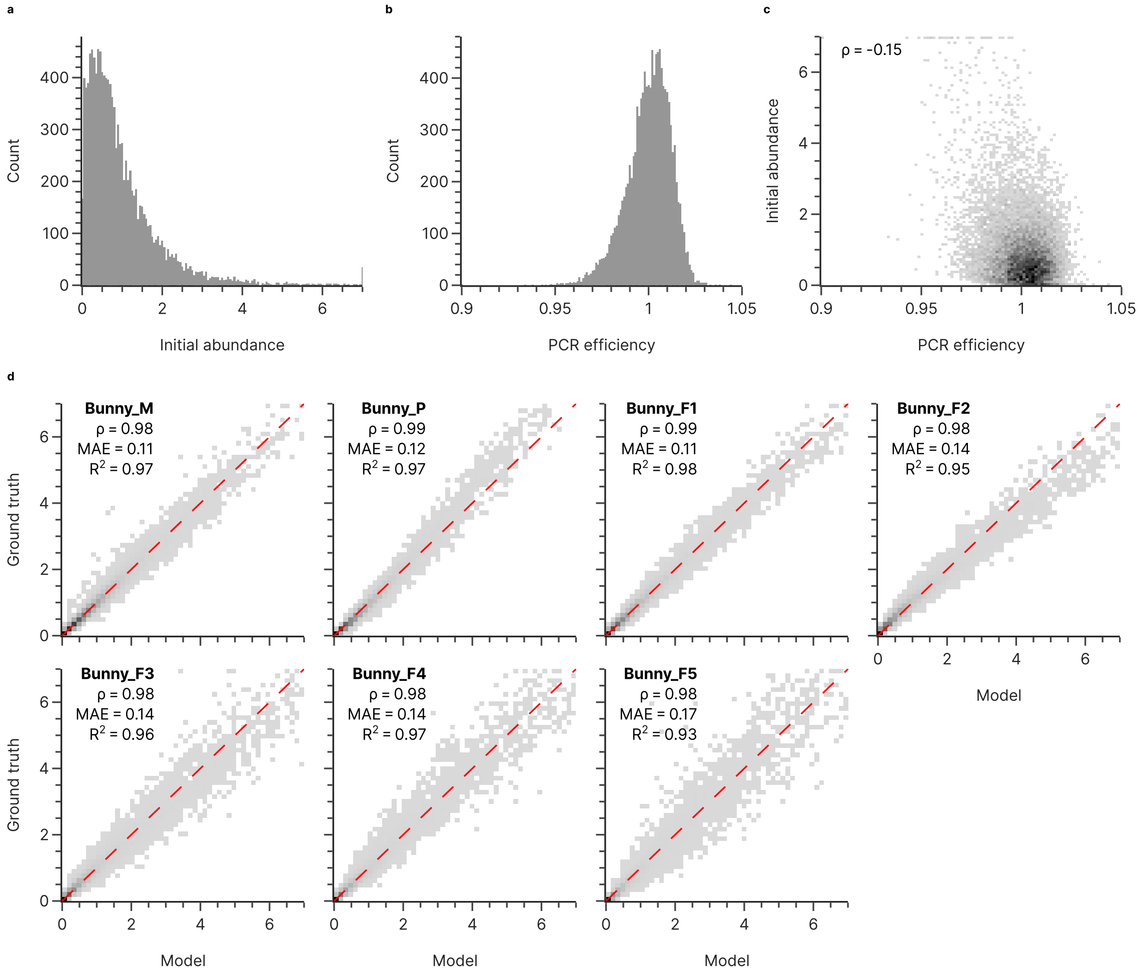


**Supplementary Fig. 32** Results and performance of the two-parameter fitting for the Koch et al. dataset. The distributions of the fitted parameters of the initial abundance (a) and the amplification efficiency (b) are shown, as well as the correlation between both parameters (c). For each sequencing endpoint, the plots in (d) show the agreement between the experimental abundance (Ground truth, y-axis) and the expected abundance based on the PCR model and the fitted parameters (Model, x-axis). Panels (c) and (d) show relative density, and (d) includes the diagonal of perfect agreement (red, dashed line), as well as corresponding metrics (Spearman's rank correlation coefficient $\rho$, mean absolute error MAE, and coefficient of determination $R^2$). In panels (a-c), the values exceeding the axis limits have been clipped to the axis limits for visual clarity.

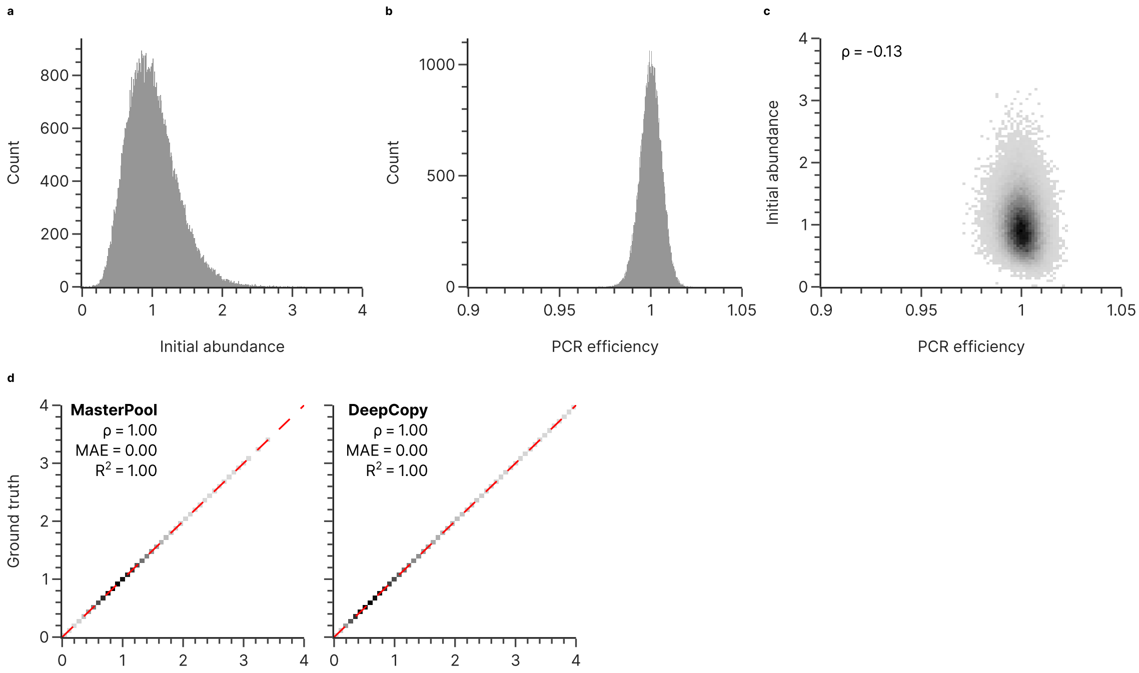


**Supplementary Fig. 33** Results and performance of the two-parameter fitting for the Erlich et al. dataset. The distributions of the fitted parameters of the initial abundance (a) and the amplification efficiency (b) are shown, as well as the correlation between both parameters (c). For each sequencing endpoint, the plots in (d) show the agreement between the experimental abundance (Ground truth, y-axis) and the expected abundance based on the PCR model and the fitted parameters (Model, x-axis). Panels (c) and (d) show relative density, and (d) includes the diagonal of perfect agreement (red, dashed line), as well as corresponding metrics (Spearman's rank correlation coefficient $\rho$, mean absolute error MAE, and coefficient of determination $R^2$). In panels (a-c), the values exceeding the axis have been clipped to the axis limits for visual clarity.

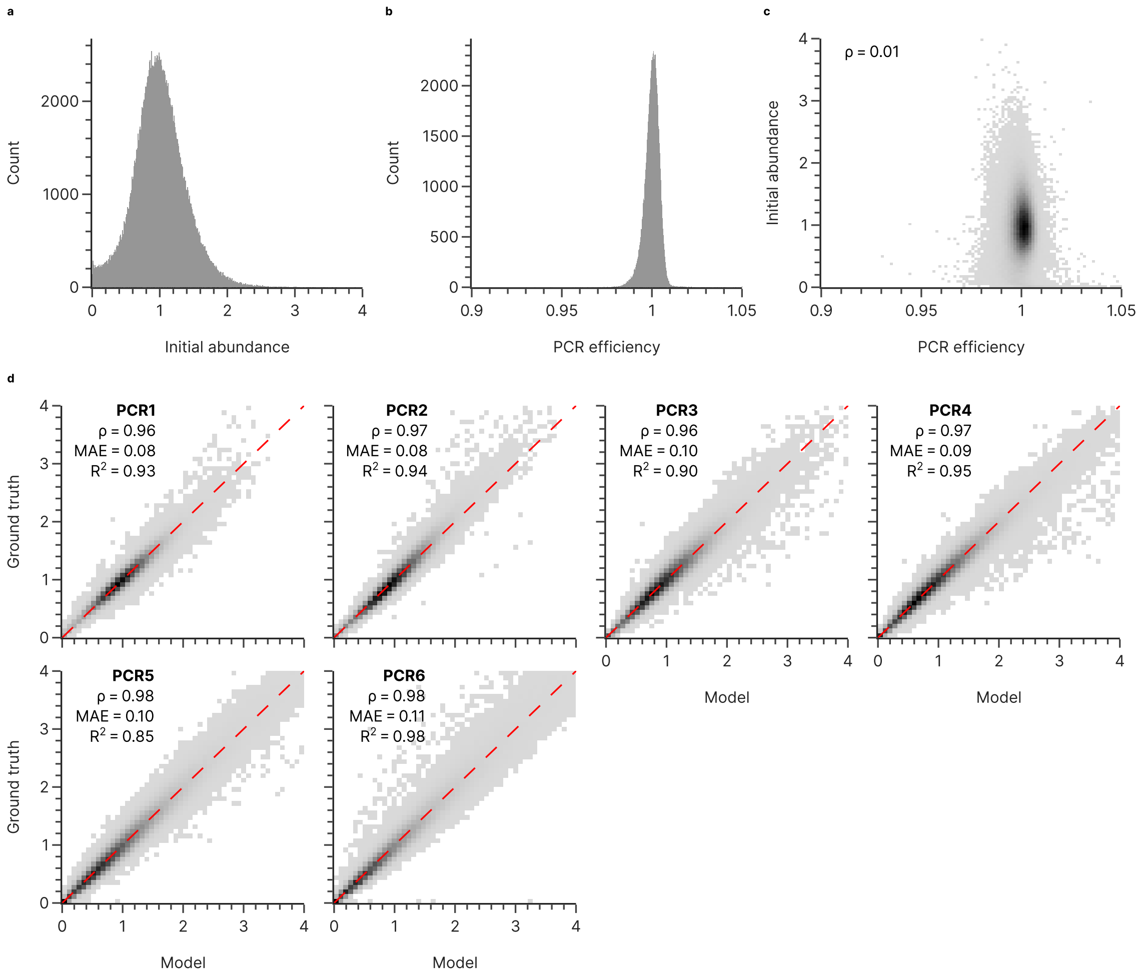


**Supplementary Fig. 34** Results and performance of the two-parameter fitting for the Song et al. dataset. The distributions of the fitted parameters of the initial abundance (a) and the amplification efficiency (b) are shown, as well as the correlation between both parameters (c). For each sequencing endpoint, the plots in (d) show the agreement between the experimental abundance (Ground truth, y-axis) and the expected abundance based on the PCR model and the fitted parameters (Model, x-axis). Panels (c) and (d) show relative density, and (d) includes the diagonal of perfect agreement (red, dashed line), as well as corresponding metrics (Spearman's rank correlation coefficient $\rho$, mean absolute error MAE, and coefficient of determination $R^2$). In panels (a-c), the values exceeding the axis have been clipped to the axis limits for visual clarity.

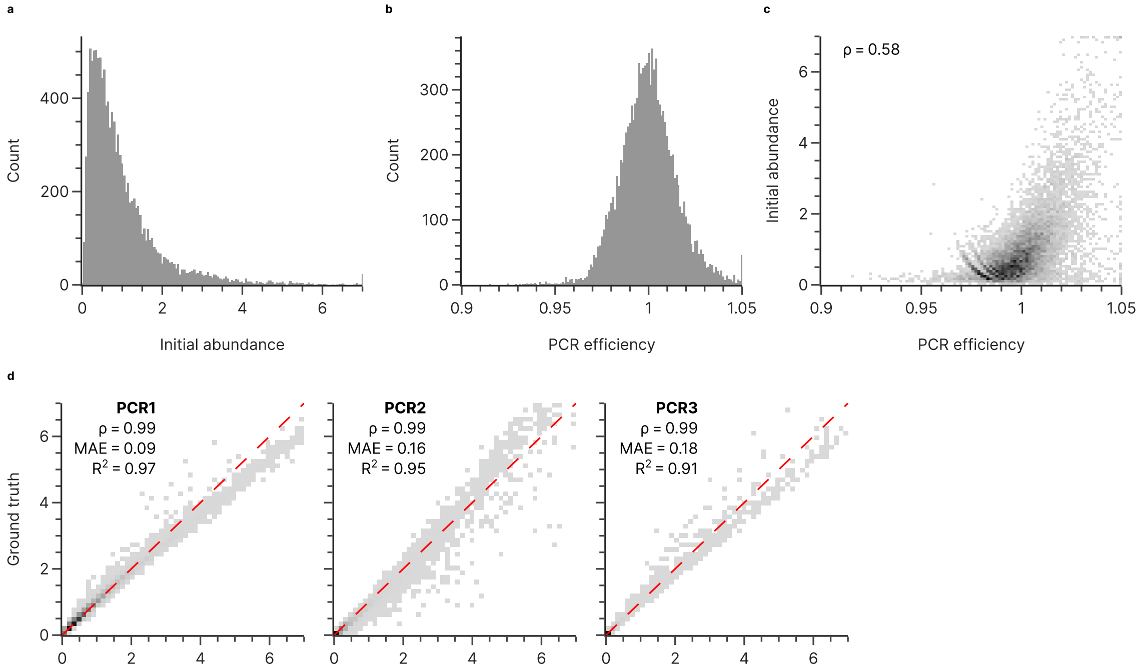


**Supplementary Fig. 35** Results and performance of the two-parameter fitting for the Gao et al. dataset. The distributions of the fitted parameters of the initial abundance (a) and the amplification efficiency (b) are shown, as well as the correlation between both parameters (c). For each sequencing endpoint, the plots in (d) show the agreement between the experimental abundance (Ground truth, y-axis) and the expected abundance based on the PCR model and the fitted parameters (Model, x-axis). Panels (c) and (d) show relative density, and (d) includes the diagonal of perfect agreement (red, dashed line), as well as corresponding metrics (Spearman's rank correlation coefficient $\rho$, mean absolute error MAE, and coefficient of determination $R^2$). In panels (a-c), the values exceeding the axis have been clipped to the axis limits for visual clarity.

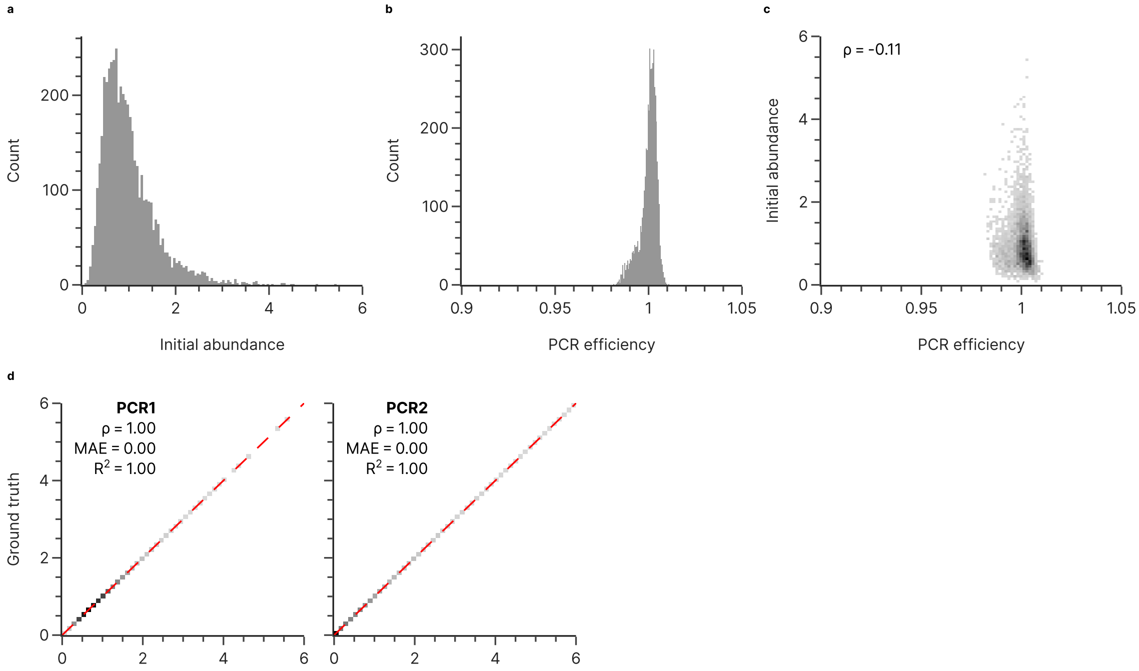


**Supplementary Fig. 36** Results and performance of the two-parameter fitting for the Choi et al. dataset. The distributions of the fitted parameters of the initial abundance (a) and the amplification efficiency (b) are shown, as well as the correlation between both parameters (c). For each sequencing endpoint, the plots in (d) show the agreement between the experimental abundance (Ground truth, y-axis) and the expected abundance based on the PCR model and the fitted parameters (Model, x-axis). Panels (c) and (d) show relative density, and (d) includes the diagonal of perfect agreement (red, dashed line), as well as corresponding metrics (Spearman's rank correlation coefficient $\rho$, mean absolute error MAE, and coefficient of determination $R^2$). In panels (a-c), the values exceeding the axis have been clipped to the axis limits for visual clarity.

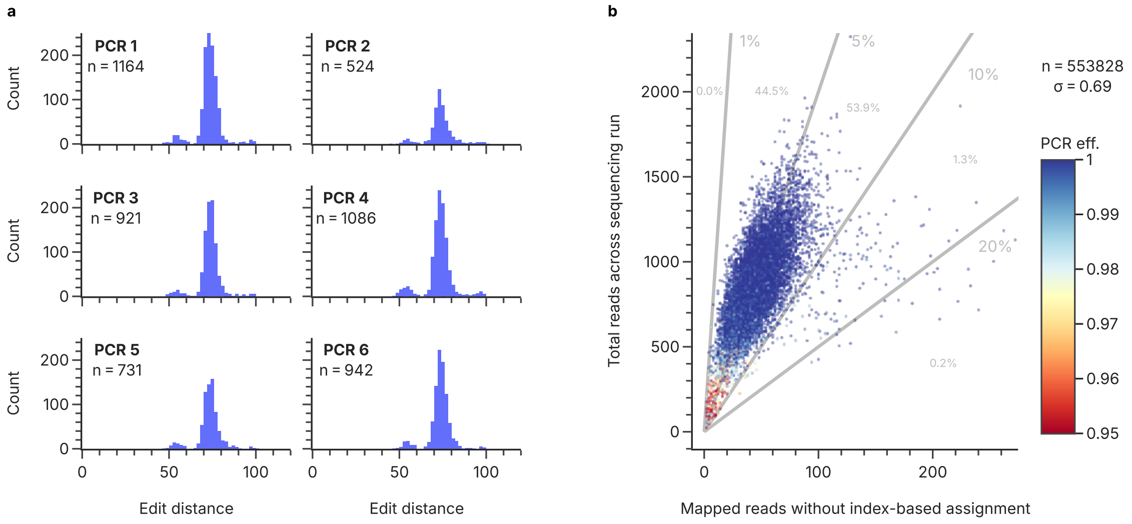


**Supplementary Fig. 37** Occurrence of sequences of the GCall pool during demultiplexing and post-processing of the sequencing data. The distribution of minimum edit distances of all sequences removed during post-processing to the design sequences (a) shows that no real reads are lost during adapter removal and mapping. The occurrence of design sequences without experimental assignment after demultiplexing (b) shows that all sequences are found in the unassigned data at approximately the same rate, proportional to their overall presence.

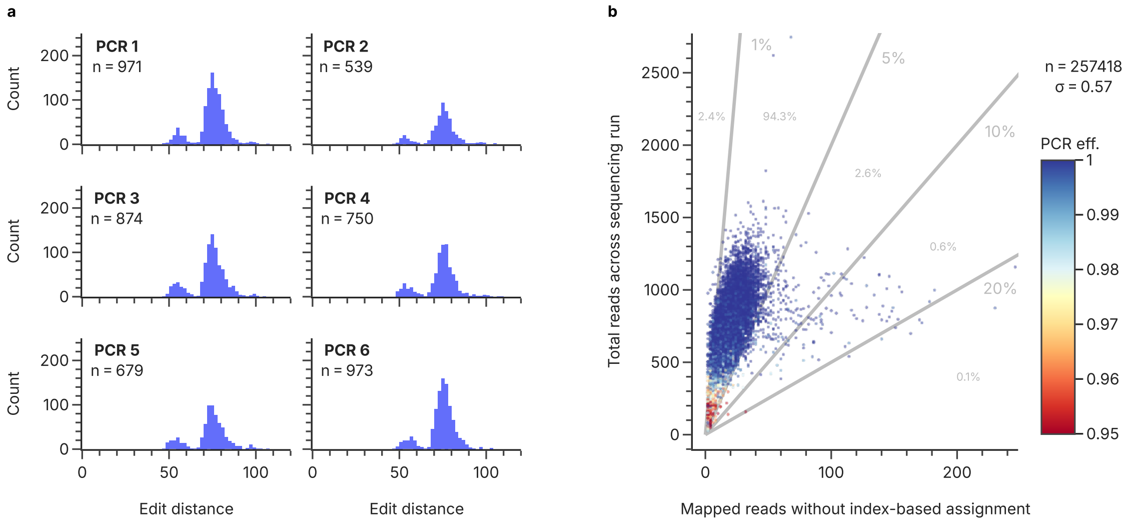


**Supplementary Fig. 38** Occurrence of sequences of the GCfix pool during demultiplexing and post-processing of the sequencing data. The distribution of minimum edit distances of all sequences removed during post-processing to the design sequences (a) shows that no real reads are lost during adapter removal and mapping. The occurrence of design sequences without experimental assignment after demultiplexing (b) shows that all sequences are found in the unassigned data at approximately the same rate, proportional to their overall presence.

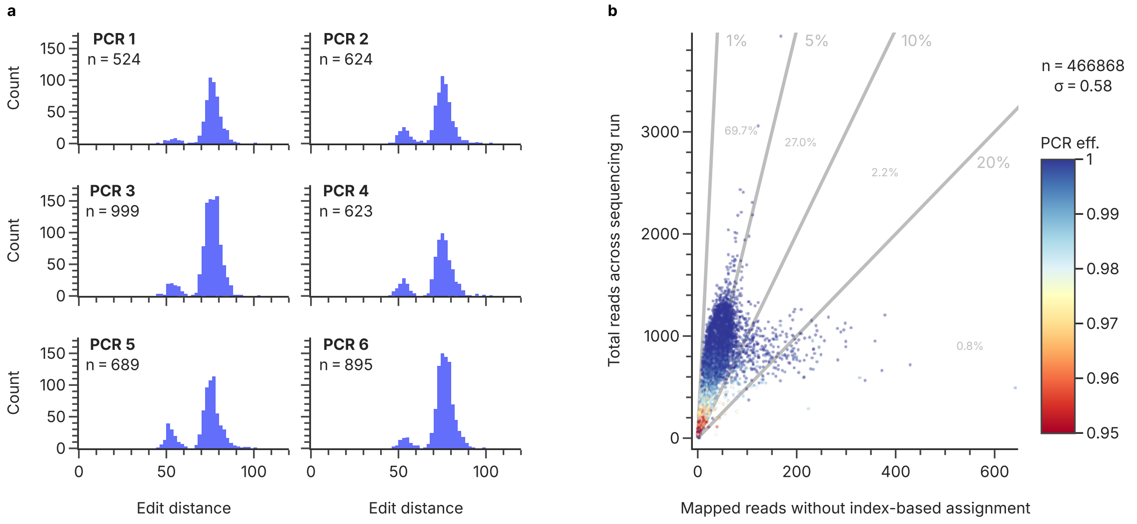


**Supplementary Fig. 39** Occurrence of sequences of the validation pool using the GCall/GCfix conditions during demultiplexing and post-processing of the sequencing data. The distribution of minimum edit distances of all sequences removed during post-processing to the design sequences (a) shows that no real reads are lost during adapter removal and mapping. The occurrence of design sequences without experimental assignment after demultiplexing (b) shows that all sequences are found in the unassigned data at approximately the same rate, proportional to their overall presence.

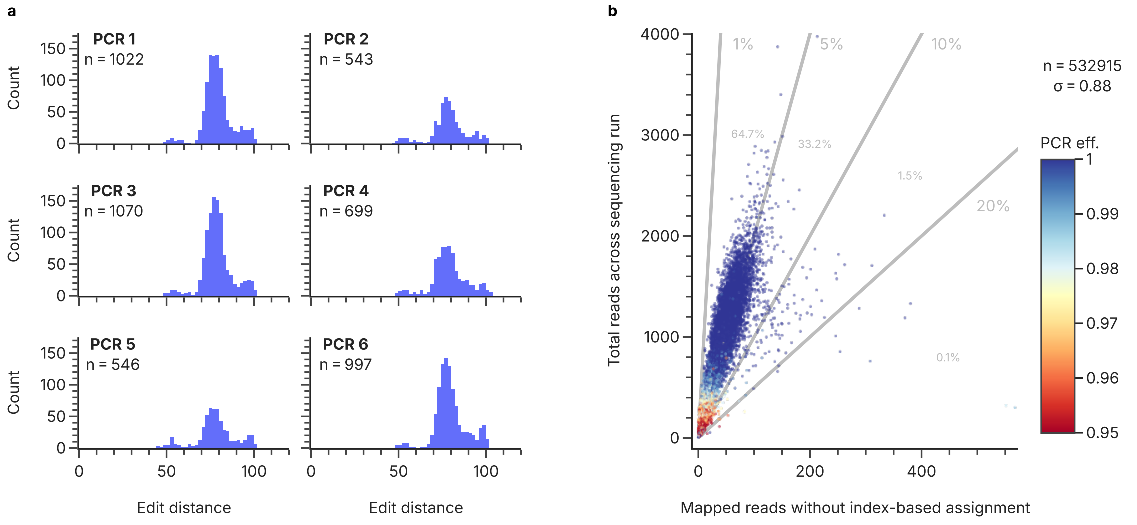


**Supplementary Fig. 40** Occurrence of sequences of the validation pool using the Erlich et al. conditions during demultiplexing and post-processing of the sequencing data. The distribution of minimum edit distances of all sequences removed during post-processing to the design sequences (a) shows that no real reads are lost during adapter removal and mapping. The occurrence of design sequences without experimental assignment after demultiplexing (b) shows that all sequences are found in the unassigned data at approximately the same rate, proportional to their overall presence.

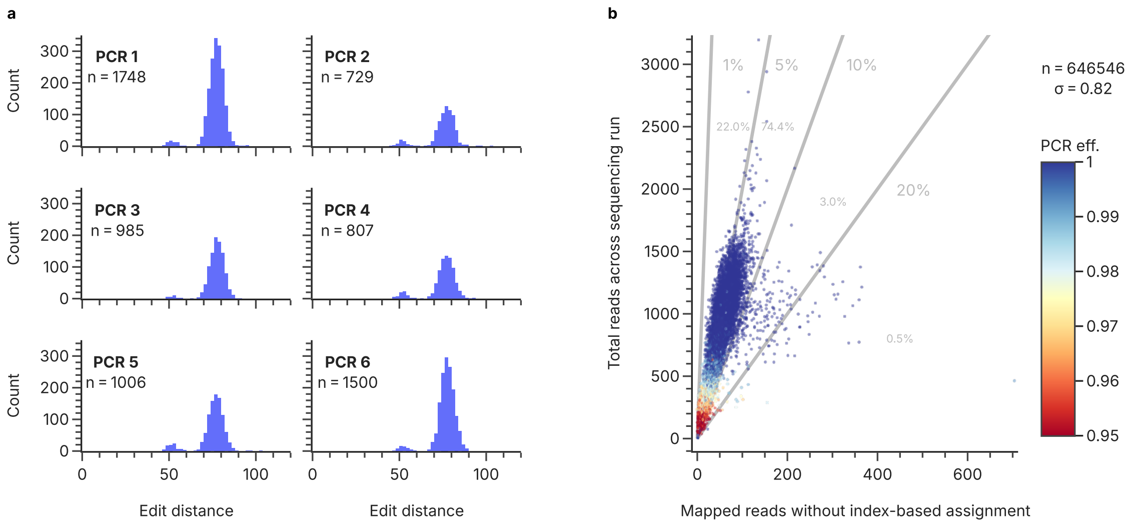


**Supplementary Fig. 41** Occurrence of sequences of the validation pool using the Erlich et al. conditions, in the internal repeat of this experiment, during demultiplexing and post-processing of the sequencing data. The distribution of minimum edit distances of all sequences removed during post-processing to the design sequences (a) shows that no real reads are lost during adapter removal and mapping. The occurrence of design sequences without experimental assignment after demultiplexing (b) shows that all sequences are found in the unassigned data at approximately the same rate, proportional to their overall presence.

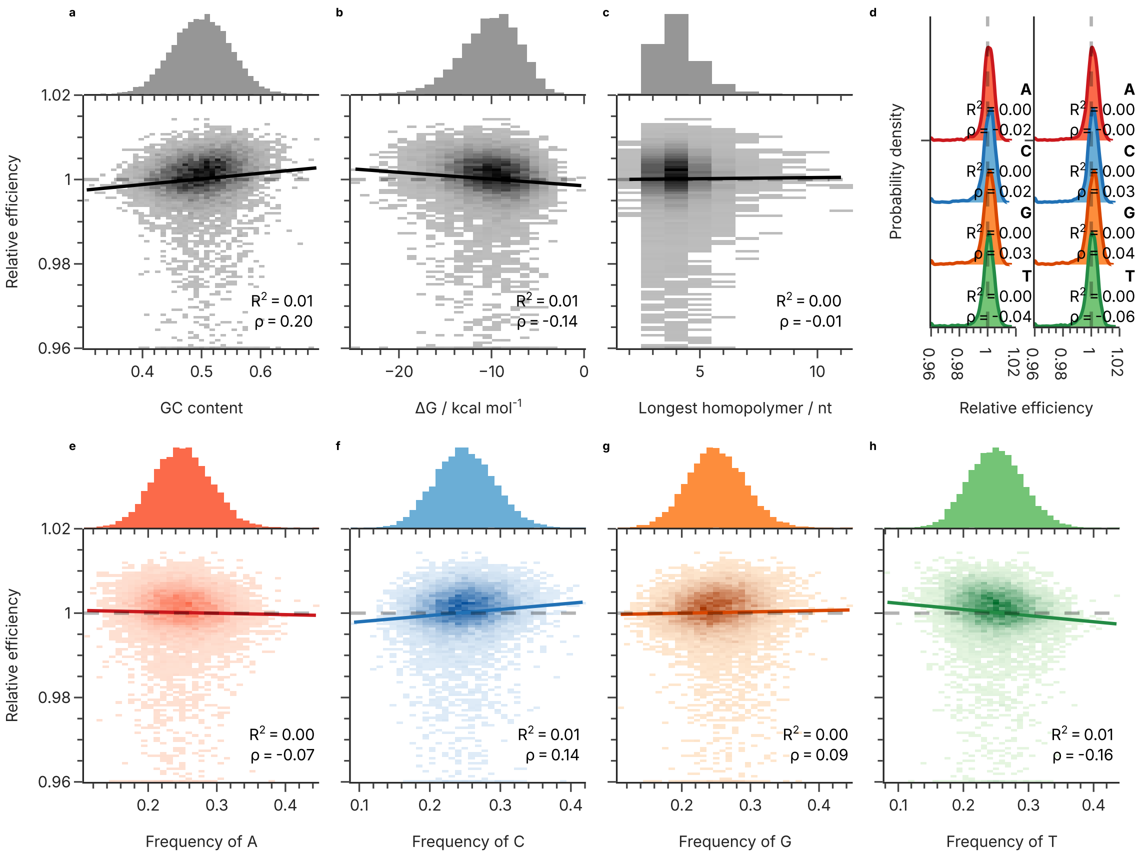


**Supplementary Fig. 42** Correlation between estimated amplification efficiency and sequence properties for the GCall dataset. The sequence properties shown are the GC content (a), the free energy calculated by mfold (b), the length of the longest homopolymer in each sequence (c), the first and last nucleotide (d), as well as the frequency of A (e), C (f), G (g), and T (h) nucleotides in each sequence. Also shown are histograms of all sequence properties (top of panels), linear regressions (solid lines), the corresponding coefficients of determination ($R^2$), as well as Spearman's rank correlation coefficients ($\rho$).

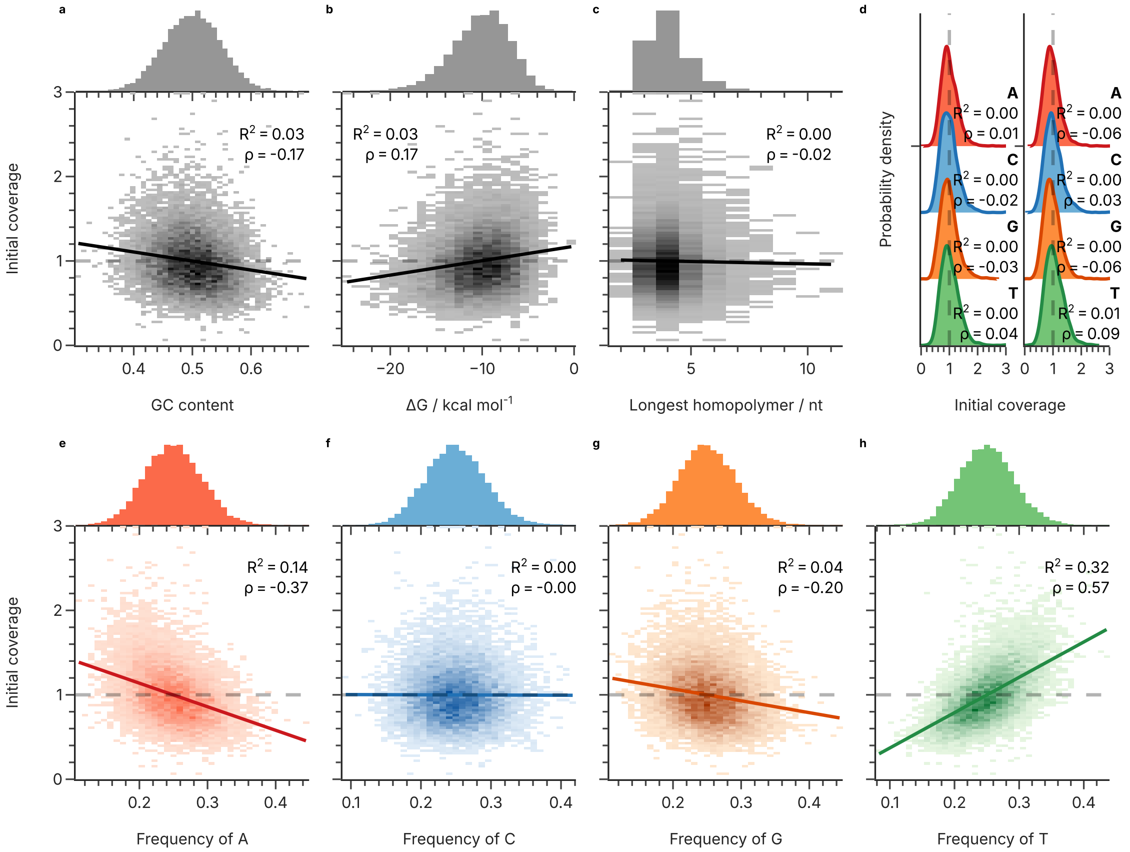


**Supplementary Fig. 43** Correlation between estimated initial abundance and sequence properties for the GCall dataset. The sequence properties shown are the GC content (a), the free energy calculated by mfold (b), the length of the longest homopolymer in each sequence (c), the first and last nucleotide (d), as well as the frequency of A (e), C (f), G (g), and T (h) nucleotides in each sequence. Also shown are histograms of all sequence properties (top of panels), linear regressions (solid lines), the corresponding coefficients of determination ($R^2$), as well as Spearman's rank correlation coefficients ($\rho$).

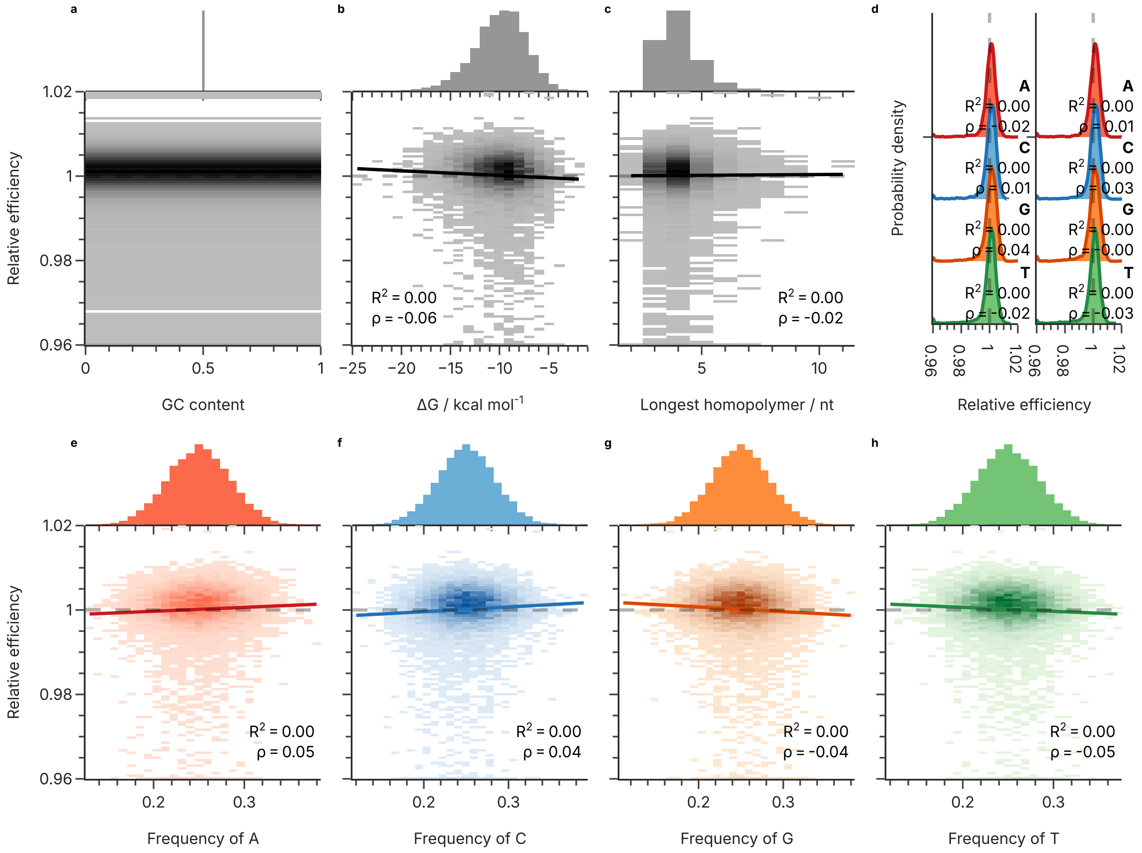


**Supplementary Fig. 44** Correlation between estimated amplification efficiency and sequence properties for the GCfix dataset. The sequence properties shown are the GC content (a), the free energy calculated by mfold (b), the length of the longest homopolymer in each sequence (c), the first and last nucleotide (d), as well as the frequency of A (e), C (f), G (g), and T (h) nucleotides in each sequence. Also shown are histograms of all sequence properties (top of panels), linear regressions (solid lines), the corresponding coefficients of determination ($R^2$), as well as Spearman's rank correlation coefficients ($\rho$).

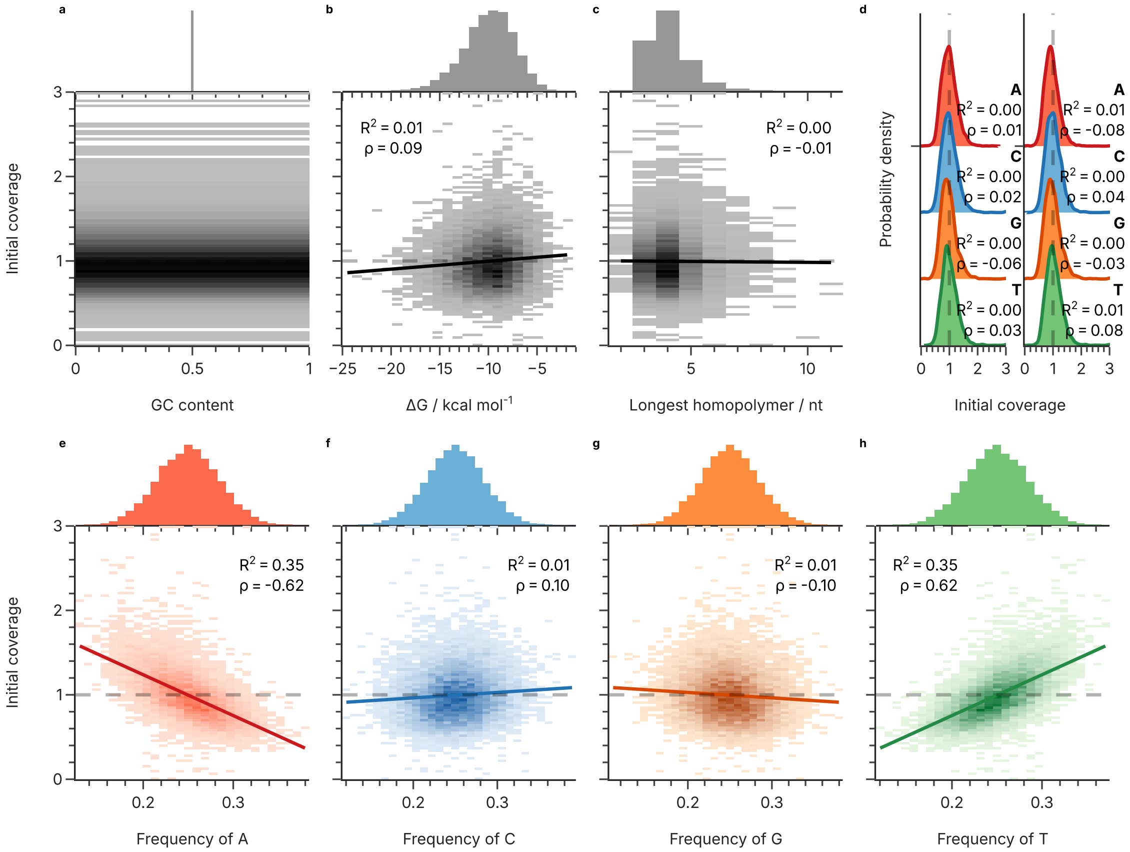


**Supplementary Fig. 45** Correlation between estimated initial abundance and sequence properties for the GCfix dataset. The sequence properties shown are the GC content (a), the free energy calculated by mfold (b), the length of the longest homopolymer in each sequence (c), the first and last nucleotide (d), as well as the frequency of A (e), C (f), G (g), and T (h) nucleotides in each sequence. Also shown are histograms of all sequence properties (top of panels), linear regressions (solid lines), the corresponding coefficients of determination ($R^2$), as well as Spearman's rank correlation coefficients ($\rho$).

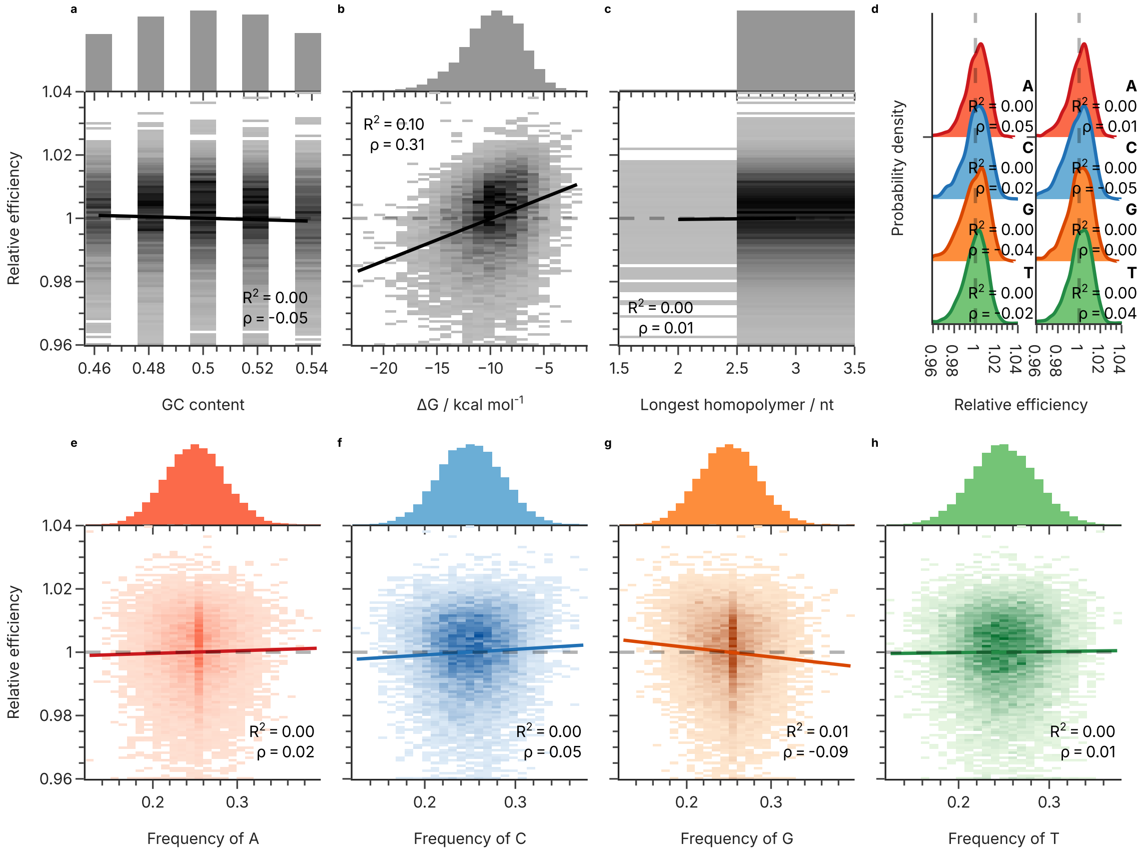


**Supplementary Fig. 46** Correlation between estimated amplification efficiency and sequence properties for the Koch et al. dataset. The sequence properties shown are the GC content (a), the free energy calculated by mfold (b), the length of the longest homopolymer in each sequence (c), the first and last nucleotide (d), as well as the frequency of A (e), C (f), G (g), and T (h) nucleotides in each sequence. Also shown are histograms of all sequence properties (top of panels), linear regressions (solid lines), the corresponding coefficients of determination ($R^2$), as well as Spearman's rank correlation coefficients ($\rho$).

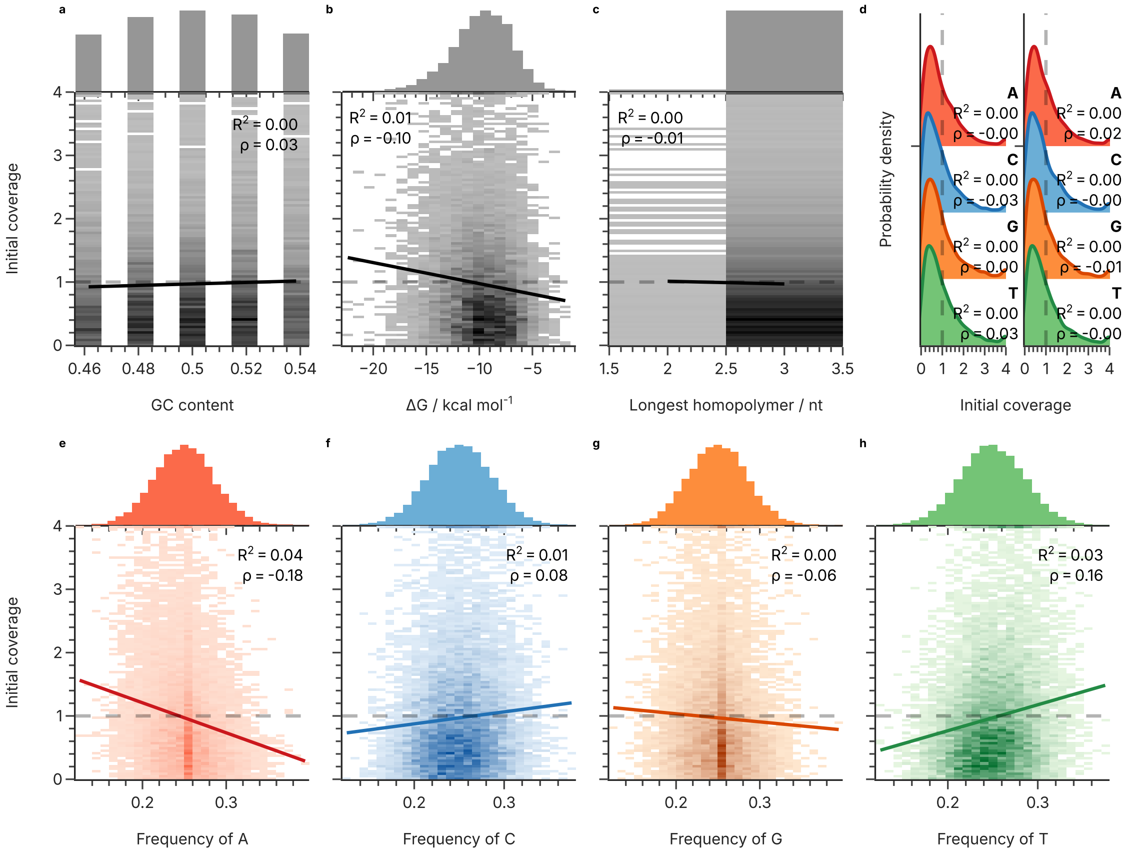


**Supplementary Fig. 47** Correlation between estimated initial abundance and sequence properties for the Koch et al. dataset. The sequence properties shown are the GC content (a), the free energy calculated by mfold (b), the length of the longest homopolymer in each sequence (c), the first and last nucleotide (d), as well as the frequency of A (e), C (f), G (g), and T (h) nucleotides in each sequence. Also shown are histograms of all sequence properties (top of panels), linear regressions (solid lines), the corresponding coefficients of determination ($R^2$), as well as Spearman's rank correlation coefficients ($\rho$).

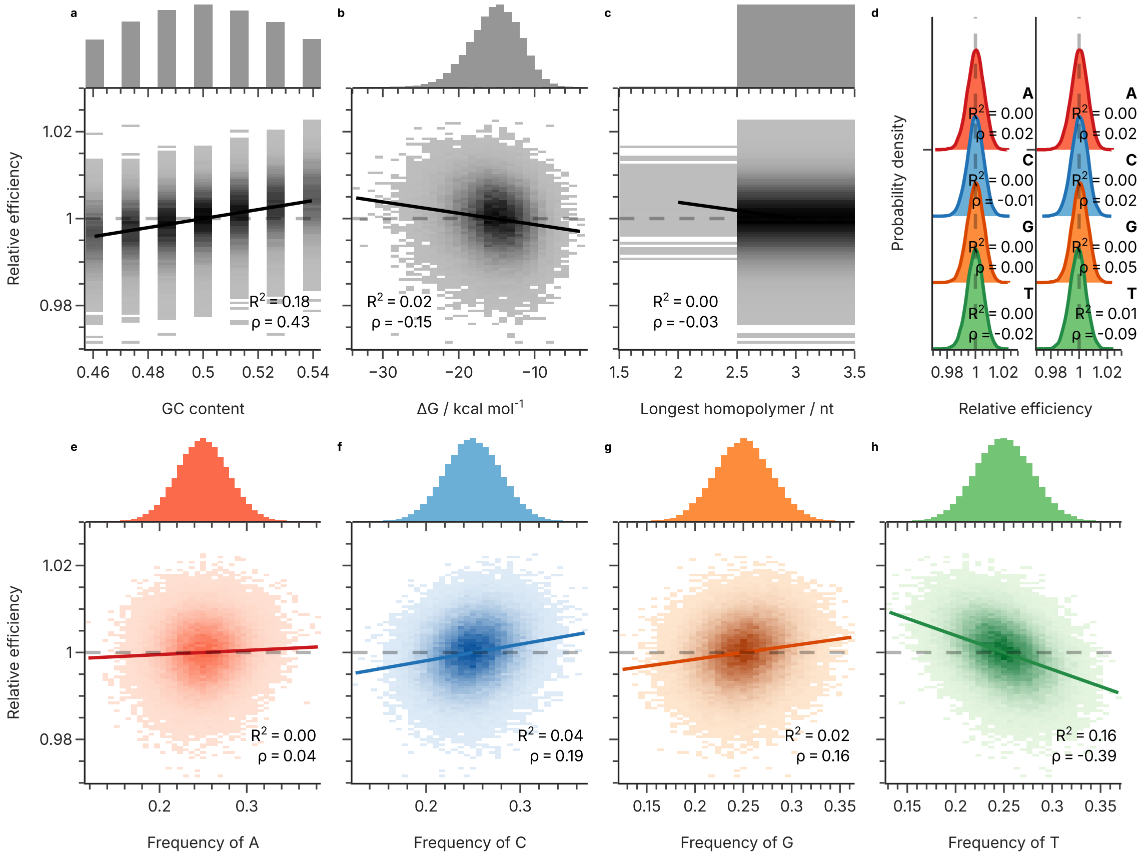


**Supplementary Fig. 48** Correlation between estimated amplification efficiency and sequence properties for the Erlich et al. dataset. The sequence properties shown are the GC content (a), the free energy calculated by mfold (b), the length of the longest homopolymer in each sequence (c), the first and last nucleotide (d), as well as the frequency of A (e), C (f), G (g), and T (h) nucleotides in each sequence. Also shown are histograms of all sequence properties (top of panels), linear regressions (solid lines), the corresponding coefficients of determination ($R^2$), as well as Spearman's rank correlation coefficients ($\rho$).

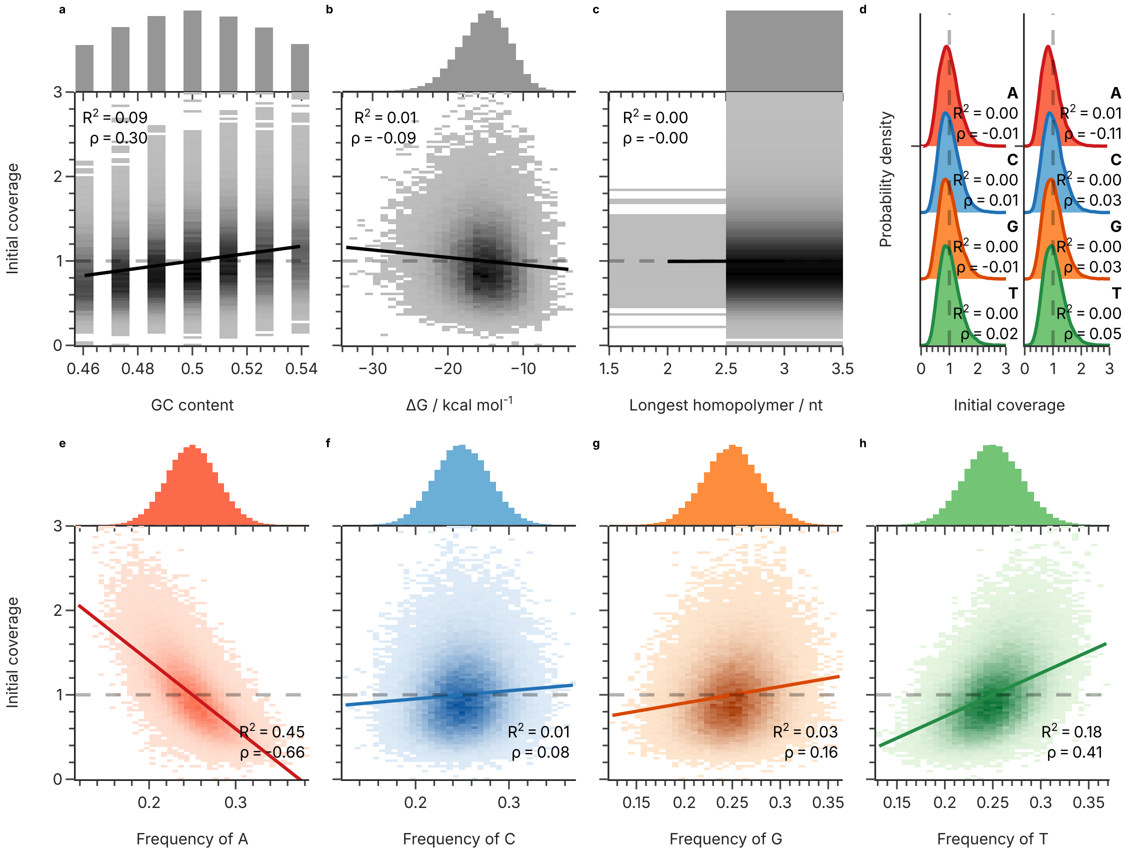


**Supplementary Fig. 49** Correlation between estimated initial abundance and sequence properties for the Erlich et al. dataset. The sequence properties shown are the GC content (a), the free energy calculated by mfold (b), the length of the longest homopolymer in each sequence (c), the first and last nucleotide (d), as well as the frequency of A (e), C (f), G (g), and T (h) nucleotides in each sequence. Also shown are histograms of all sequence properties (top of panels), linear regressions (solid lines), the corresponding coefficients of determination ($R^2$), as well as Spearman's rank correlation coefficients ($\rho$).

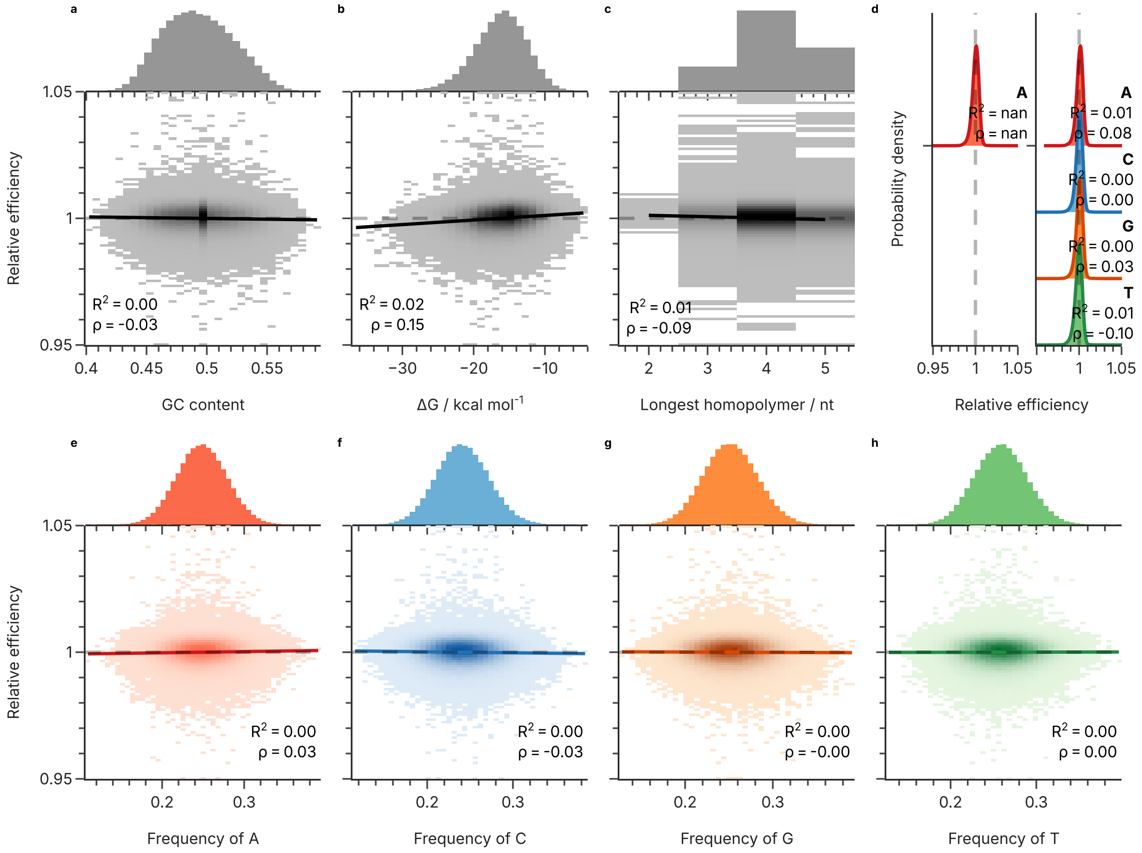


**Supplementary Fig. 50** Correlation between estimated amplification efficiency and sequence properties for the Song et al. dataset. The sequence properties shown are the GC content (a), the free energy calculated by mfold (b), the length of the longest homopolymer in each sequence (c), the first and last nucleotide (d), as well as the frequency of A (e), C (f), G (g), and T (h) nucleotides in each sequence. Also shown are histograms of all sequence properties (top of panels), linear regressions (solid lines), the corresponding coefficients of determination ($R^2$), as well as Spearman's rank correlation coefficients ($\rho$).

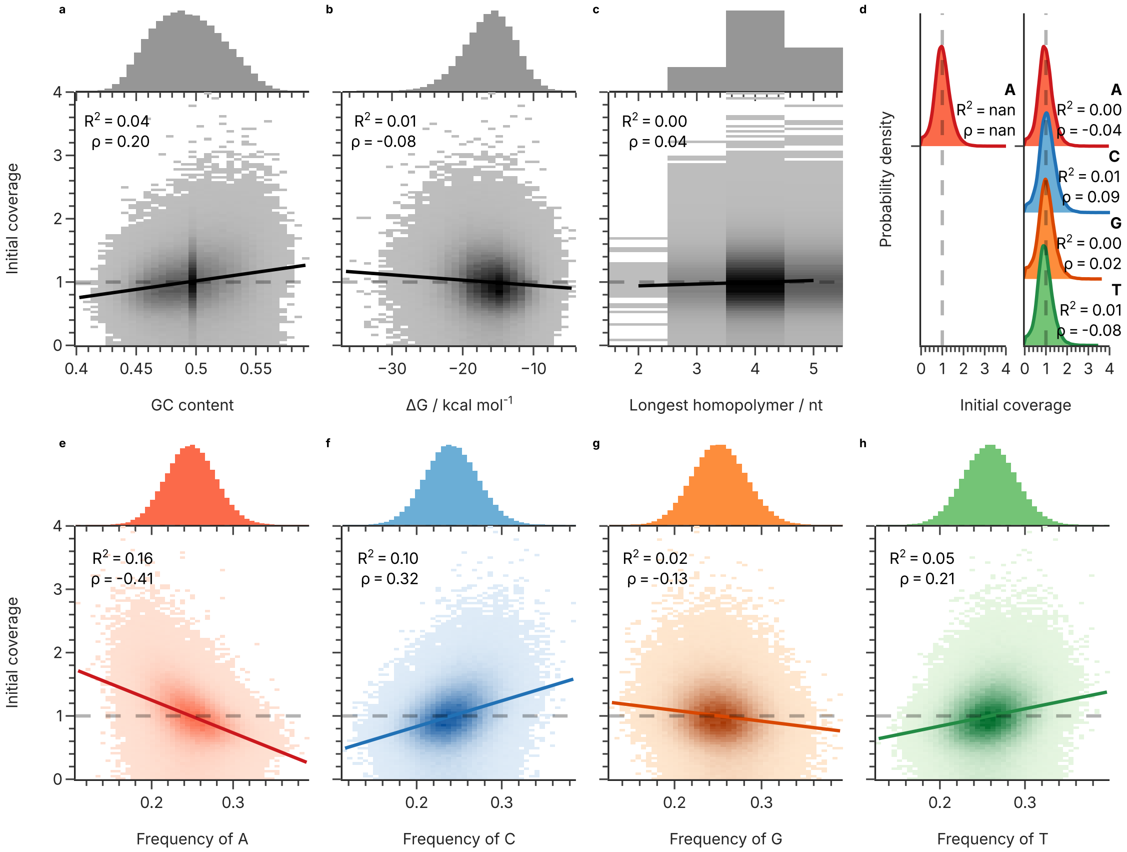


**Supplementary Fig. 51** Correlation between estimated initial abundance and sequence properties for the Song et al. dataset. The sequence properties shown are the GC content (a), the free energy calculated by mfold (b), the length of the longest homopolymer in each sequence (c), the first and last nucleotide (d), as well as the frequency of A (e), C (f), G (g), and T (h) nucleotides in each sequence. Also shown are histograms of all sequence properties (top of panels), linear regressions (solid lines), the corresponding coefficients of determination ($R^2$), as well as Spearman's rank correlation coefficients ($\rho$).

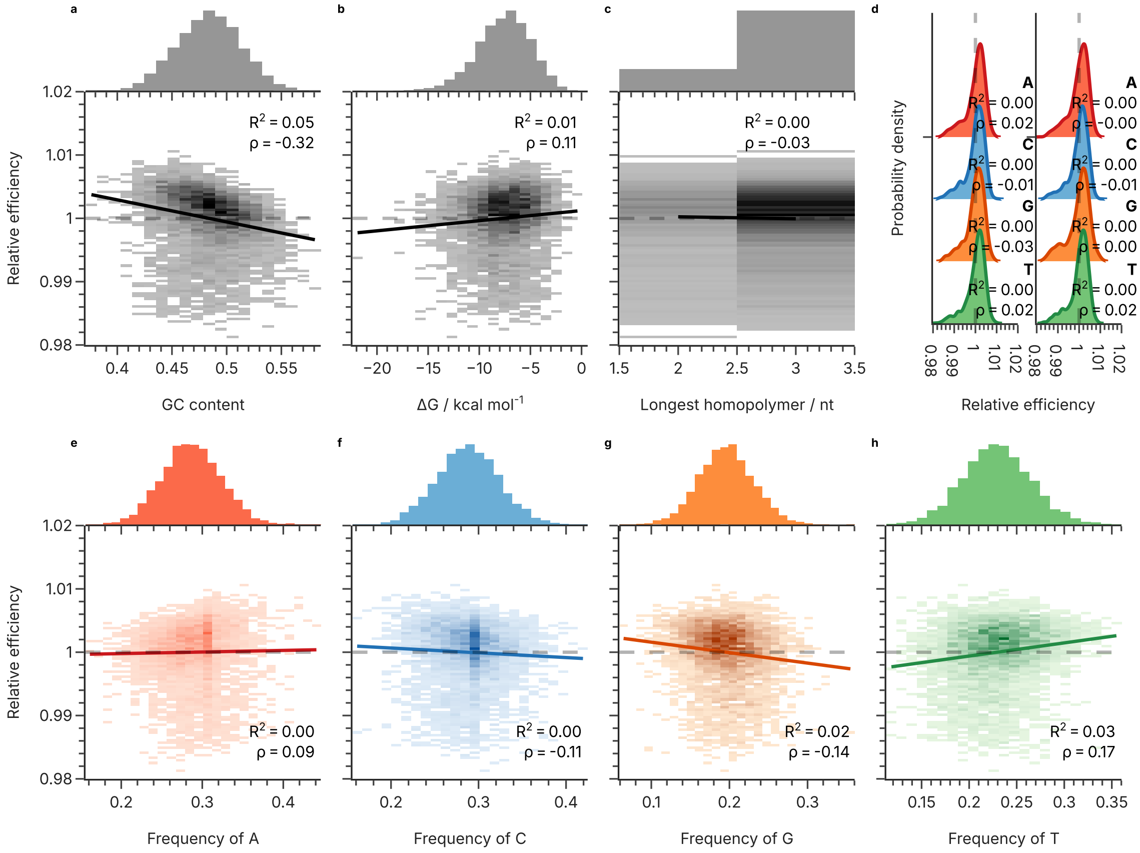


**Supplementary Fig. 52** Correlation between estimated amplification efficiency and sequence properties for the Choi et al. dataset. The sequence properties shown are the GC content (a), the free energy calculated by mfold (b), the length of the longest homopolymer in each sequence (c), the first and last nucleotide (d), as well as the frequency of A (e), C (f), G (g), and T (h) nucleotides in each sequence. Also shown are histograms of all sequence properties (top of panels), linear regressions (solid lines), the corresponding coefficients of determination ($R^2$), as well as Spearman's rank correlation coefficients ($\rho$).

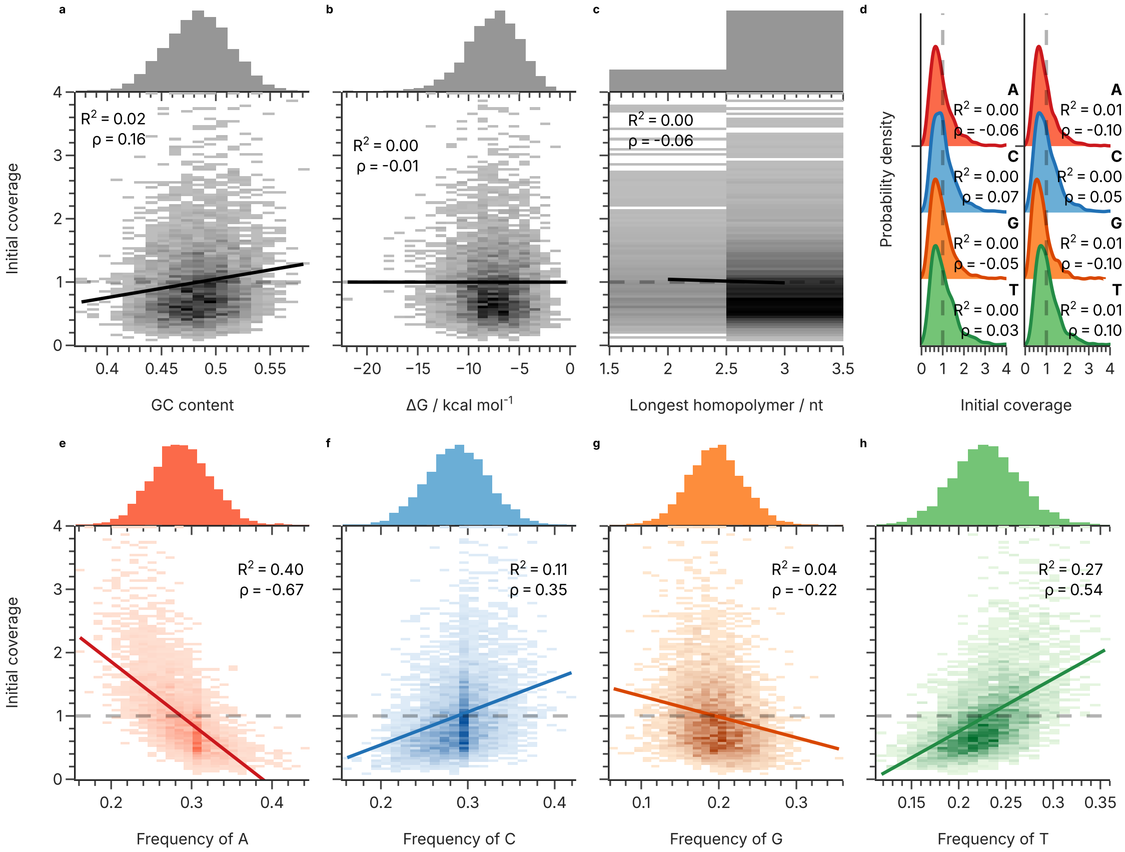


**Supplementary Fig. 53** Correlation between estimated initial abundance and sequence properties for the Choi et al. dataset. The sequence properties shown are the GC content (a), the free energy calculated by mfold (b), the length of the longest homopolymer in each sequence (c), the first and last nucleotide (d), as well as the frequency of A (e), C (f), G (g), and T (h) nucleotides in each sequence. Also shown are histograms of all sequence properties (top of panels), linear regressions (solid lines), the corresponding coefficients of determination ($R^2$), as well as Spearman's rank correlation coefficients ($\rho$).

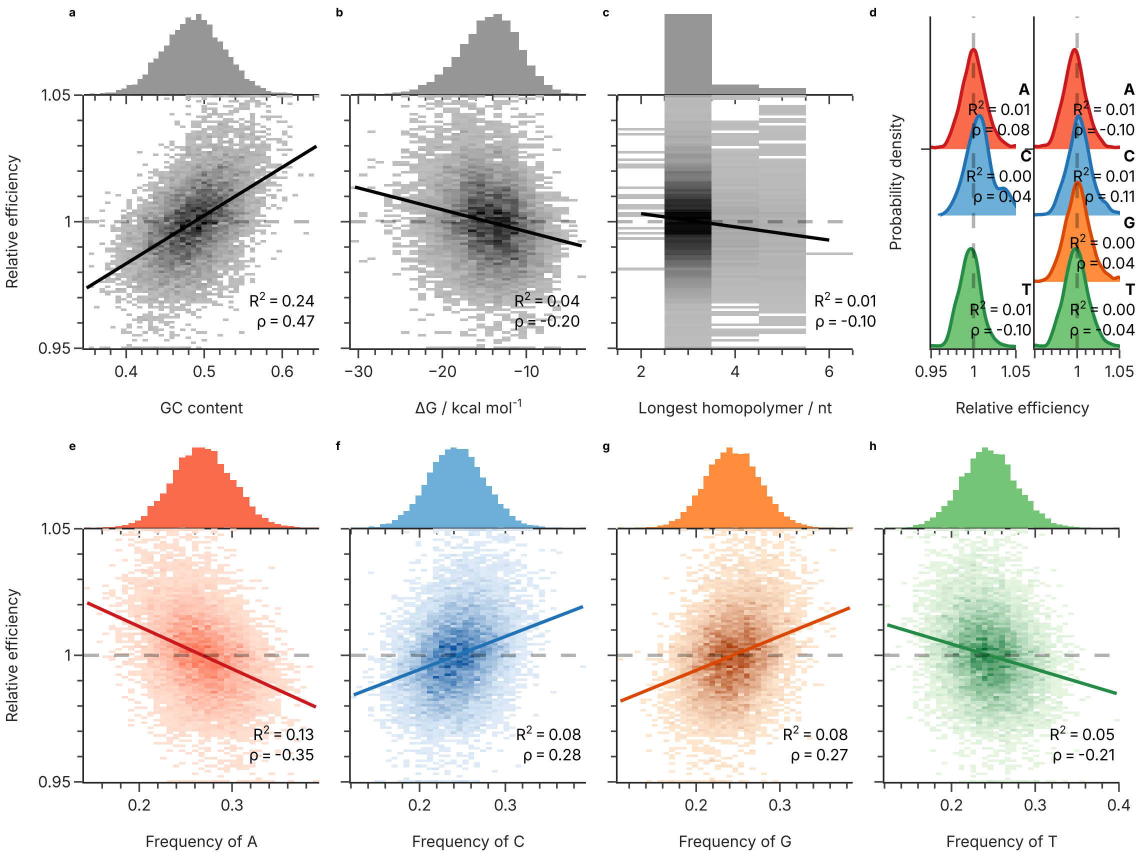


**Supplementary Fig. 54** Correlation between estimated amplification efficiency and sequence properties for the Gao et al. dataset. The sequence properties shown are the GC content (a), the free energy calculated by mfold (b), the length of the longest homopolymer in each sequence (c), the first and last nucleotide (d), as well as the frequency of A (e), C (f), G (g), and T (h) nucleotides in each sequence. Also shown are histograms of all sequence properties (top of panels), linear regressions (solid lines), the corresponding coefficients of determination ($R^2$), as well as Spearman's rank correlation coefficients ($\rho$).

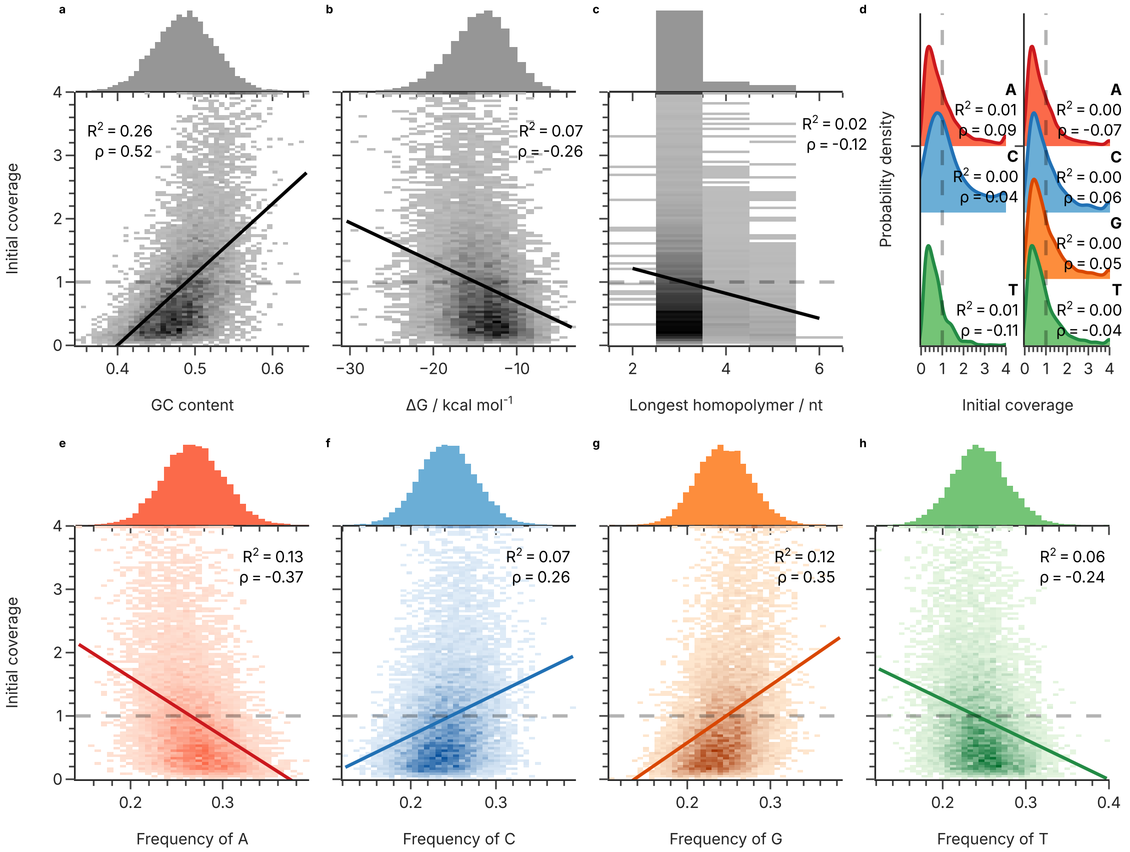


**Supplementary Fig. 55** Correlation between estimated initial abundance and sequence properties for the Gao et al. dataset. The sequence properties shown are the GC content (a), the free energy calculated by mfold (b), the length of the longest homopolymer in each sequence (c), the first and last nucleotide (d), as well as the frequency of A (e), C (f), G (g), and T (h) nucleotides in each sequence. Also shown are histograms of all sequence properties (top of panels), linear regressions (solid lines), the corresponding coefficients of determination ($R^2$), as well as Spearman's rank correlation coefficients ($\rho$).

### Supplementary Tables

| Hyperparameter | Search values |
|---|---|
| Model | RNN, LSTM, GRU |
| Number of layers | 1, 2, 3 |
| Embedding dimension | 32, 64, 128 |
| Hidden dimension | 32, 64, 128 |
| Learning rate | $10^{-3}$, $10^{-4}$, $10^{-5}$ |
| Batch size | 64, 128, 256 |
| Weight decay | 0, $10^{-2}$, $10^{-3}$, $10^{-4}$ |

**Supplementary Table 1** Hyperparameter grid and ranges for the hyperparameter search of the RNN-based model.

**Supplementary Table 2** Overview of qPCR results.

| | | GCall | #11493 | #00006 | #09807 | #01634 |
|---|---|---|---|---|---|---|
| | Category | reference | low $\epsilon$ | medium $\epsilon$ | low $\epsilon$ | high $\epsilon$ |
| Exp. 1 | Slope | 3.52 | 3.62 | 3.59 | 4.49 | 3.50 |
| | Intercept | 12.5 | 6.40 | 4.01 | 6.31 | 4.35 |
| | $R^2$ | >0.999 | 0.999 | >0.999 | 0.994 | >0.999 |
| | $\epsilon$ | 92.2% | 88.8% | 89.8% | 67.0% | 92.9% |
| Exp. 2 | Slope | 3.50 | 3.61 | 3.56 | 4.44 | 3.55 |
| | Intercept | 12.6 | 6.33 | 4.46 | 6.29 | 4.20 |
| | $R^2$ | 0.999 | >0.999 | >0.999 | 0.991 | >0.999 |
| | $\epsilon$ | 93.0% | 89.2% | 91.0% | 68.0% | 91.3% |
| Exp. 3 | Slope | 3.42 | 3.60 | 3.54 | 4.41 | 3.49 |
| | Intercept | 12.7 | 6.34 | 4.07 | 6.24 | 4.33 |
| | $R^2$ | >0.999 | >0.999 | >0.999 | 0.993 | >0.999 |
| | $\epsilon$ | 96.0% | 89.6% | 91.8% | 68.6% | 93.3% |
| Overall | $\bar{\epsilon}$ | 93.7% | 89.2% | 90.8% | 67.8% | 92.5% |
| | SEM | 1.14% | 0.24% | 0.58% | 0.48% | 0.61% |

**Supplementary Table 3** Multiple comparisons of the qPCR results using Tukey's range test.

| Sample 1 | Sample 2 | $\Delta\epsilon$ | p-value | Confidence interval | |
|---|---|---|---|---|---|
| #00006 | #01634 | 1.67% | 0.4521 | -1.48% | 4.82% |
| #00006 | #09807 | -23.00% | $3 \times 10^{-9}$ | -26.14% | -19.85% |
| #00006 | #11493 | -1.64% | 0.4671 | -4.79% | 1.51% |
| #00006 | GCall | 2.85% | 0.0812 | -0.3% | 6.00% |
| #01634 | #09807 | -24.67% | $1 \times 10^{-9}$ | -27.81% | -21.52% |
| #01634 | #11493 | -3.31% | 0.0385 | -6.46% | -0.16% |
| #01634 | GCall | 1.18% | 0.7341 | -1.97% | 4.33% |
| #09807 | #11493 | 21.36% | $6 \times 10^{-9}$ | 18.21% | 24.50% |
| #09807 | GCall | 25.84% | $9 \times 10^{-10}$ | 22.7% | 28.99% |
| #11493 | GCall | 4.49% | 0.0059 | 1.34% | 7.64% |

**Supplementary Table 4** This table presents the positive class prevalence of each dataset for different statistically defined thresholds to categorize low-efficiency and normal-efficiency DNA sequences.

| **Dataset** | Choi_et_al | Erlich_et_al | GCall | GCfix | Gao_et_al | Koch_et_al | Song_et_al |
|---|---|---|---|---|---|---|---|
| 1 $\sigma$ | 0.146905 | 0.156250 | 0.059843 | 0.051192 | 0.141678 | 0.151776 | 0.020000 |
| 2 $\sigma$ | 0.059601 | 0.025861 | 0.027171 | 0.024679 | 0.016688 | 0.038278 | 0.034714 |
| 3 $\sigma$ | 0.011962 | 0.002556 | 0.016086 | 0.015258 | 0.003651 | 0.007020 | 0.008859 |

**Supplementary Table 5** Spearman rank correlations of the estimated parameters of the three external validation datasets.

| | Initial abundance | | | PCR efficiency | | |
|---|---|---|---|---|---|---|
| Group | GCall/fix | Erlich (ext.) | Erlich (int.) | GCall/fix | Erlich (ext.) | Erlich (int.) |
| GCall/fix | 1.00 | 0.30 | 0.28 | 1.00 | -0.23 | -0.22 |
| Erlich (ext.) | 0.30 | 1.00 | 0.63 | -0.23 | 1.00 | 0.91 |
| Erlich (int.) | 0.28 | 0.63 | 1.00 | -0.22 | 0.91 | 1.00 |

**Supplementary Table 6** Sequences used in the study.

| Name | Description | Sequence (5'-3') |
|---|---|---|
| 0F | Forward primer | ACACGACGCTCTTCCGATCT |
| 0R | Reverse primer | AGACGTGTGCTCTTCCGATCT |
| 0F_4W | Degenerate forward primer | WWWWACACGACGCTCTTCCGATCT |
| 0R_4W | Degenerate reverse primer | WWWWAGACGTGTGCTCTTCCGATCT |
| 2FUF | Forward sequencing primer | AATGATACGGCGACCACCGAGATCTACACTCTTTCCCTACA CGACGCTCTTCCGATCT |
| 2RIF-GM5 | Indexed reverse sequencing primer | CAAGCAGAAGACGGCATACGAGATCACTGTGTGACTGGAGT TCAGACGTGTGCTCTTCCGATCT |
| 2RIF-GM7 | Indexed reverse sequencing primer | CAAGCAGAAGACGGCATACGAGATGATCTGGTGACTGGAGT TCAGACGTGTGCTCTTCCGATCT |
| 2RIF-GM8 | Indexed reverse sequencing primer | CAAGCAGAAGACGGCATACGAGATTCAAGTGTGACTGGAGT TCAGACGTGTGCTCTTCCGATCT |
| 2RIF-GM10 | Indexed reverse sequencing primer | CAAGCAGAAGACGGCATACGAGATAAGCTAGTGACTGGAGT TCAGACGTGTGCTCTTCCGATCT |
| 2RIF-GM11 | Indexed reverse sequencing primer | CAAGCAGAAGACGGCATACGAGATGTAGCCGTGACTGGAGT TCAGACGTGTGCTCTTCCGATCT |
| 2RIF-GM12 | Indexed reverse sequencing primer | CAAGCAGAAGACGGCATACGAGATTACAAGGTGACTGGAGT TCAGACGTGTGCTCTTCCGATCT |
| 2RIF-GM17 | Indexed reverse sequencing primer | CAAGCAGAAGACGGCATACGAGATCTCTACGTGACTGGAGT TCAGACGTGTGCTCTTCCGATCT |
| #11493 | qPCR test sequence | ACACGACGCTCTTCCGATCTCGTGTATAGGCTGACTGTTAT GTTCGTGCAGCACGCTGCATGGCTGATCGTATGTACTTGGG CACTCTAAAGATTCAAGGCTAAGGAAGAAGGATAACGCTAC CCAACAGATCGGAAGAGCACACGTCT |
| #00006 | qPCR test sequence | ACACGACGCTCTTCCGATCTTATGTACTCTTGCACGCTTGG CGGGTGTTAAACTTCTGGTTCCGACCTAGCGCTCGTGCCTT CCGGTGAATGTAGGTTGCCGAATGACACATCCATGCCCTAT TCAGCAGATCGGAAGAGCACACGTCT |
| #09807 | qPCR test sequence | ACACGACGCTCTTCCGATCTGAGTTCAAGCGCCGTGTAGCC TGATCTGGTCCTTATACTAGCCTGTTCACAAAGATATAGAT CTGTATAAACAGCGCCATTCAAGCTGAGAGAGAGGGCCCAG GGGCCAGATCGGAAGAGCACACGTCT |
| #01634 | qPCR test sequence | ACACGACGCTCTTCCGATCTTTGCTTTGCTGTTGTGCGGTA CACCCCAACGTGATCTTTAGTCCTTGGAAAGTCCACCACAT CTCTTGTAGGGCTAGCTTATTCTAGAACAACAGCAAGGGCT GCCACAGATCGGAAGAGCACACGTCT |

**Supplementary Table 7** Overview of datasets.

| Dataset | GCall | GCfix | Koch et al. | Erlich et al. | Song et al. | Choi et al. | Gao et al. |
|---|---|---|---|---|---|---|---|
| Source | PRJEB65931 | PRJEB65931 | PRJEB35217 | PRJEB19305<br>PRJEB19307 | 10.6084/m9.figshare.16727122<br>10.6084/m9.figshare.17193128<br>10.6084/m9.figshare.18515045 | PRJNA555140 | pers. comm. |
| **Experimental design** | | | | | | | |
| #Experiments | 6 | 6 | 7 | 2 | 6 | 2 | 3 |
| Cycle counts | 15-90 | 15-90 | 44-119 | 10, 100 | 30-180 | 17, 340 | 10, 50, 100 |
| Synthesis provider | Twist | Twist | CustomArray | Twist | Twist | CustomArray | Twist |
| Polymerase | KAPA FAST | KAPA FAST | KAPA FAST | Q5 HiFi | error-prone polymerase | KAPA HiFi | Q5 HiFi |
| **Pool design** | | | | | | | |
| #Sequences | 12000 | 12000 | 12000 | 72000 | 210000 | 5173 | 11520 |
| Seq. length | 108 | 108 | 104 | 152 | 164 | 93 | 146 |
| Seq. constraints | - | GC | GC, HP | GC, HP | GC, HP | - | GC, motifs |
| Seq. randomized | Yes | Yes | No | No | No | No | No |
| **Pre-processing** | | | | | | | |
| #Seq. removed | 2 | 6 | 35 | 0 | 53 | 408 | 15 |
| %Seq. removed | 0.017% | 0.05% | 0.29% | 0% | 0.025% | 7.9% | 0.13% |
| **Parameter fit** | | | | | | | |
| $R^2_{\text{baseline}}$ | 0.750 | 0.758 | 0.868 | 0.755 | 0.424 | 0.684 | 0.434 |
| $R^2_{\text{model}}$ | 0.859 | 0.868 | 0.957 | 1.000 | 0.967 | 1.000 | 0.911 |
| $df_1$ | 11998 | 11994 | 11965 | 72000 | 209947 | 4765 | 11505 |
| $df_2$ | 47992 | 47976 | 59825 | 0 | 839788 | 0 | 11505 |
| $F(df_1, df_2)$ | 3.08 | 3.33 | 10.2 | - | 65.8 | - | 5.38 |
